# Identification of the keystone species in non-alcoholic fatty liver disease by causal inference and dynamic intervention modeling

**DOI:** 10.1101/2020.08.06.240655

**Authors:** Dingfeng Wu, Na Jiao, Ruixin Zhu, Yida Zhang, Wenxing Gao, Sa Fang, Yichen Li, Sijing Cheng, Chuan Tian, Ping Lan, Rohit Loomba, Lixin Zhu

## Abstract

**Objective:** Keystone species are required for the integrity and stability of an ecological community, and therefore, are potential intervention targets for microbiome related diseases.

**Design:** Here we describe an algorithm for the identification of keystone species from cross-sectional microbiome data of non-alcoholic fatty liver disease (NAFLD) based on causal inference theories and dynamic intervention modeling (DIM).

**Results:** Eight keystone species in the gut of NAFLD, represented by *P. loveana*, *A. indistinctus* and *D. pneumosintes*, were identified by our algorithm, which could efficiently restore the microbial composition of the NAFLD toward a normal gut microbiome with 92.3% recovery. These keystone species regulate intestinal amino acids metabolism and acid-base environment to promote the growth of the butyrate-producing Lachnospiraceae and Ruminococcaceae species.

**Conclusion:** Our method may benefit microbiome studies in the broad fields of medicine, environmental science and microbiology.

**SUMMARY:** **What is already known about this subject?**

- Non-alcoholic fatty liver disease (NAFLD) is a complex multifactorial disease whose pathogenesis remains unclear.
- Dysbiosis in the gut microbiota affects the initiation and development of NAFLD, but the mechanisms is yet to be established.
- Keystone species represent excellent candidate targets for gut microbiome-based interventions, as they are defined as the species required for the integrity and stability of the ecological system.

**What are the new findings?**

- NAFLD showed significant dysbiosis in butyrate-producing Lachnospiraceae and Ruminococcaceae species.
- Microbial interaction networks were constructed by the novel algorithm with causal inference.
- Keystone species were identified form microbial interaction networks through dynamic intervention modeling based on generalized Lotka-Volterra model.
- Eight keystone species of NAFLD with the highest potential for restoring the microbial composition were identified.

**How might it impact on clinical practice in the foreseeable future?**

- An algorithm for the identification of keystone species from cross-sectional microbiome data based on causal inference theories and dynamic intervention modeling.
- Eight keystone species in the gut of NAFLD, represented by *P. loveana*, *A. indistinctus* and *D. pneumosintes*, which could efficiently restore the microbial composition of the NAFLD toward a normal gut microbiome.
- Our method may benefit microbiome studies in the broad fields of medicine, environmental science and microbiology.

## INTRODUCTION

Non-alcoholic fatty liver disease (NAFLD) is a complex multifactorial disease whose pathogenesis remains unclear. The advanced form of NAFLD with hepatic inflammation is non-alcoholic steatohepatitis (NASH). Studies have shown that dysbiosis in the gut microbiota affects the initiation and development of NAFLD.^1–3^ With the whole genome sequencing data of the gut microbiome, we have identified 37 microbial markers that can distinguish between mild and severe NAFLD patients.^4^ In addition, we and others observed that microbe-derived metabolites, including endogenous ethanol,^5^ bile acids^6^ and amino acids,^7^ contribute to the pathogenesis of NAFLD. Accordingly, many microbial intervention therapies targeting the gut microbiota, such as prebiotics, probiotics, and fecal microbiota transplantation (FMT) are being considered to treat NAFLD.^8^ However, these microbial interventions have not achieved satisfactory effect,^9, 10^ which may be partly explained by the structural complexity of the ecosystem we have in our gut. Therefore, further studies of the dynamic changes in the microbiome during disease development and the mechanisms behind the changes are essential for the identification of microbial targets in the precise treatment of NAFLD and other diseases related to the gut microbiome.

Keystone species represent excellent candidate targets for gut microbiome-based interventions,^11^ as they are defined as the species required for the integrity and stability of the ecological system. The alteration of the keystone species could affect much of the entire community through the interactions among the members of the ecosystem.^12^ Culture dependent studies on keystone species have been conducted in several diseases, such as periodontal disease^13^ and *Clostridium difficile* infection.^14^ The potential imperfection with the *in vitro* approaches in these studies is that most microbes could not be isolated or cultured, therefore, these methods poorly recapitulate the actual micro-environments.

The next generation sequencing techniques allowed the study of the keystone species without culture, such as the approach based on the co-occurrence network analysis.^15^ The disadvantages of the co-occurrence network are that correlation does not imply causation,^11, 15^ and that co-occurrence network does not support dynamic simulation.^16, 17^ These limitations can be addressed by causal inference analysis. The causal inference methods using time-series data are the ideal approachhes for the reconstruction of the microbial interaction networks.^18, 19^ However, due to the requirement for large sample size and frequent sampling time points, these methods are not practically applicable. By contrast, cross-sectional data is the most common data type in biomedical research. Xiao et al. pioneered the causal inference method for the study of microbial interactions with cross-sectional data. The method they developed, zero/sign-pattern inference of Jacobian matrix, requires independent steady-state data,^20^ which is hardly available from biomedical studies.

Here, for the first time, we implemented the causal inference theories by Robins^21^ and Pearl,^22^ for the construction of microbial interaction networks with cross-sectional microbiome dataset. Subsequently, the keystone species of NAFLD were identified by dynamic intervention modeling (DIM) with the interaction networks. Our algorithm was validated with an independent NAFLD cohort.

## MATERIALS AND METHODS

### Study cohorts

This study was approved by the Institutional Review Board of Tongji University, the Children and Youth Institutional Review Board (CYIRB) of the State University of New York at Buffalo and the UCSD Institutional Review Board. Enrollment of the discovery cohort from New York and the validating cohort from California were described previously.^5, 23^ Briefly, for the discovery cohort, fecal samples of adolescents collected from 22 biopsy-proven patients with NASH, 25 obese patients and 16 healthy controls (table S1) were pyrosequenced on a 454-FLX-Titanium Genome Sequencer (Roche 454 Life Sciences, Branford, Connecticut, USA). Our validating cohort consists of 31 healthy controls, 14 NAFLD patients without advanced fibrosis and 24 NAFLD patients with cirrhosis (table S1) and were sequenced with Illumina MiSeq. Patients were not involved in the design, or conduct, or reporting, or dissemination plans of in this research.

Operational taxonomic units (OTUs) were *de novo* clustered at 97% sequence identity and chimeras removed with Qiime2 after quality control of the 16S sequencing data.^24^ Taxonomy classification was assigned using classify-sklearn based on a Naive Bayes classifier against the Silva-132-99 reference sequences. OTUs that cannot be precisely annotated to any species were reassigned to species with the most similar sequences in the same genus (or family) by NCBI Blast with the default setting. And then, OTUs unassigned at class level were removed for further analysis. 97 and 190 OTUs were obtained with the sample coverage ≥ 0.5 from the discovery and validation cohorts, respectively.

### Differential abundance analysis and evaluation of the sample discrimination ability of microbes

The Wilcoxon rank-sum test was performed to identify microbes with differential abundance between study groups and the Benjamini-Hochberg was applied to control the false positive rate (FDR) in multiple comparisons (FDR ≤ 0.05). These differential microbes were then individually evaluated for their ability in sample discrimination using area under the receiver operating characteristic curve (AUC).

### Keystone species identification

The pipeline mainly consists of two steps (figure 1): 1) a novel causal inference-based method was implemented to construct the microbial causal interaction networks; and 2) keystone species were identified with dynamic intervention modeling (DIM) based on microbial interactions.

**Figure 1.**
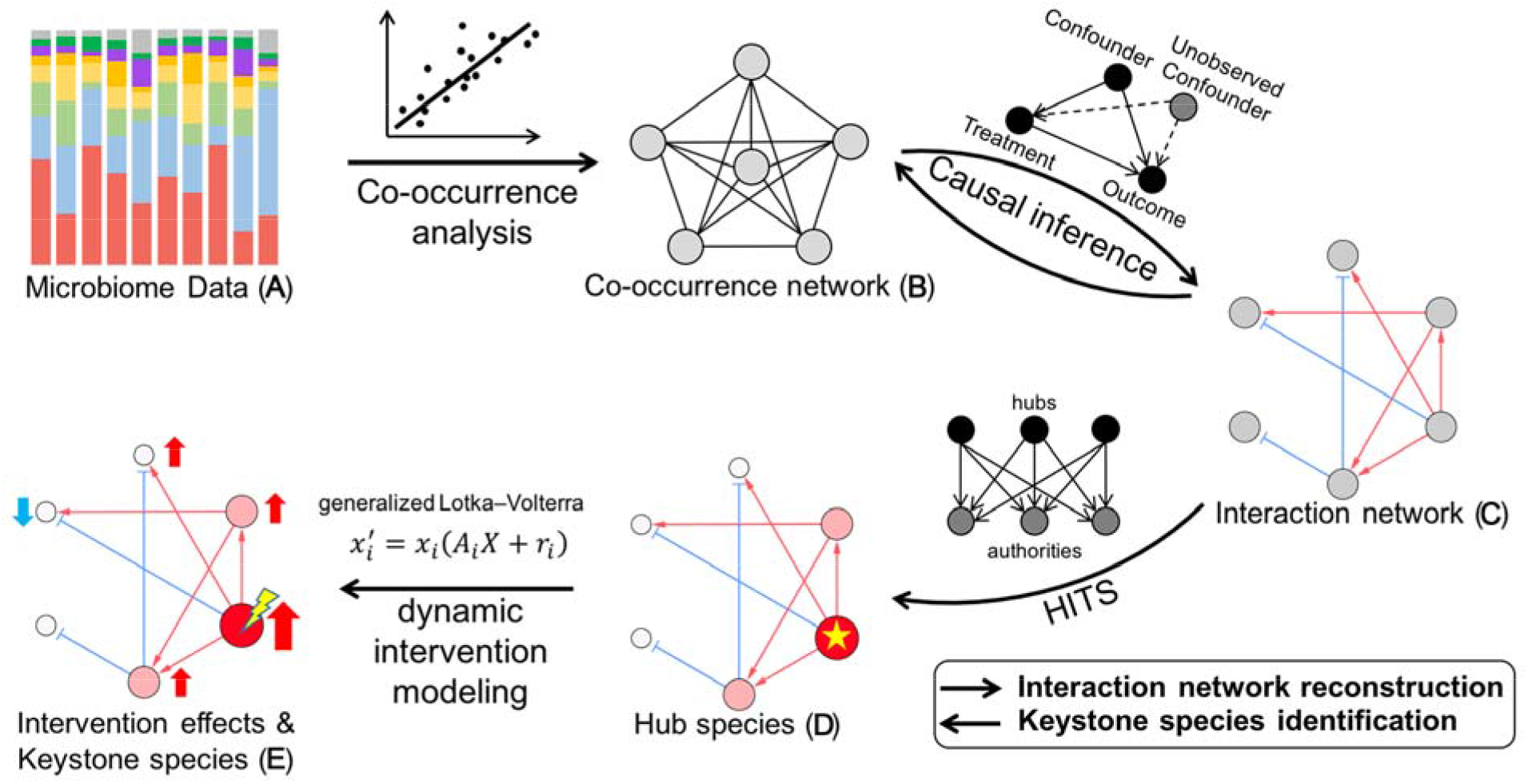
The strategy for gut microbial interaction network construction and keystone species identification. Based on the abundance data (A), co-occurrence network (B) was constructed as a priori knowledge by co-occurrence analysis; With the abundance data (A) and the microbial co-occurrence network (B), causal inference was then conducted to construct the microbial interaction network (C); Hub species (D) were identified from the microbial interaction network (C) by HITS; Finally, based on microbial interactions (C) and topological importance (D), intervention effects of microbes were evaluated, and keystone species (E) with the maximum intervention effect on the entire ecosystem were identified.

#### 1) Interaction network construction by causal inference

Causal inference needs a priori knowledge. Considering the compositional characteristics of microbial sequencing data, we used SparCC to calculate the co-occurrences of the microbial species in the gut microbiome.^25^ SparCC avoids the inaccuracies that usually occur with relative abundances in correlation analysis. Only relationships with p≤0.01 (permutation test with 1000 permutations) were included in the priori network.

Causal inference was performed by integrating two mainstream causal inference frameworks, Robins’ potential outcome (counterfactual) model^21^ and Pearl’s graphical model.^22^ Our approach effectively controls the influence of confounders and infer causality more accurately on microbial cross-sectional cohort. We extended the application of causal inference to microbiome analysis with generalized Lotka-Volterra (gLV) model, a well-known dynamic model for microbiome study.^26^ Meanwhile, iterative optimization strategy was used to improve the reliability of priori network and the accuracy of the inferred causal relations. Permutation test was performed to determine significant causalities (p ≤0.01) (Supplementary Notes).

#### 2) Keystone species identification

By definition,^12, 27^ keystone species is one of the core driving factors to maintain or to damage the integrity and stability of the microbial communities. Therefore, the keystone species of the microbiome represent effective intervention targets. Here, we applied DIM to evaluate the impact of targeting each candidate keystone species on the entire gut microbiome based on the microbial interactions, so as to determine the keystone species that has the highest impact in the gut microbiome.

The topological importance of a species was a key indicator to evaluate its impact on other species in the entire ecosystem, and thus the important basis in our keystone species identification. To assess the strength of the microbial interactions obtained by causal inference, Hyperlink-Induced Topic Search (HITS), a network node importance evaluation algorithm^28^ was used to quantify the influence of every species in the community.^15^ The significances of HITS scores were assessed by permutation test with 1000 permutations. Species with significantly higher HITS scores (p ≤ 0.01) were defined as hub species.

As reported previously,^26^ gLV equation was applied to construct the dynamic model of microbial interactions and abundance changes in dynamic intervention modeling (DIM). DIM was conducted to evaluate the abilities of targeting each candidate keystone species on restoring a normal microbial structure from a diseased microbiome. The performance of the intervention by each candidate species, the intervention scores, were adjusted by integrating the HITS scores of the species in the normal microbiome to reward the interventions that preferentially restored the species with greater topological significance in the normal microbial network. Finally, the Iterative Feature Elimination (IFE) algorithm was used to determine the optimal combinations of the keystone species with the maximum cumulative intervention score (Supplementary Notes).

### Microbial functional enrichment analysis

With information of the gene annotations and the integrated modules from KEGG database, functional enrichment analysis was performed to assess the biological functions of microbial species. The KEGG Orthologs (KOs) information of microbes were predicted via PICRUST2.^29^ The enrichment analysis of KEGG modules was conducted with one-sided Fisher’s exact test, and adjusted with Benjamini-Hochberg method. Modules with FDR ≤ 0.01 were considered significantly enriched modules.

### Statistical analysis

All statistical analyses were conducted in Python (3.6.0). Statistical significance was determined by the two-sided Wilcoxon rank-sum test, Permutation test or one-sided Fisher’s exact test.

## RESULTS

### Abundance changes of the gut microbial species in obesity and NASH

Compared with normal controls, differential analysis identified 39 differential species in the gut of NASH patients (figure 2, table S2). These differential species mainly belonged to class Clostridiales, including Family XI, Lachnospiraceae, Ruminococcaceae, Peptostreptococcaceae, and Family III families. In NASH, the differential species of Family XI were up-regulated, while those of Lachnospiraceae and Ruminococcaceae were down-regulated (figure 2 and table S2). Similar changes were also observed in obesity. Interestingly, the gut microbiota exhibited a gradual change in the abundances of the differential species during the disease development from normal to obese and then to NASH (figure S1). In addition, these differential species possessed strong discrimination abilities for normal and NASH samples (figure S1), for example, *P. loveana*, *A. indistinctus(OTU57)* and *D. pneumosintes* achieved an AUC of 0.794, 0.766, 0.832, respectively.

**Figure 2.**
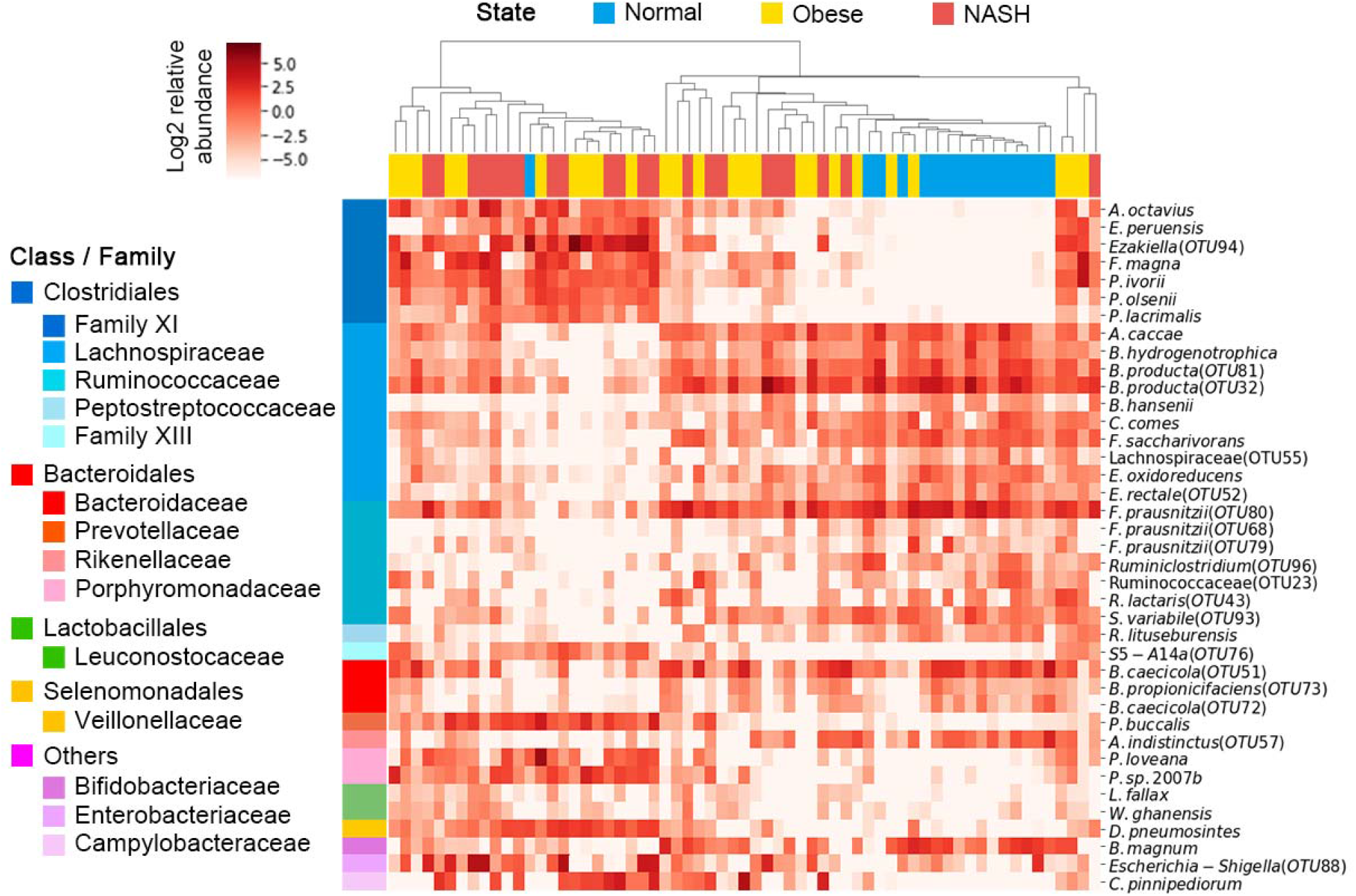
The abundance heatmap of species with significantly differential changes (FDR<0.05) between Normal and NASH. (Full names of the species: *Anaerococcus octavius, Ezakiella peruensis, Ezakiella(OTU94), Finegoldia magna, Peptoniphilus ivorii, Peptoniphilus olsenii, Peptoniphilus lacrimalis, Anaerostipes caccae, Blautia hydrogenotrophica, Blautia producta(OTU81), Blautia producta(OTU32), Blautia hansenii, Coprococcus comes, Fusicatenibacter saccharivorans*, Lachnospiraceae(OTU55), *Eubacterium oxidoreducens, Eubacterium rectale(OTU52), Faecalibacterium prausnitzii(OTU80), Faecalibacterium prausnitzii(OTU68), Faecalibacterium prausnitzii(OTU79), Ruminiclostridium(OTU96)*, Ruminococcaceae(OTU23), *Ruminococcus lactaris(OTU43), Subdoligranulum variabile(OTU93), Romboutsia lituseburensis, S5-A14a(OTU76), Bacteroides caecicola(OTU51), Bacteroides propionicifaciens(OTU73), Bacteroides caecicola(OTU72), Prevotella buccalis, Alistipes indistinctus(OTU57), Porphyromonas loveana, Porphyromonas sp. 2007b, Leuconostoc fallax, Weissella ghanensis, Dialister pneumosintes, Bifidobacterium magnum, Escherichia-Shigella(OTU88), Campylobacter pinnipediorum*)

### Distinct patterns of the microbial interactions in the gut of normal, obese and NASH subjects

Microbial interaction networks in the gut of the normal, obese and NASH subjects were constructed using a novel causal inference algorithm described in the Method section (figure 3A, B, C), and the hub species of each study group were then identified based on the network topological properties (figure 3D, E, F).

**Figure 3.**
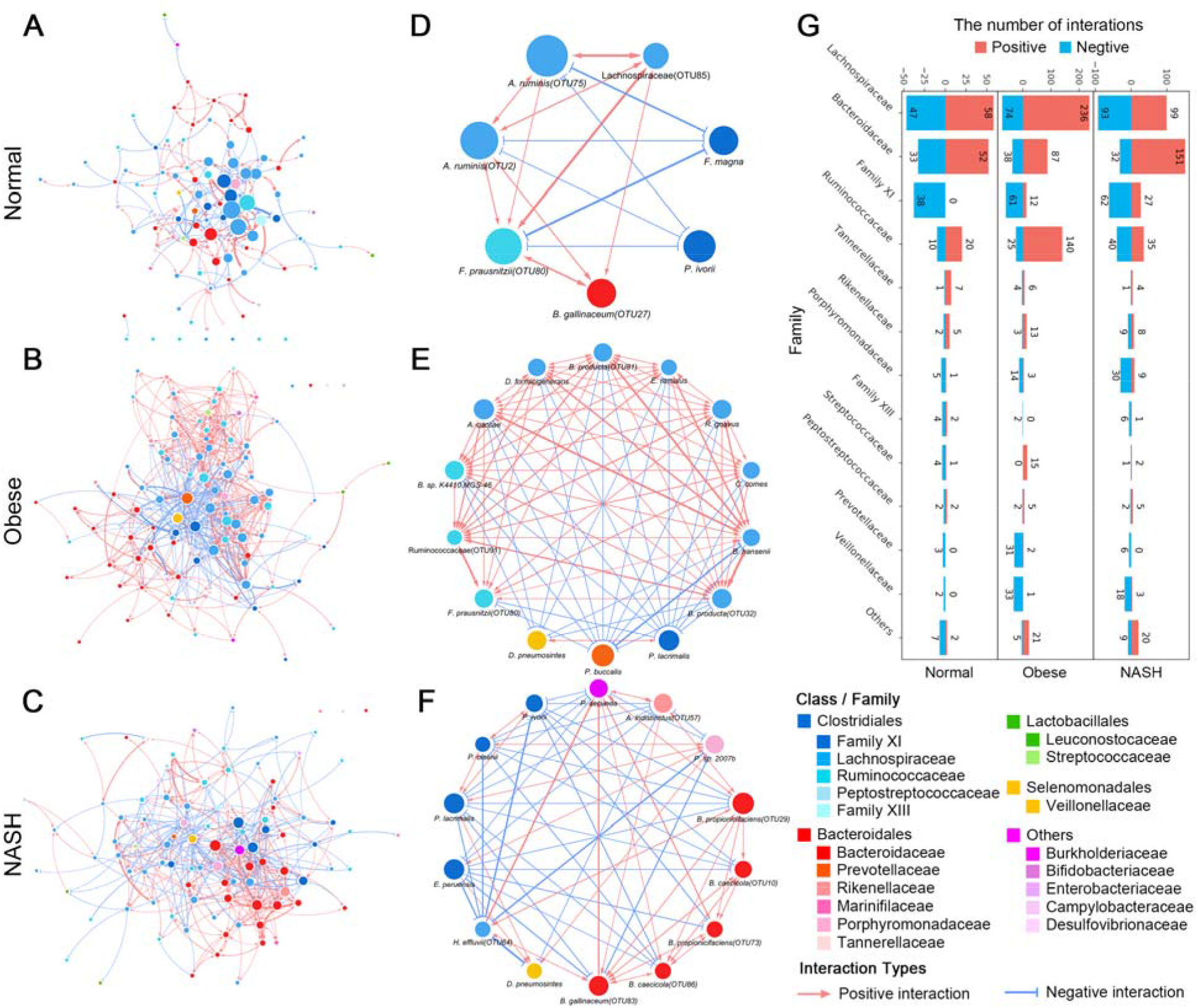
Microbial interaction networks and hub species in the gut of normal, obese and NASH subjects. Microbial interaction networks of normal (A), obese (B) and NASH (C) study groups and the corresponding sub-networks of hub species of normal (D), obese (E) and NASH (F) groups were shown. Colors of nodes indicated the taxonomy of the species and colors of edges indicated positive (red) or negative (blue) interaction between every two species. (G) The distribution of interactions for every family in microbial interaction networks of normal, obese and NASH.

The gut microbiome of normal, obese and NASH groups exhibited distinct patterns of microbial interactions (figure 3). In normal subjects, the microbial interaction network exhibited a heterogeneous pattern with less connections among the microbial species, characteristics of a typical scale-free network (figure 3A). The hub species that belong to Lachnospiraceae (Positive / Negative = 58/47), Ruminococcaceae (P / N = 20/10) and Bacteroidaceae (P / N = 52/33) mainly imposed positive impact, while those belong to Family XI (P / N = 0/38) exerted mainly negative impact on other microbes in the normal gut microbiome (figure 3G).

Compared to normals, the microbial interactions in the gut of obesity and NASH exhibited immense alterations with relatively homogeneous patterns and more connections among the members of the microbial community (figure 3B, C). Although the species of Lachnospiraceae, Family XI and Bacteroidaceae still remained the hub species in the networks, more hub species including species of the families Prevotellaceae, Veillonellaceae, Peptoniphilus, Porphyromonadaceae etc. were identified as hub species in obesity and NASH. The obesity and NASH specific hub species included *Prevotella buccalis*, *Alistipes indistinctus*, *Porphyromonas sp. 2007b*, and *Dialister pneumosintes* (figure 3D, E, F).

### DIM identified keystone species that drive the changes of the diseased gut microbiome toward a normal microbiome

Alteration in the abundance of a keystone species is expected to induce changes in other species and profoundly impact the intestinal homeostasis. Therefore, in this study, DIM was performed to simulate the dynamic alterations of the gut microbiome upon microbial intervention, and Iterative Feature Elimination (IFE) was conducted to identify those species that collectively exert the highest impact on the entire microbiome.

Consistent with their topologically central positions in the microbial interaction networks, the hub species were highly influential in the gut microbiome according to our DIM analyses (figure S2A and 4A). Targeting hub species in obese patients, such as *P. buccalis* (Intervention score, IS=0.397), *Ruminococcus torques* (IS=0.288), *Blautia hansenii* (IS=0.284), *Anaerostipes caccae* (IS=0.281), and *D*. *pneumosintes* (IS=0.279), were able to cause significant changes in the structures of the gut microbiome, toward restoring a normal gut microbiome (table S2). Similarly, with NASH gut microbiome, targeting hub species such as *P*. *sp. 2007b* (IS=0.376), *D*. *pneumosintes* (IS=0.294), *Peptoniphilus lacrimalis* (IS=0.289), *Agathobacter ruminis* (IS=0.254), and *Ezakiella peruensis* (IS=0.247) were able to cause significant changes in the gut toward restoring a normal gut microbiome (table S2). Targeting single species other than the hub species produced negligible impact on the composition of the gut microbiomes (figure S2C).

**Figure 4.**
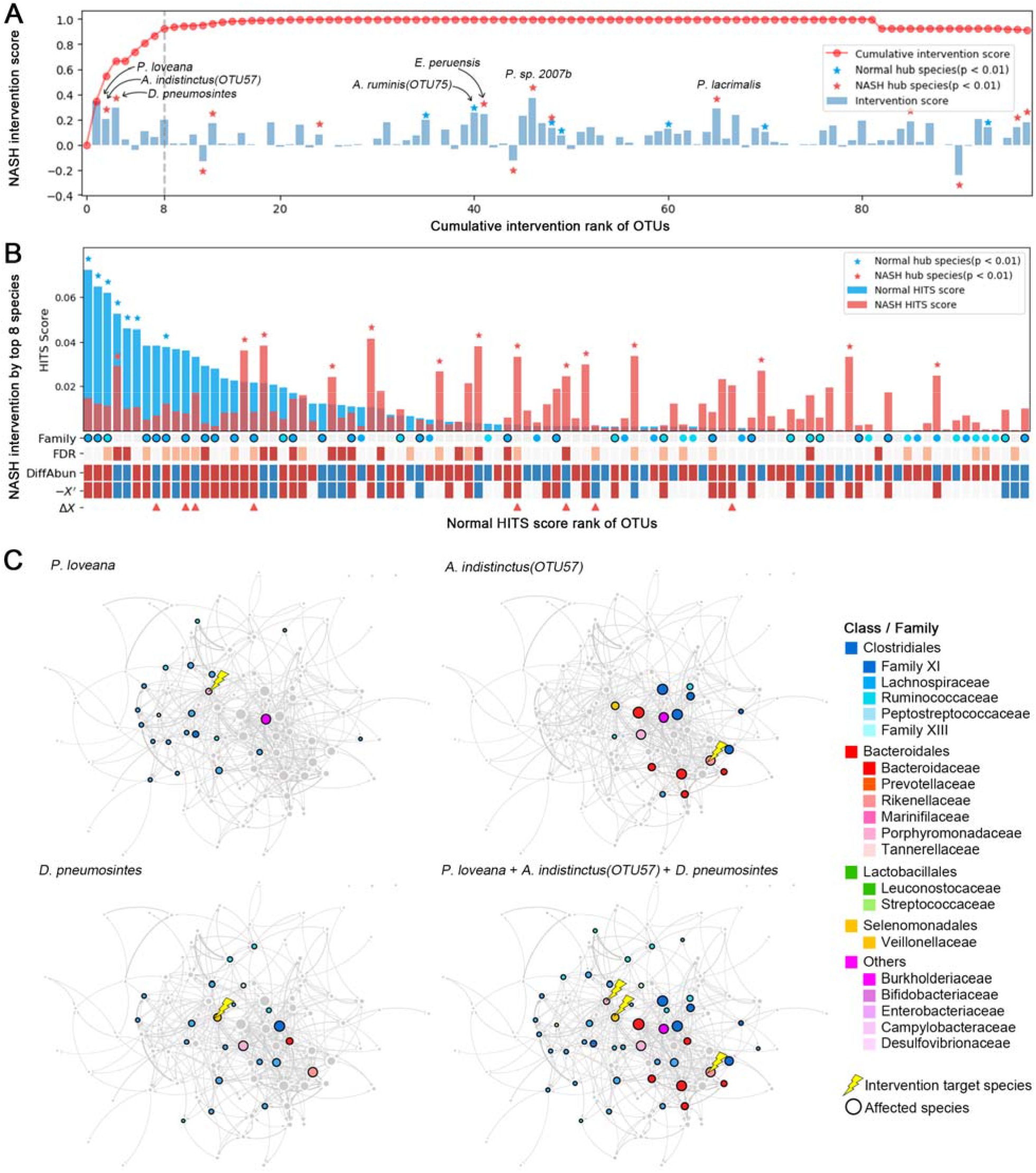
Dynamic intervention modeling of NASH microbiome. (A) Intervention scores (IS) of the gut microbes. The IS of each microbe was shown in the bar plot and the hub species were marked with stars. The red curve indicated the cumulative intervention scores (CIS) of the microbes sequentially selected by DIM. The first 8 keystone species, achieving a CIS greater than 0.9, were indicated by the dashed line. (B) Effect of microbial intervention on NASH microbiome according to DIM with the top 8 keystone species from (A). The topological importance (HITS scores) of species in normal microbiome were ranked in the bar plot. The species of Lachnospiraceae (Blue) and Ruminococcaceae (Light blue) were marked on the Family axis, with black borders indicating that the abundance recovered to normal levels after intervention. DiffAbun: abundance change from normal to NASH (Red: increase; Blue: decrease), with FDR indicated above (Red: FDR<0.01; Light red: FDR<0.05). -*X*’: negative representation of instant microbial abundance changes upon the intervention (Red: > 0; Blue: < 0). 8 keystone species for intervention were marked by triangles in Δ*X*. (C)The intervention effects of the first three keystone species (*P. loveana*, *A. indistinctus* and *D. pneumosinte*).

Intervention targeting multiple species simultaneously would produce a better outcome than targeting a single species.^11^ Therefore, we took a novel approach of integrating the DIM and IFE algorithms to identify the keystone species combinations that have the highest potential for microbial interventions. We identified 11 and 8 keystone species from the obese and the NASH microbiomes, respectively. Cumulative intervention scores (CIS) of the identified keystone species combinations were 0.903 and 0.923, respectively (figure S2A, 4A and table S2), suggesting that the microbiome in the gut of the patients can be maximally restored toward a normal microbiome by targeting these keystone species (figure S2B and 4B). Targeting 8 keystone species in the NASH caused changes in the species of Lachnospiraceae and Ruminococcaceae, that are the hub species in normal microbiome, to increase toward the abundances found in normal microbiome (figure 4B). Meanwhile, hub species in NASH microbiome also responded to the intervention, and most of the differential species in NASH responded to the keystone intervention with abundances changed toward normal microbiome.

Among 8 NASH keystone species, the first three keystone species, *P. loveana*, *A. indistinctus*, and *D. pneumosintes*, exhibited strong intervention capacity (CIS=0.664), and the other five keystone species played minor roles (figure 4A). Similarly, the first three keystone species of obesity exhibited strong intervention capacity (CIS=0.587, figure S2A). The first three keystone species were hub species (p<0.05) in normal or diseased microbial network, and they caused major changes when they are targeted for microbial intervention in NASH patients (figure 4C and table S3). *P. loveana* was elevated in NASH, therefore, we removed *P. loveana* as a microbial intervention. Removing *P. loveana* mainly increased the abundance of the Lachnospiraceae species, most of which were the hub species in normal microbiome. *A. indistinctus* was decreased in NASH. Adding *A. indistinctus* mainly elevated species of Bacteroidaceae and reduced Family XI. *D. pneumosintes* was elevated in NASH. Removing *D. pneumosintes* increased the abundances of Lachnospiraceae and Ruminococcaceae species. Intervention with these three species simultaneously could restore the abundances of many gut microbes, especially the species of Lachnospiraceae and Ruminococcaceae (both being abundant families in normal microbiomes), thereby promoting the reconstruction of a normal intestinal microbiome (figure 4C).

### Potential mechanisms for the keystone species to impact the NASH microbiome

*P. loveana*, *A. indistinctus* and *D. pneumosintes* were identified as the top keystone species in the NASH microbiome, and exhibited the highest capabilities for the intervention of the NASH microbiome. In order to understand the mechanisms behind the massive alterations in the microbiome induced by targeting these keystone species, we obtained their genome information with PICRUST2,^29^ and performed the functional (KEGG Module) enrichment analysis (figure 5A and table S4).

**Figure 5.**
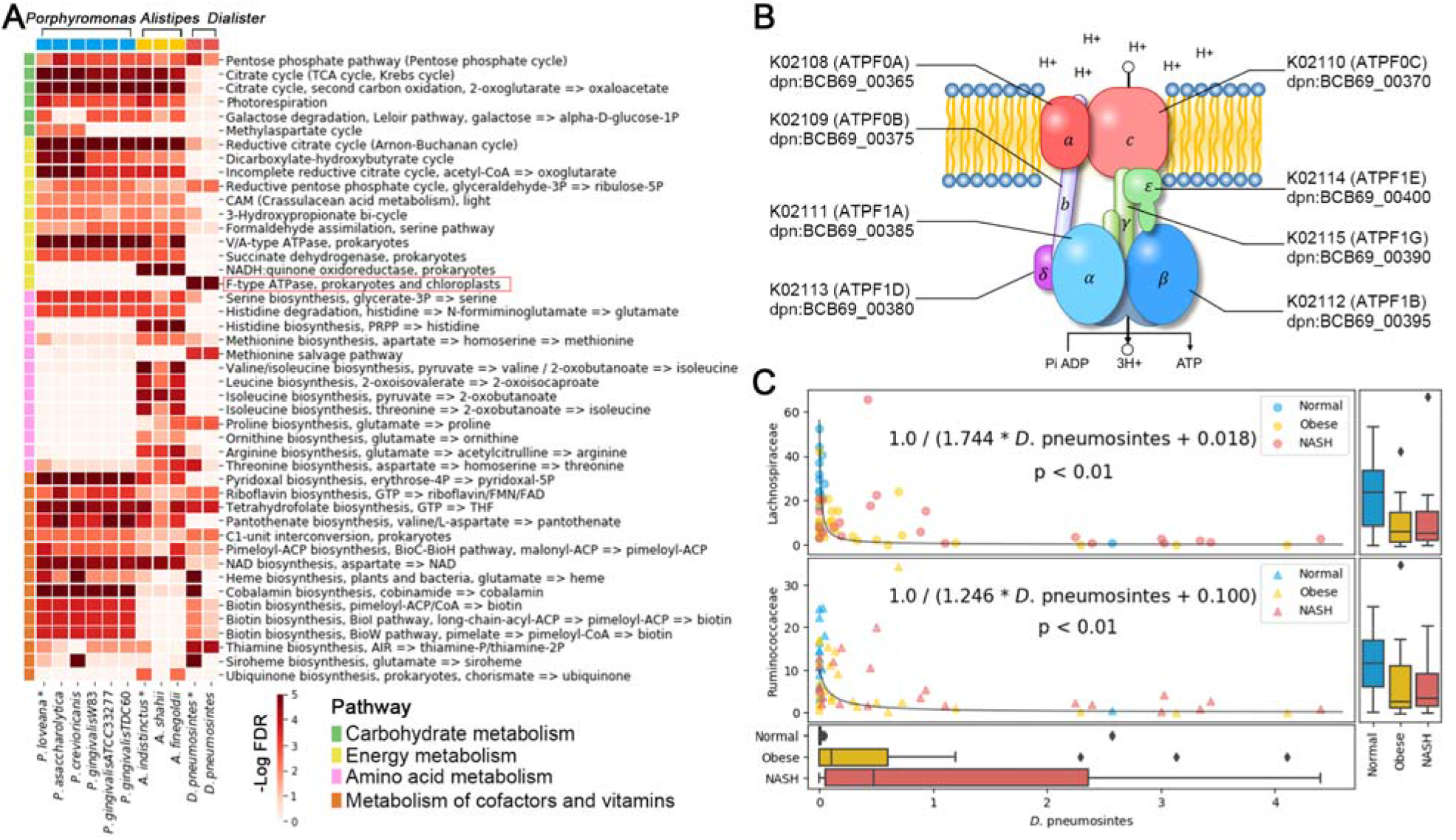
Functional enrichment analysis on keystone species. (A) The species *P. loveana*, *A. indistinctus* and *D. pneumosintes* were subjected to KEGG module enrichment analysis based on their gene (KO) annotations using PICRUST2. Modules were classified according to the KEGG pathways and marked with different colors. The heatmap plotted the significance of enrichment (−log FDR). (B) *D. pneumosintes* encodes the entire eight genes for F-type ATPase complex. This protein complex can utilize the membrane proton gradient for ATP production. (C) The abundance change of *D. pneumosintes* was significantly correlated to the abundance changes of Family Lachnospiraceae and Ruminococcaceae, fitting with a reciprocal function in both cases. The box plots at the axis are the abundance distribution of Lachnospiraceae, Ruminococcaceae and *D. pneumosintes* in the gut of normal. obese and NASH subjectes.

As shown in figure 5A, the genome of *P. loveana* is severely deficient in amino acid production genes while rich in cofactors and vitamins production genes. Amino acids, such as glutamate, glycine, alanine, tyrosine, aspartate, valine, etc., are required substrates for the production of cofactors and vitamins,^30, 31^ and may be used for the production of other molecules including short-chain fatty acids (SCFA).^32^ Therefore, the increased abundance of *P. loveana* would lead to enhanced amino acid consumptions in NASH. A similar gene enrichment pattern was observed for the keystone *D. pneumosintes*, indicating that *D. pneumosintes* may contribute to elevated amino acid consumption in NASH.

Keystone species *A. indistinctus* participated in the production of a variety of amino acid including serine, threonine, valine, isoleucine, leucine, arginine, proline, glutamate and histidine (figure 5A). The down-regulation of *A. indistinctus* (FDR=0.020, table S2) suggested reduced microbial synthesis of amino acids.

Importantly, *D. pneumosintes* encodes the entire eight genes for F-type ATPase complex (figure 5A, B), implicating that *D. pneumosintes* can utilize H^+^ in the intestinal lumen to synthesize ATP. In our intervention simulation, targeting *D. pneumosintes* caused the changes in the abundances of Lachnospiraceae and Ruminococcaceae species (figure 4C and table S3). Consistently, correlation analyses showed that the abundance of *D*. *pneumosintes* was negatively correlated with those of Lachnospiraceae and Ruminococcaceae (p<0.01, figure 5C). Previous studies have shown that the Lachnospiraceae and Ruminococcaceae species are very sensitive to environmental pH, which seriously affects their abilities of butyric acid production.^33, 34^ Therefore, the increased abundance of *D. pneumosintes* in NASH (FDR=0.004, table S2) might change the intestinal pH, and consequently decrease the abundance and butyric acid production of Lachnospiraceae and Ruminococcaceae.

### Validation of the method for keystone species identification with an independent NAFLD cohort

We observed a similar pattern of microbial change in the validation NAFLD cohort (table S5). Briefly, Lachnospiraceae and Ruminococcaceae species exhibited significantly reduced abundance in the gut of NAFLD patients (p<0.01). In addition, *A*. *indistinctus*, a keystone species in the NASH microbiome in the discovery cohort, was also significantly down-regulated (p=0.01 in normal *vs*. NASH-cirrhosis and p=0.017 in normal *vs*. NAFLD without fibrosis, figure S3A and table S5). These differential species also exhibited strong sample discrimination abilities with the highest AUC=0.894 (Erysipelotrichaceae(OTU96) in Normal *vs*. NASH-cirrhosis, figure S3B).

Again, the Lachnospiraceae and Ruminococcaceae species were the hub species of the normal gut microbiota, playing key roles in maintaining the homeostasis of the normal microbial communities (table S5).

The keystone species identified from the validation cohort were partly overlapping with those identified from the discovery cohort. The common keystone species from both cohorts included *B. producta*, *B. barnesiae* and *A. caccae* (table S5). Consistently, targeting these keystone species for microbial intervention was able to restore the abundances of Lachnospiraceae and Ruminococcaceae (figure S4).

## DISCUSSION

Here we report a new algorithm for the keystone species identification in the gut microbiome, based on current causal inference theories and the DIM with gLV model. We identified the NASH keystone species combination, represented by *P*. *loveana*, *A*. *indistinctus*, and *D*. *pneumosintes*, that showed the highest potential for the microbial intervention of NASH.

The most outstanding characteristic of the gut microbiome in both adolescent (discovery cohort) and adult (validation cohort) NAFLD seemed to be decreased abundances in Lachnospiraceae and Ruminococcaceae, two dominant families in Clostridiales. As the major butyrate-producing bacteria in the intestine,^35^ Lachnospiraceae and Ruminococcaceae, may play important roles in suppressing intestinal inflammation via the stimulatory effect of butyrate on T regulatory cells in the mucosa,^36^ and consequently suppress the pathogenesis of NASH.^37, 38^ The structural and functional importance of these two bacterial families make them desirable targets in the microbial intervention of NAFLD.

With NASH microbiome, the keystone species, especially *P. loveana*,*A. indistinctus* and *D. pneumosintes*, could rapidly alter the abundance of Lachnospiraceae and Ruminococcaceae species and restore the microbial composition toward a normal gut microbiome. These species are able to impact the other community members with fermentation products. *P. loveana* was elevated in NASH, and therefore was removed from the NASH microbiome for intervention. Reduced abundance of *P. loveana* leads to decreased consumption of amino acids, therefore, leaving more resources for the growth of other bacteria including Lachnospiraceae and Ruminococcaceae. *A. indistinctus* was decreased in NASH, and therefore was added to the NASH microbiome for intervention. *A. indistinctus* is equipped with many genes related to amino acid synthesis. In addition to support protein synthesis of the microbial communities, these amino acids produced in the gut may serve as microbial fermentation substrates for SCFA production,^39^ such as butyrate synthesis from threonine, lysine, and glutamate.^40^ As such, the increased presence of *A. indistinctus* may promote not only the growth of other members of the microbial community with its amino acid production, but also the intestinal balanced immunity with SCFA production.^37, 38^ *D. pneumosintes* was elevated in NASH, and therefore was removed from the NASH microbiome for intervention. *D. pneumosintes* encodes all eight genes of the F-type ATPase complex that can use protons in the intestinal environment for ATP synthesis. Thus reducing *D. pneumosintes* may help maintain a low intestinal pH that promotes the growth of the butyrate-producing Lachnospiraceae and Ruminococcaceae species.^33, 34^

Compared to the keystone species with the top CIS scores, the abundances of Lachnospiraceae and Ruminococcaceae species were more profoundly altered in the NASH microbiome, yet they did not achieve the highest IS in DIM and their performance in IFE were not impressive either. Our results indicate that the keystone species may not be the most abundant species in the microbial community, and that targeting species mostly altered in disease may not be an effective microbial intervention strategy. In contrast, *P. loveana*,*A. indistinctus* and *D. pneumosintes*, the keystone species that out-performed other keystone species in IFE, were not the top altered species in disease, but they exhibited the highest potential in restoring a normal gut microbiome. Their ability for microbial intervention of NASH may be attributed to their metabolic products that have profound influence on other members of the microbial community, and these broad influences allowed them the special roles in maintaining the integrity and stability of the normal microbiome.

In summary, we proposed a novel algorithm for microbial keystone identification from cross-sectional microbiome data based on causal inference analysis and DIM. The identified keystone species in the gut of NAFLD, represented by *P. loveana*, *A. indistinctus* and *D. pneumosintes*, could efficiently modulate the microbial composition of the NAFLD, especially Lachnospiraceae and Ruminococcaceae, toward a normal gut microbiome. Validated with an independent NAFLD cohort, our method suggested a novel potential microbial treatment for NAFLD. Our method for microbial keystone species identification may benefit microbiome studies in the broad fields of medicine, environmental science and microbiology.

## Contributors

LZ, RL and RZ conceived and designed the project. Each author has contributed significantly to the submitted work. DW and NJ drafted the manuscript. RZ, YZ, WG, SF, YL, SC, CT, PL, RL and LZ revised the manuscript. All authors read and approved the final manuscript.

## Patient consent for publication

Not required.

## Data and materials availability statement

The raw sequencing reads and associated meta-data are archived at MG-Rast (http://metagenomics.anl.gov/linkin.cgi?project=1195) and EMBL-EBI (https://www.ebi.ac.uk/ena/data/view/PRJEB28350). The code used in this study is available on GitHub at https://github.com/ddhmed/NAFLD_keystone.

