## Supplementary for "Identification of the keystone species in non-alcoholic fatty liver disease by causal inference and dynamic intervention modeling"

### Supplementary Materials

#### Supplementary Notes

**Microbial interaction network construction based on causal inference theories**

Based on Robins'[1] and Pearl's[2] causal inference theories, and the generalized Lotka–Volterra (gLV) dynamics model for characterizing microbial interactions,[3] we designed an algorithm for microbial interaction network construction for cross-sectional data. The specific implementation process is as follows.

**Construction of microbial interaction graphical model based on correlation analysis.** Correlation analyses were performed to construct the graphical model of microbial interactions. In consideration of the characteristics of the microbial sequencing data, SparCC[4] with its outstanding performance for microbial compositional data was chosen to construct the microbial co-occurrence networks with permutation test (1000 permutations). Significant microbial relationships (p<0.01) are extracted to construct the co-occurrence networks, which is the priori knowledge for causal inference analyses.

**Identification of causality between species**. Unlike parameter estimation in machine learning, the core of causal inference theories is "identification". Based on the graphical model,[2] we exhausted all feasible strategies to identify the potential causal relations, such as back-door criterion, front-door criterion, and do-calculus.

**Estimation of causal effect sizes**. The strength of the causal effect was estimated with a microbial interaction model. The generalized Lotka–Volterra model (Equation 1), a classical dynamic model, was applied to estimate the causal effect sizes between microbes.

$$x_i' = x_i(A_i X + r_i) \tag{1}$$

This gLV equation describes the dynamic changes of taxa abundance regulated by their interactions. Conside s often stable and resilient,[5] which means $x_i' \approx 0$.

$x_i' = x_i\left(\sum a_{ij}x_j + r_i\right) \approx 0$ r a microbial community of N different taxa, where $x_i'$ is the growth rate of taxon $i$; $A_i = (a_{i1}, \dots, a_{iN})$ is the interaction vector, representing the integrated regulatory effect of all taxa on taxon $i$; $x_i$ is the abundance of taxon $i$; $X = (x_1, \dots, x_N)$ is the abundance vector of all taxa; $r_i > 0$ is the inherent growth rate of taxon $i$.

The gut microbiome of normal or disease state i (2)

Where $a_{ij}$ is the interaction coefficient (per capita effect) of taxon $j$ on the growth rate of taxon $i$. In this work, we consider $x_i \neq 0$ in NASH state. Let $s_j = \frac{a_{ij}}{-a_{ii}}$ and $t_i = \frac{r_i}{-a_{ii}}$. From equation 2, we obtain:

$$x_i \approx \sum_{j\neq i} s_j x_j + t_i \tag{3}$$

This equation implies linear relationship between the abundance of taxa in the steady-state. Combined with the causal relationship identification from step 2, we could estimate the causal effect size between taxa through linear regression.

$$x_i = \sum_{j\neq i} s_j x_j + t_i \quad \text{, where } s_j \text{ was defined as the causal effect of } j \text{ on } i \tag{4}$$

The estimation of interaction effect sizes based on the gLV equation allows the subsequent implementation of the dynamic intervention model which is also based on the gLV equation.

**Iterative optimization of causal inference**. The significance of causality was assessed by the permutation test. On the other hand, in order to achieve higher accuracy in causal inference, an iterative optimization strategy was implemented to improve the graphical model, making use of the significant (p≤0.01) causal pairs with a certain learning rate (default 0.5). The adjusted graphical model was then subjected to a new round of causal inference. These steps were repeated until convergence.

The above algorithm was developed and implemented based on the DoWhy (https://microsoft.github.io/dowhy/) causal inference framework developed by Microsoft. All codes of this algorithm are available at the online python project (https://github.com/ddhmed/NAFLD_keystone).

**Keystone species identification based on dynamic intervention modeling**

Keystone species are defined as the species required for maintaining the homeostasis of ecological communities. The alteration of the keystone species could affect the entire community through the interactions among the members of the ecosystem.[6] Therefore, keystone species may be targeted in microbial interventions. We proposed to identify the keystone species with a dynamic intervention modeling (DIM) algorithm with cross-sectional data, as detailed in the following four steps:

**Topological importance evaluation of the species in the interaction networks**. The microbial interaction networks of normal and diseased states were constructed by causal inference, in which the impact of each microbe on the community could be described by network topological importance. Microbial interaction network constructed by causal inference was a directed graph that contains effect intensity and direction. Therefore, the HITS algorithm which computes authorities (Equation 5) and hubs (Equation 6) for nodes in the network was applied[7]. HITS hubs score was used as microbial topological importance score and its significance was evaluated by permutation test with 1000 random networks that have an equal number of nodes and interactions.

$$a(u) = \sum h(v), \quad a(u) = \frac{a(u)}{\max(a(u))} \tag{5}$$

$$h(v) = \sum a(u), \quad h(v) = \frac{h(v)}{\max(h(v))} \tag{6}$$

**Dynamic intervention simulation.** With the cross-sectional data, we are able to implement the gLV modeling by focusing on the characteristics of the steady-state of the diseased microbiome (Equation 2). With this approach, we evaluated the impact of the intervention on microbial species at the diseased microbiome by introducing the intervention operation targeting each potential keystone species (Equation 7).

$$x_i' = x_i\left[\sum a_{ij}(x_j + \Delta x_j) + r_i\right] \tag{7}$$

Where $\Delta x_j$ is the intervention against taxon $j$. Considering that the microbiome is still in steady state at the moment of intervention, $x_i' \approx 0$. And the effect of the intervention on taxa could be expressed as:

$$x_i' = x_i \sum a_{ij}\,\Delta x_j, \quad \text{where } \Delta x_j = -DiffAbun_j \tag{8}$$

Where the $DiffAbun_j$ indicates the abundance change of taxon $j$ from normal to the disease state.

**Intervention scoring**. In order to comprehensively evaluate the effectiveness of interventions, we designed the intervention score ($IS$) based on the changes of taxa during the intervention.

$$IS = sign(DiffAbun) * \ sign(-X') * HITS\_Score_{normal} \tag{9}$$

Where the $DiffAbun$ is a vector, representing the abundance change of all taxa from normal to disease. And $X'$ is the vector of abundance change of all taxa after intervention. $HITS\_Score_{normal}$ is the topological importances of taxa in the normal state. $IS$ reflects the potential of the intervention to restore the microbiome to normal state.

**Searching for the optimal combinations of the keystone species for microbial intervention**. Iterative Feature Elimination (IFE), a feature selection strategy based on the greedy algorithm, was used to search for the optimal combination of microbial species for the intervention. The specific operation of the IFE is as follows. First, the intervention score of all taxa, $IS_{all}$, is calculated. Then, one taxon was removed from the current combination each time and the intervention score of the remaining taxa combined, $IS_{leave-one-out}$, was calculated, and the combination with the highest score, $argmax(IS_{leave-one-out})$, was retained for the next removal operation. Repeat the above steps until the optimal combination of the keystone species for microbial intervention was found.

The above algorithm was developed and implemented in python. All codes of this algorithm are available in the online project (https://github.com/ddhmed/NAFLD_keystone).

**Supplementary Figures**

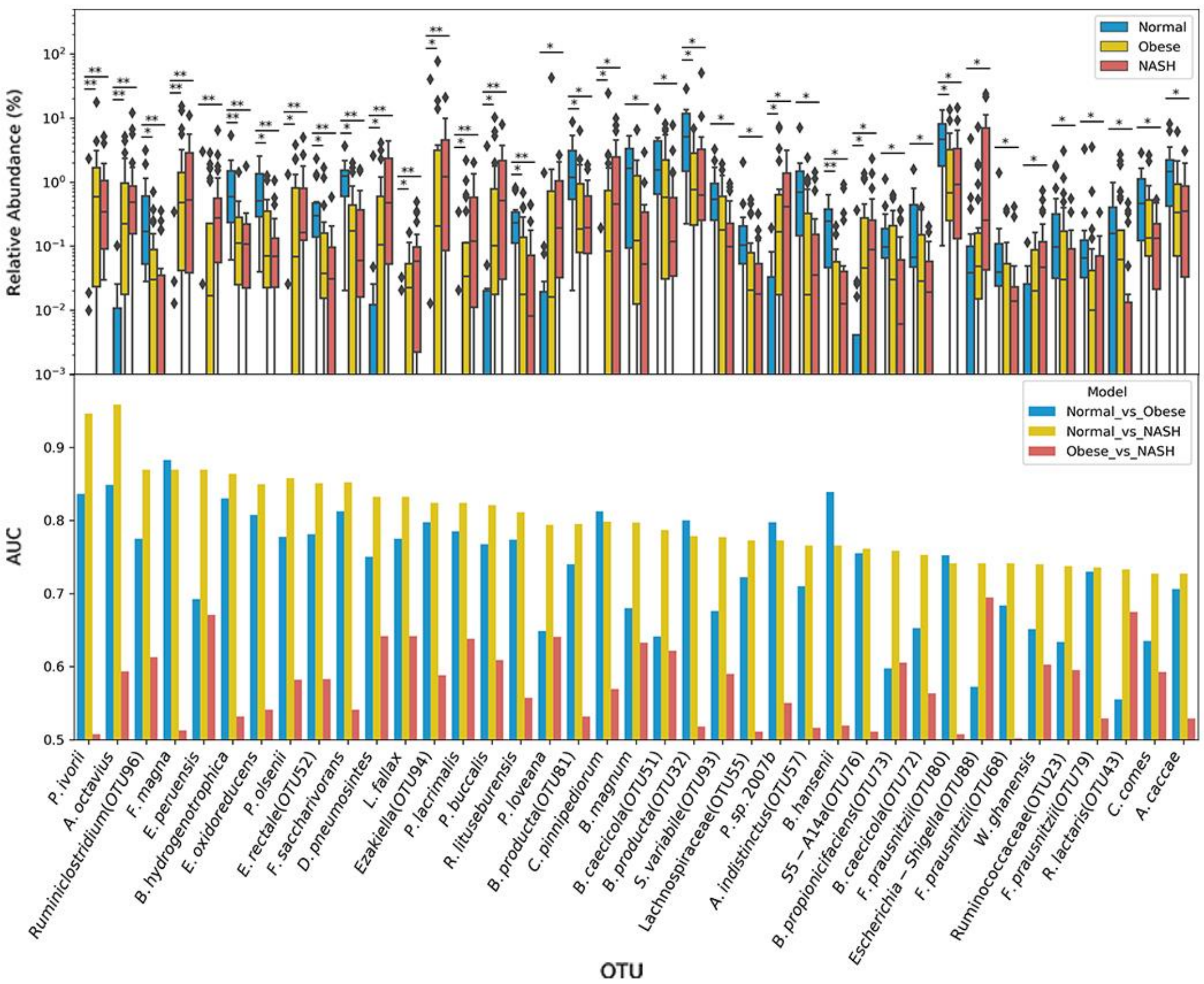

**Figure S1 The abundance changes of the differential microbes (FDR<0.05, upper) and their sample discrimination abilities (bottom).**

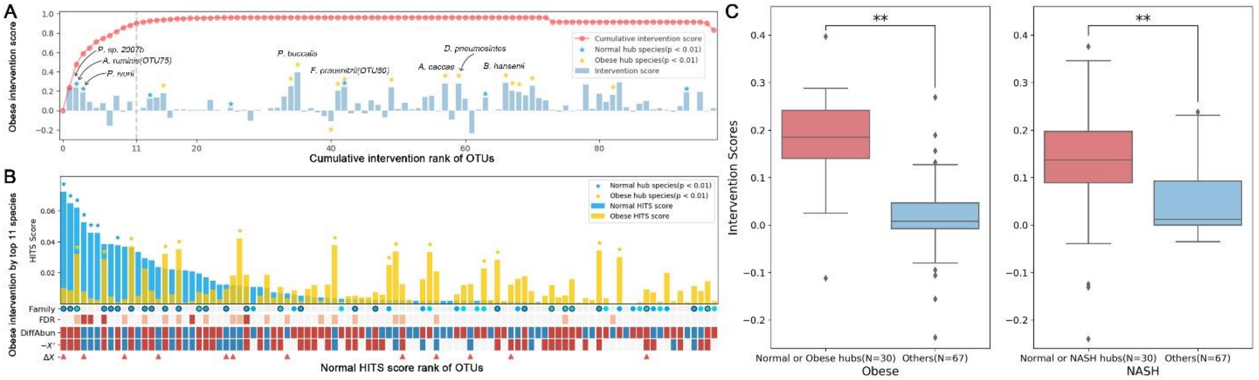

**Figure S2 Dynamic intervention modeling of obesity microbiome.** (A) Intervention score (IS) of microbes in obesity. The IS of each microbe was shown in the bar plot and the hub species were marked with stars. The red curve indicated cumulative intervention scores (CIS) of the microbes sequentially selected by DIM. The first 11 keystone species, achieving the cumulative intervention score>0.9, were indicated by the dashed line. (B) Effect of microbial intervention on obesity microbiome according to DIM with the top 11 keystone species from (A). The topological importances (HITS scores) of species were ranked in the bar plot. The species of Lachnospiraceae(Blue) and Ruminococcaceae (Light blue) were marked in the Family axis. Nodes with black borders indicated that the abundance of species recovered to normal levels after intervention. DiffAbun: abundance change from normal to obesity (Red: increase; Blue: decrease), with FDR indicated above (Red: FDR<0.01; Light red: FDR<0.05). $-X'$: negative representation of instant microbial abundance changes upon the intervention (Red: > 0; Blue: < 0). 11 keystone species for intervention were marked by triangles in $\Delta X$. (C) Distribution of the intervention scores of the microbial species. ** indicates p<0.01 with the Wilcoxon rank-sum test.

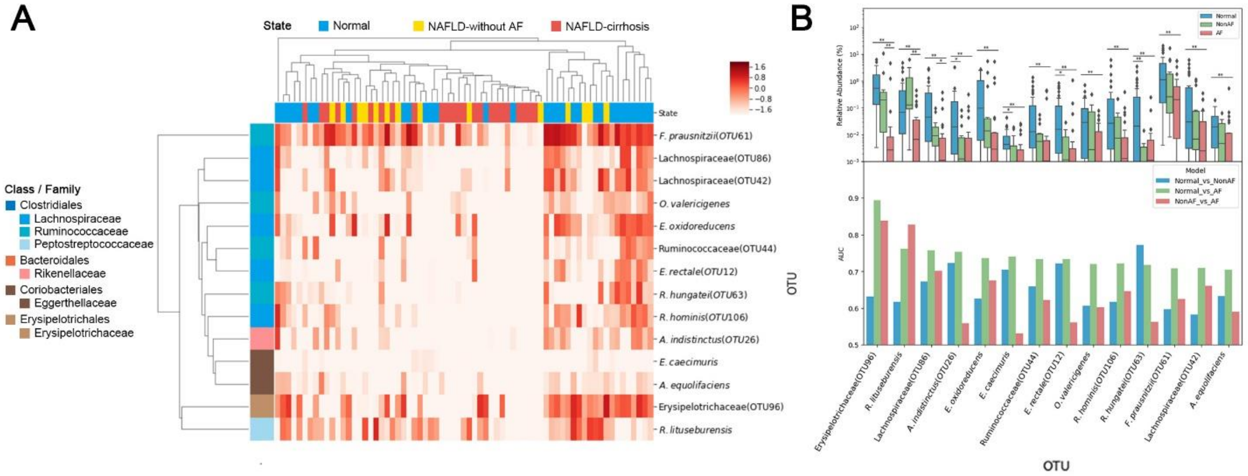

**Figure S3 The differential abundance analysis in the NAFLD cohort from California.** (A) The abundance heatmap of the differential species (p<0.01) between normal and NAFLD-cirrhosis. (B) The abundance changes of differential microbes (p<0.01, upper) and their sample discrimination abilities (bottom) between normal and NAFLD-cirrhosis.

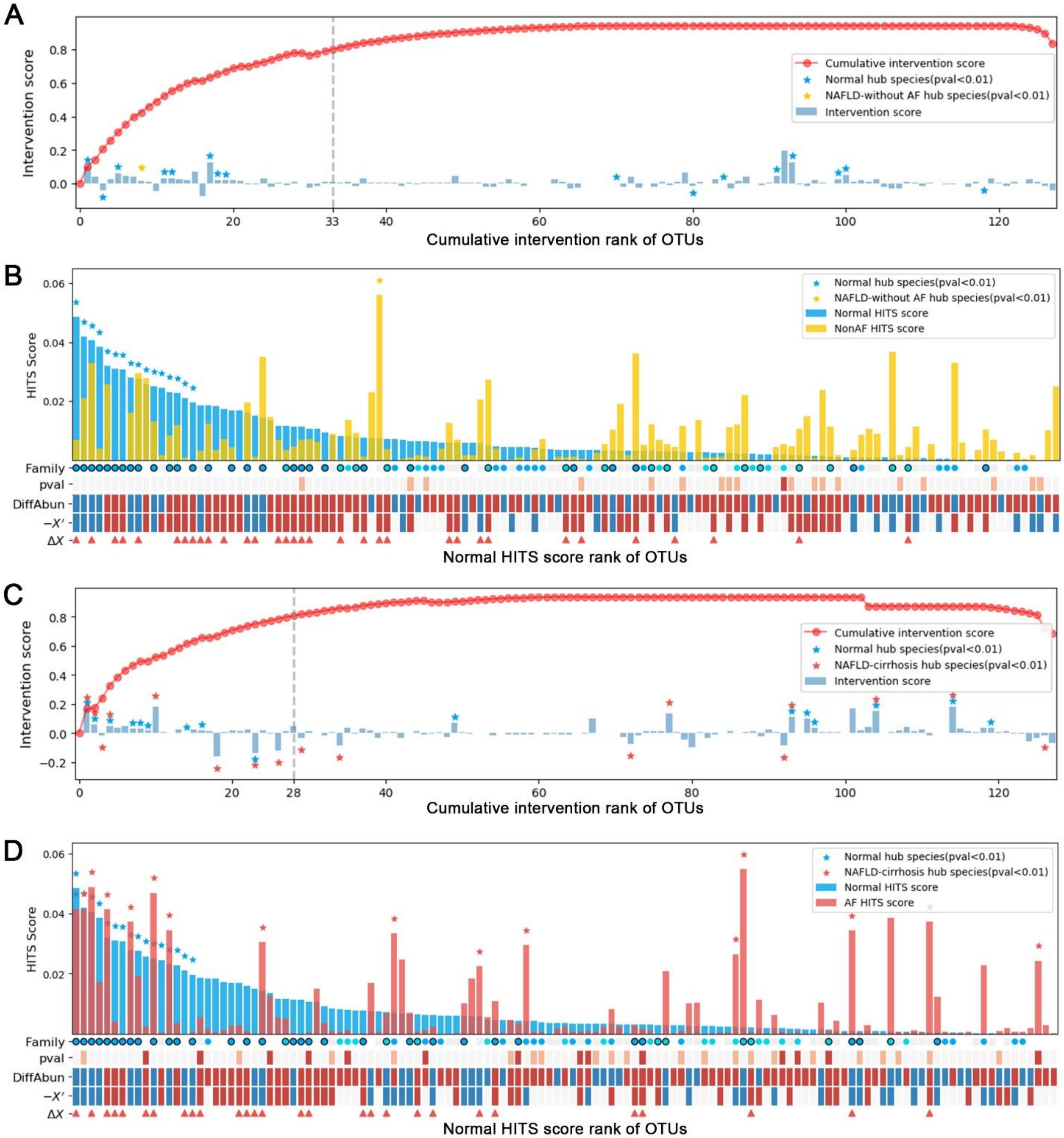

**Figure S4 Dynamic intervention modeling based on microbial interaction network of the validation NAFLD cohort.** (A) Intervention score of microbes in NAFLD-without AF. (B) Effect of microbial intervention with keystone species in NAFLD-without AF. (C) Intervention score of microbes in NAFLD-cirrhosis. The first 28 keystone species, achieving the cumulative intervention score>0.8, were indicated by the dashed line. (D) Effect of microbial intervention with keystone species in NAFLD-cirrhosis.

### Supplemental Tables

**Table S1. Characteristics of the study groups of the discovery and validation cohorts.**

| | New York cohort | | | California cohort | | |
|---|---|---|---|---|---|---|
| Parameter | Normal | Obese | NASH | Normal | NAFLD-without AF | NAFLD-cirrhosis |
| Samples | 16 | 25 | 22 | 31 | 14 | 24 |
| Sex(F/M) | F6/M10 | F12/M13 | F10/M12 | 5M/26F | 3M/11F | 5M/19F |
| Age(year) | 14.4±1.8 | 12.7±3.2 | 13.6±3.5 | 61.68±0.26 | 59.50±0.75 | 67.75±0.32 |
| BMI | 20.4±0.1 | 33.4±0.3 | 34.0±0.4 | 27.83±0.24 | 31.41±0.50 | 33.14±0.44 |
| AST | ND | 28.4±0.6 | 51.7±1.3 | 23.90±1.77 | 21.64±1.64 | ND |
| ALT | ND | 27.7±0.6 | 66.9±1.9 | 19.35±1.24 | 20.42±1.58 | ND |

Data are Mean ± standard error.

Abbreviations: F, female; M, male; AST, aspartate aminotransferase; ALT, alanine aminotransferase; ND, not determined; AF, advanced fibrosis.

**Table S2. Differential abundance, HITS and intervention scores of the species in discovery cohort.**

| Abb. | Taxonomy | | | | Relative Abundance (%) | | |
|---|---|---|---|---|---|---|---|
| | OTU | Class | Family | Species | normal_abun | obese_abun | nash_abun |
| *P. loveana* | f6944e5be5cca661767014ab95867c4dc3848235 | Bacteroidales | Porphyromonadaceae | *Porphyromonas loveana* | 0.101 | 1.966 | 0.698 |
| *A. indistinctus(OTU57)* | 9075bd7b4542aad1faf685d8960946088cca0810 | Bacteroidales | Rikenellaceae | *Alistipes indistinctus(OTU57)* | 1.109 | 0.343 | 0.196 |
| *D. pneumosintes* | c84919371da9c309c2b7a50ab6ed836c0c9555e6 | Selenomonadales | Veillonellaceae | *Dialister pneumosintes* | 0.167 | 0.583 | 1.212 |
| *A. caccae* | 4c6d7a5409ae36dbf1be703072de0fc8872933ca | Clostridiales | Lachnospiraceae | *Anaerostipes caccae* | 1.946 | 0.771 | 0.587 |
| Lachnospiraceae(OTU85) | d32c3a758a590f2e70b54af0f90eceb9730958bc | Clostridiales | Lachnospiraceae | Lachnospiraceae(OTU85) | 0.087 | 0.311 | 0.191 |
| *B. barnesiae* | 795dbf8c019483a25ae9a8827a62a936185d84a0 | Bacteroidales | Bacteroidaceae | *Bacteroides barnesiae* | 1.275 | 2.301 | 1.055 |
| *Escherichia-Shigella(OTU88)* | b1c85b23cb036d5783eb5cb420b42d5816b804f1 | Enterobacteriales | Enterobacteriaceae | *Escherichia-Shigella(OTU88)* | 0.385 | 0.536 | 4.909 |
| *B. producta(OTU32)* | b4039debc95fcd3157de7b17572bed80a93ae889 | Clostridiales | Lachnospiraceae | *Blautia producta(OTU32)* | 6.924 | 1.691 | 3.981 |
| *F. prausnitzii(OTU68)* | 7fe4f3b04e35b20bb4516344debfbb6154e330d3 | Clostridiales | Ruminococcaceae | *Faecalibacterium prausnitzii(OTU68)* | 0.142 | 0.050 | 0.043 |
| *B. caecicola(OTU84)* | 4973d0786e190acfd6718eb8bf6732cb978c023c | Bacteroidales | Bacteroidaceae | *Bacteroides caecicola(OTU84)* | 0.249 | 0.248 | 0.137 |
| *B. coprosuis(OTU62)* | 9e6194334ec79f0f19d760dc2ac0e90c42aa0e7d | Bacteroidales | Bacteroidaceae | *Bacteroides coprosuis(OTU62)* | 0.738 | 0.368 | 0.336 |

| | | | | | | | |
|---|---|---|---|---|---|---|---|
| *B. caecicola(OTU86)* | 3ca0f6463b44a9076deedb b5f3bb7ed00fcb358d | Bacteroidales | Bacteroidaceae | *Bacteroides caecicola(OTU86)* | 0.044 | 0.071 | 0.111 |
| *B. propionicifaciens(OTU73)* | 1c2c5de41c9879a9f2929f 356acca72066f2e209 | Bacteroidales | Bacteroidaceae | *Bacteroides propionicifaciens(OTU 73)* | 0.193 | 0.148 | 0.060 |
| *M. faecis* | fde0738b497ff50d42b1a7 c7a66a1d1576c53494 | Clostridiales | Lachnospiraceae | *Merdimonas faecis* | 0.036 | 0.067 | 0.043 |
| *C. pinnipediorum* | 3a21d49f9431e24145b80 c43adc46af78fd5f00f | Campylobacterale s | Campylobacteraceae | *Campylobacter pinnipediorum* | 0.012 | 1.457 | 2.101 |
| *R. lituseburensis* | 4a750c3b25e9ba149a232 3ba3b62ee0ca7940d83 | Clostridiales | Peptostreptococcaceae | *Romboutsia lituseburensis* | 0.288 | 0.100 | 0.073 |
| *A. indistinctus(OTU20)* | 69f5addfb12c83bce0beaa 7a738668e8a772480e | Bacteroidales | Rikenellaceae | *Alistipes indistinctus(OTU20)* | 0.380 | 0.154 | 0.060 |
| *C. catus* | 6172539e0c22e9c7314bc 68485a8fc72755422d5 | Clostridiales | Lachnospiraceae | *Coprococcus catus* | 0.067 | 0.041 | 0.100 |
| *E. ramulus* | 82d482e629b615acdbb8a 62b39115e47040c7d71 | Clostridiales | Lachnospiraceae | *Eubacterium ramulus* | 0.277 | 0.278 | 0.148 |
| *B. gallinaceum(OTU1)* | 213246e0ef70d6a711664e 7bd453dfe1c4e30106 | Bacteroidales | Bacteroidaceae | *Bacteroides gallinaceum(OTU1)* | 0.059 | 0.077 | 0.078 |
| *B. xylanolyticus* | e5db91f58295e8e2597c18 53ff383348b92c2d10 | Clostridiales | Lachnospiraceae | *Bacteroides xylanolyticus* | 0.199 | 0.135 | 0.062 |
| *B. producta(OTU81)* | 94b075dce1f20d6e7f9610 93eefe1a744fbac31c | Clostridiales | Lachnospiraceae | *Blautia producta(OTU81)* | 2.095 | 0.787 | 0.453 |
| Ruminococcaceae(OTU91) | dc838e370410fb0d47634 70f21542ac3f052167a | Clostridiales | Ruminococcaceae | Ruminococcaceae(OTU 91) | 0.145 | 0.083 | 0.050 |

| | | | | | | | |
|---|---|---|---|---|---|---|---|
| *P. olsenii* | 6d16c8dfa0d7dce3f6e27a904b61a21c2be8d0f5 | Clostridiales | Family XI | *Peptoniphilus olsenii* | 0.084 | 0.578 | 0.705 |
| *P. chartae(OTU69)* | 72d37863abe6d7f5bf4e982b8dbc3a0bb9a0ec6b | Bacteroidales | Tannerellaceae | *Parabacteroides chartae(OTU69)* | 1.071 | 1.326 | 0.562 |
| *O. denticanis* | af877f3cca95c2c63622bf43466341fb27fd9f9c | Bacteroidales | Marinifilaceae | *Odoribacter denticanis* | 0.163 | 0.084 | 0.208 |
| *I. bartlettii* | 22cb39282371a1bb23780ab7580e545a8662974f | Clostridiales | Peptostreptococcaceae | *Intestinibacter bartlettii* | 0.069 | 0.098 | 0.233 |
| Ruminococcaceae(OTU46) | 46c2c0eb93a1ac9fe10cf77d44de3ad899afab65 | Clostridiales | Ruminococcaceae | Ruminococcaceae(OTU46) | 0.898 | 0.496 | 0.303 |
| *L. fallax* | 5a61829814af31711bd597f814142ea04f651806 | Lactobacillales | Leuconostocaceae | *Leuconostoc fallax* | 0.003 | 0.034 | 0.101 |
| *Ruminiclostridium(OTU96)* | a4c6b012d72033d40026c27a32be2901526531ce | Clostridiales | Ruminococcaceae | *Ruminiclostridium(OTU96)Ruminiclostridium(OTU96)* | 0.559 | 0.093 | 0.051 |
| *Ezakiella(OTU94)* | 568373ca075c2cf3245a60cc7d25de80c9a90f43 | Clostridiales | Family XI | *Ezakiella(OTU94)* | 2.514 | 5.172 | 3.315 |
| *S. variabile(OTU93)* | cbf95306034ca33994a754185ac09707603d4a6e | Clostridiales | Ruminococcaceae | *Subdoligranulum variabile(OTU93)* | 0.794 | 0.408 | 0.204 |
| *R. gnavus* | 8adac8a13ae26336b44e1401350c58c3b7905593 | Clostridiales | Lachnospiraceae | *Ruminococcus gnavus* | 0.479 | 0.440 | 0.294 |
| *B. zoogleoformans* | 78d664d8199a1a5050c114fb3669304f3d9a8d4e | Bacteroidales | Bacteroidaceae | *Bacteroides zoogleoformans* | 0.154 | 0.035 | 0.071 |
| *F. prausnitzii(OTU80)* | 42d8f985e8c26f2b842279c4d945a4e78743b687 | Clostridiales | Ruminococcaceae | *Faecalibacterium prausnitzii(OTU80)* | 5.158 | 2.249 | 2.516 |

| | | | | | | | |
|---|---|---|---|---|---|---|---|
| *F. prausnitzii(OTU79)* | b57894411d721c03980b7 4f932b5d661aee5ddd1 | Clostridiales | Ruminococcaceae | *Faecalibacterium prausnitzii(OTU79)* | 0.286 | 0.194 | 0.046 |
| *B. caecicola(OTU78)* | 792f1a299b5d9bf02072d3 70497e5d2b18c6378a | Bacteroidales | Bacteroidaceae | *Bacteroides caecicola(OTU78)* | 0.624 | 0.388 | 0.336 |
| *H. effluvii(OTU77)* | 5cada88510f75d5e2aac52 d66708ccc43f8114fa | Clostridiales | Lachnospiraceae | *Hungatella effluvii(OTU77)* | 0.097 | 0.151 | 0.132 |
| *S5-A14a(OTU76)* | 58d72faf04d64350b1b667 4360ced8602b36eb94 | Clostridiales | Family XIII | *S5-A14a(OTU76)* | 0.027 | 0.272 | 0.301 |
| *A. ruminis(OTU75)* | 53e93035aec5d823c3e67e c85c1bb14945ca2050 | Clostridiales | Lachnospiraceae | *Agathobacter ruminis(OTU75)* | 3.648 | 1.421 | 1.833 |
| *E. peruensis* | c17c40fbf5843a9a0c17aa 02432b3902ffe021a4 | Clostridiales | Family XI | *Ezakiella peruensis* | 0.189 | 0.440 | 0.757 |
| *B. caecicola(OTU70)* | 67a04bc26f26894432770 b3cc66a7ba0b0ac3cab | Bacteroidales | Bacteroidaceae | *Bacteroides caecicola(OTU70)* | 0.043 | 0.052 | 0.077 |
| *B. coprosuis(OTU67)* | 6b7dc0529e91be865d421 1f5c3dfd65fc0b1b79d | Bacteroidales | Bacteroidaceae | *Bacteroides coprosuis(OTU67)* | 1.594 | 1.716 | 0.831 |
| *H. effluvii(OTU64)* | b301c74849d58d56977e3 bfb75fff7c930096735 | Clostridiales | Lachnospiraceae | *Hungatella effluvii(OTU64)* | 0.021 | 0.038 | 0.034 |
| *D. formicigenerans* | 832cb85976b1b559b471a 849211e41e9fc33e0d8 | Clostridiales | Lachnospiraceae | *Dorea formicigenerans* | 0.719 | 0.569 | 0.393 |
| *P. sp. 2007b* | 7ee6bdbeafe5f04b4d10a6 82bab889c7852be4cd | Bacteroidales | Porphyromonadaceae | *Porphyromonas sp. 2007b* | 0.025 | 0.947 | 0.964 |
| *P. buccalis* | 98cc48030d2670f449368 5c3bb1fbd7bf2182cc5 | Bacteroidales | Prevotellaceae | *Prevotella buccalis* | 0.235 | 1.172 | 1.395 |

| | | | | | | | |
|---|---|---|---|---|---|---|---|
| *P. ivorii* | 4055b3f1e3e0a8f04d0a83f7de0e7ca9fcdc0222 | Clostridiales | Family XI | *Peptoniphilus ivorii* | 0.147 | 1.631 | 1.077 |
| Lachnospiraceae(OTU55) | 671e184eef8fe77c8d41237140afb0c9774475d5 | Clostridiales | Lachnospiraceae | Lachnospiraceae(OTU55) | 0.252 | 0.087 | 0.051 |
| *P. chartae(OTU54)* | 18b62d123674facfbde7a0a670bdcbdeec61dca3 | Bacteroidales | Tannerellaceae | *Parabacteroides chartae(OTU54)* | 0.191 | 0.291 | 0.673 |
| *B. hydrogenotrophica* | b7958ee8f163b8c7f8efcd7f093f897421c2eeeb | Clostridiales | Lachnospiraceae | *Blautia hydrogenotrophica* | 1.025 | 0.206 | 0.194 |
| *E. rectale(OTU52)* | 7c2e0be05c81e92be893416502901cc796a11a97 | Clostridiales | Lachnospiraceae | *Eubacterium rectale(OTU52)* | 0.502 | 0.193 | 0.081 |
| *B. caecicola(OTU51)* | 0d181ab99c59f5465506730734e655becc2135a3 | Bacteroidales | Bacteroidaceae | *Bacteroides caecicola(OTU51)* | 2.774 | 1.804 | 0.792 |
| *W. ghanensis* | 2b6187b111e334718a1171a3395f8de7528eff3d | Lactobacillales | Leuconostocaceae | *Weissella ghanensis* | 0.016 | 0.045 | 0.128 |
| *E. rectale(OTU47)* | b21cc5dfad113e06eff6eeaefb9fb221379b1659 | Clostridiales | Lachnospiraceae | *Eubacterium rectale(OTU47)* | 0.185 | 0.074 | 0.101 |
| *S. variabile(OTU44)* | 5ecedc2ba1f718c8bf27df3525e075dcbc0f1819 | Clostridiales | Ruminococcaceae | *Subdoligranulum variabile(OTU44)* | 0.465 | 0.165 | 0.116 |
| *R. lactaris(OTU43)* | 7c8124a599ef55c3aca8d4c1edee81189d249762 | Clostridiales | Ruminococcaceae | *Ruminococcus lactaris(OTU43)* | 0.274 | 0.376 | 0.066 |
| *B. gallinaceum(OTU42)* | b83e3de5cc0156ec15810fb19eacc9c855a1f0c1 | Bacteroidales | Bacteroidaceae | *Bacteroides gallinaceum(OTU42)* | 0.066 | 0.069 | 0.035 |
| *B. propionicifaciens(OTU41)* | 23b75a012b2bec213e848893b24842d695014dee | Bacteroidales | Bacteroidaceae | *Bacteroides propionicifaciens(OTU41)* | 0.042 | 0.023 | 0.034 |

| | | | | | | | |
|---|---|---|---|---|---|---|---|
| *F. magna* | 36b71e0fa60b370e95082d0cee5ba3ed92ea4f14 | Clostridiales | Family XI | *Finegoldia magna* | 0.024 | 2.109 | 1.830 |
| *R. lactaris(OTU38)* | bf41a92ce95bc3c5068b7e5705d3eb75dd0441f1 | Clostridiales | Lachnospiraceae | *Ruminococcus lactaris(OTU38)* | 0.140 | 0.083 | 0.299 |
| *B. hansenii* | 3ba90965ce76880f6bbcddc96b6ab309ac932ee1 | Clostridiales | Lachnospiraceae | *Blautia hansenii* | 0.241 | 0.035 | 0.114 |
| *Ruminiclostridium(OTU36)* | 3bbf321699fe62c76c1931fff91d8e078feeba87 | Clostridiales | Ruminococcaceae | *Ruminiclostridium(OTU36)* | 0.059 | 0.027 | 0.069 |
| *C. comes* | 3ae4d87f6c2b8a813fb60f584165d9dd2821d2d5 | Clostridiales | Lachnospiraceae | *Coprococcus comes* | 0.688 | 0.398 | 0.143 |
| *P. lacrimalis* | 06a157c47ae3d15ba27d3409e47b2d1bda1228d9 | Clostridiales | Family XI | *Peptoniphilus lacrimalis* | 0.023 | 0.217 | 0.403 |
| *F. plautii* | 5f4f6d97004ada3407550a3afccd0f8b6bd61758 | Clostridiales | Ruminococcaceae | *Flavonifractor plautii* | 0.058 | 0.041 | 0.077 |
| *F. saccharivorans* | d77ce8b9e5a334c7e4103f6f21bf217540134ced | Clostridiales | Lachnospiraceae | *Fusicatenibacter saccharivorans* | 1.208 | 0.382 | 0.284 |
| *Bilophila(OTU30)* | 04f6d8267903b2cc853efc25678b5d11da252ba8 | Desulfovibrionales | Desulfovibrionaceae | *Bilophila(OTU30)* | 0.040 | 0.067 | 0.091 |
| *B. magnum* | 6c4a13cb8c2989b4ddf90ce4827e1a46ec08fdbd | Bifidobacteriales | Bifidobacteriaceae | *Bifidobacterium magnum* | 1.878 | 1.000 | 0.295 |
| *B. gallinaceum(OTU27)* | 833ee1018bc65e8c8dfd0e8c1a1ee5f25d4153d3 | Bacteroidales | Bacteroidaceae | *Bacteroides gallinaceum(OTU27)* | 0.155 | 0.243 | 0.139 |
| *A. indistinctus(OTU26)* | ebab44d6a7d001286bddcc959257cb9ac84ca9b4 | Bacteroidales | Rikenellaceae | *Alistipes indistinctus(OTU26)* | 0.222 | 0.091 | 0.072 |

| | | | | | | | |
|---|---|---|---|---|---|---|---|
| *L. pacaense* | 7f50d4d8d87d26b57bb3d a8edfb1b5d72f05e75c | Clostridiales | Lachnospiraceae | *Lachnoclostridium pacaense* | 0.020 | 0.049 | 0.134 |
| *B. coprophilus* | 055eb03b82092827561fa 816d990711e65a10a2d | Bacteroidales | Bacteroidaceae | *Bacteroides coprophilus* | 0.693 | 1.100 | 3.552 |
| Ruminococcaceae(OTU23) | 777b862accae329073ba4e 71b9b913d7922b771b | Clostridiales | Ruminococcaceae | Ruminococcaceae(OTU 23) | 0.365 | 0.268 | 0.069 |
| Ruminococcaceae(OTU22) | 71c28cdc242a9120ca397 0ba66b423c771079a9c | Clostridiales | Ruminococcaceae | Ruminococcaceae(OTU 22) | 0.161 | 0.113 | 0.071 |
| Lachnospiraceae(OTU18) | 3af07d551903d8f0450ac4 2e478b0858857f99b5 | Clostridiales | Lachnospiraceae | Lachnospiraceae(OTU1 8) | 0.220 | 0.155 | 0.060 |
| *E. oxidoreducens* | 6e7e6fb1a2a243f338a53d ddf99816665d92bd49 | Clostridiales | Lachnospiraceae | *Eubacterium oxidoreducens* | 0.825 | 0.229 | 0.155 |
| *B. caecicola(OTU16)* | e79e9f25d3414a038116ba 80ba9c90b4fea3e6d8 | Bacteroidales | Bacteroidaceae | *Bacteroides caecicola(OTU16)* | 0.049 | 0.050 | 0.095 |
| *B. sp. K4410.MGS-46* | 98896715b47780296c7b9 18cad24c28f4d058945 | Clostridiales | Ruminococcaceae | *Butyricicoccus sp. K4410.MGS-46* | 0.191 | 0.150 | 0.109 |
| *UBA1819(OTU14)* | 629b41b4214225423c5ff0 0fdbf5dfb2e2ba196d | Clostridiales | Ruminococcaceae | *UBA1819(OTU14)* | 0.089 | 0.032 | 0.244 |
| *C. amygdalinum* | 4cef6fc61a9c38eea885a1 8cde8de772f7412697 | Clostridiales | Lachnospiraceae | *Clostridium amygdalinum* | 0.009 | 0.031 | 0.018 |
| *S. thermophilus TH1435* | b5f49eebe9a619dafc2725 fd6b4b6db0a0476c8f | Lactobacillales | Streptococcaceae | *Streptococcus thermophilus TH1435* | 0.303 | 0.144 | 0.083 |
| *R. torques* | ef71fca08a960f52b392e3f 728c68d4779a06da8 | Clostridiales | Lachnospiraceae | *Ruminococcus torques* | 0.375 | 0.137 | 0.132 |

| | | | | | | | |
|---|---|---|---|---|---|---|---|
| *A. octavius* | 93a203cd147eaca4fefa76a477e453dd6a16f551 | Clostridiales | Family XI | *Anaerococcus octavius* | 0.010 | 0.943 | 1.549 |
| *B. caecicola(OTU10)* | 92d2b54526a297955ecf1d5be6e4d8695afa8172 | Bacteroidales | Bacteroidaceae | *Bacteroides caecicola(OTU10)* | 0.403 | 0.435 | 0.395 |
| *D. longicatena* | 0b814f25f0aac03c48bd8f719c6dba91f99df818 | Clostridiales | Lachnospiraceae | *Dorea longicatena* | 0.309 | 0.168 | 0.252 |
| *F. prausnitzii(OTU8)* | 7d217b5dac70526010aee823a23b9274e8fed212 | Clostridiales | Ruminococcaceae | *Faecalibacterium prausnitzii(OTU8)* | 2.616 | 1.918 | 1.618 |
| *O. valericigenes* | 759d3bce74029b0421b6b7829823329beec0cbda | Clostridiales | Ruminococcaceae | *Oscillibacter valericigenes* | 0.039 | 0.072 | 0.081 |
| *R. hominis* | 341e2fd8e931fa32697ef5a3cc97bbddddb81f32 | Clostridiales | Lachnospiraceae | *Roseburia hominis* | 0.260 | 0.164 | 0.184 |
| *P. secunda* | 595b5c69e0c0ab678b4b1457da6abbd3142afa3f | Betaproteobacteriales | Burkholderiaceae | *Parasutterella secunda* | 0.117 | 0.667 | 0.789 |
| *B. caecicola(OTU4)* | 2a069c745b4e3079b4be010d8e8d15866089a47c | Bacteroidales | Bacteroidaceae | *Bacteroides caecicola(OTU4)* | 8.597 | 11.336 | 12.917 |
| Ruminococcaceae(OTU3) | b0f51adf671d76a5f07c3b5bc16ab4d5c4ff609b | Clostridiales | Ruminococcaceae | Ruminococcaceae(OTU3) | 0.078 | 0.044 | 0.038 |
| *A. ruminis(OTU2)* | 543e9142bfd56e3312822277113e06cbb521aa91 | Clostridiales | Lachnospiraceae | *Agathobacter ruminis(OTU2)* | 0.463 | 0.104 | 0.142 |
| *R. lactaris(OTU0)* | 2725fb5dd2001597d9a103735ec633f4fec723af | Clostridiales | Lachnospiraceae | *Ruminococcus lactaris(OTU0)* | 0.269 | 0.234 | 0.173 |
| *B. caecicola(OTU72)* | 0cfa2d530d8bb8deb60488dd1a5ae30b0783b626 | Bacteroidales | Bacteroidaceae | *Bacteroides caecicola(OTU72)* | 0.290 | 0.098 | 0.043 |

| | | | | | | | |
|---|---|---|---|---|---|---|---|
| *B. gallinaceum(OTU83)* | d8d5f629119e6c50f9e7c9 5b6240649272c930cd | Bacteroidales | Bacteroidaceae | *Bacteroides gallinaceum(OTU83)* | 0.155 | 0.282 | 0.041 |
| *B. propionicifaciens(OTU29)* | 20218ad750875d16c9e56 42a9a5b42f360b4964d | Bacteroidales | Bacteroidaceae | *Bacteroides propionicifaciens(OTU 29)* | 0.439 | 0.371 | 0.200 |

Cont.

| Abb. | Differential abundance | | | | | |
|---|---|---|---|---|---|---|
| | obese_vs_normal_abun_pval | obese_vs_normal_abun_fdr | nash_vs_normal_abun_pval | nash_vs_normal_abun_fdr | nash_vs_obese_pval | nash_vs_obese_abun_fdr |
| *P. loveana* | 1.12E-01 | 2.46E-01 | 2.21E-03 | 1.07E-02 | 9.85E-02 | 7.72E-01 |
| *A. indistinctus(OTU57)* | 2.48E-02 | 8.90E-02 | 5.70E-03 | 2.05E-02 | 8.48E-01 | 9.68E-01 |
| *D. pneumosintes* | 7.53E-03 | 3.48E-02 | 5.42E-04 | 4.38E-03 | 9.63E-02 | 7.72E-01 |
| *A. caccae* | 2.75E-02 | 9.51E-02 | 1.80E-02 | 4.48E-02 | 7.33E-01 | 9.36E-01 |
| Lachnospiraceae(OTU85) | 8.73E-01 | 9.10E-01 | 3.29E-01 | 4.32E-01 | 4.24E-01 | 9.36E-01 |
| *B. barnesiae* | 6.88E-01 | 7.68E-01 | 1.93E-01 | 2.84E-01 | 3.01E-01 | 8.85E-01 |
| *Escherichia-Shigella(OTU88)* | 4.38E-01 | 5.99E-01 | 1.20E-02 | 3.52E-02 | 2.25E-02 | 7.72E-01 |
| *B. producta(OTU32)* | 1.34E-03 | 1.30E-02 | 3.76E-03 | 1.66E-02 | 8.31E-01 | 9.68E-01 |
| *F. prausnitzii(OTU68)* | 4.95E-02 | 1.50E-01 | 1.20E-02 | 3.52E-02 | 9.83E-01 | 9.83E-01 |
| *B. caecicola(OTU84)* | 3.23E-01 | 4.97E-01 | 5.15E-01 | 5.95E-01 | 7.98E-01 | 9.68E-01 |
| *B. coprosuis(OTU62)* | 1.57E-01 | 3.17E-01 | 1.93E-01 | 2.84E-01 | 9.83E-01 | 9.83E-01 |
| *B. caecicola(OTU86)* | 6.59E-01 | 7.68E-01 | 1.69E-01 | 2.65E-01 | 3.11E-01 | 8.88E-01 |
| *B. propionicifaciens(OTU73)* | 2.97E-01 | 4.73E-01 | 7.14E-03 | 2.39E-02 | 2.16E-01 | 8.85E-01 |
| *M. faecis* | 6.30E-01 | 7.55E-01 | 8.82E-01 | 9.01E-01 | 6.16E-01 | 9.36E-01 |
| *C. pinnipediorum* | 8.35E-04 | 1.16E-02 | 1.91E-03 | 1.07E-02 | 4.18E-01 | 9.36E-01 |
| *R. lituseburensis* | 3.43E-03 | 1.96E-02 | 1.21E-03 | 7.31E-03 | 5.02E-01 | 9.36E-01 |
| *A. indistinctus(OTU20)* | 9.57E-01 | 9.78E-01 | 2.61E-01 | 3.47E-01 | 8.81E-02 | 7.72E-01 |
| *C. catus* | 3.71E-01 | 5.53E-01 | 4.69E-01 | 5.61E-01 | 1.38E-01 | 7.90E-01 |

| | | | | | | |
|---|---|---|---|---|---|---|
| *E. ramulus* | 6.59E-01 | 7.68E-01 | 9.41E-01 | 9.51E-01 | 3.54E-01 | 9.27E-01 |
| *B. gallinaceum(OTU1)* | 8.41E-01 | 9.07E-01 | 7.34E-01 | 7.91E-01 | 4.62E-01 | 9.36E-01 |
| *B. xylanolyticus* | 6.88E-01 | 7.68E-01 | 7.61E-02 | 1.45E-01 | 1.17E-01 | 7.72E-01 |
| *B. producta(OTU81)* | 1.03E-02 | 4.34E-02 | 2.11E-03 | 1.07E-02 | 7.09E-01 | 9.36E-01 |
| Ruminococcaceae(OTU91) | 6.69E-01 | 7.68E-01 | 1.24E-01 | 2.11E-01 | 1.25E-01 | 7.72E-01 |
| *P. olsenii* | 3.01E-03 | 1.96E-02 | 1.95E-04 | 2.68E-03 | 3.37E-01 | 9.09E-01 |
| *P. chartae(OTU69)* | 4.30E-01 | 5.99E-01 | 5.06E-01 | 5.95E-01 | 8.23E-01 | 9.68E-01 |
| *O. denticanis* | 5.60E-02 | 1.55E-01 | 8.71E-01 | 8.99E-01 | 1.27E-01 | 7.72E-01 |
| *I. bartlettii* | 6.30E-01 | 7.55E-01 | 1.24E-01 | 2.11E-01 | 3.94E-01 | 9.36E-01 |
| Ruminococcaceae(OTU46) | 5.11E-02 | 1.50E-01 | 3.85E-02 | 8.49E-02 | 5.22E-01 | 9.36E-01 |
| *L. fallax* | 3.28E-03 | 1.96E-02 | 5.42E-04 | 4.38E-03 | 9.63E-02 | 7.72E-01 |
| *Ruminiclostridium(OTU96)* | 3.28E-03 | 1.96E-02 | 1.21E-04 | 2.35E-03 | 1.86E-01 | 8.85E-01 |
| *Ezakiella(OTU94)* | 1.47E-03 | 1.30E-02 | 7.50E-04 | 5.20E-03 | 3.01E-01 | 8.85E-01 |
| *S. variabile(OTU93)* | 5.95E-02 | 1.56E-01 | 3.94E-03 | 1.66E-02 | 2.91E-01 | 8.85E-01 |
| *R. gnavus* | 4.38E-01 | 5.99E-01 | 2.61E-01 | 3.47E-01 | 5.79E-01 | 9.36E-01 |
| *B. zoogleoformans* | 8.15E-03 | 3.59E-02 | 5.10E-02 | 1.05E-01 | 6.39E-01 | 9.36E-01 |
| *F. prausnitzii(OTU80)* | 6.95E-03 | 3.37E-02 | 1.20E-02 | 3.52E-02 | 9.32E-01 | 9.83E-01 |
| *F. prausnitzii(OTU79)* | 1.39E-02 | 5.63E-02 | 1.41E-02 | 3.81E-02 | 7.33E-01 | 9.36E-01 |
| *B. caecicola(OTU78)* | 8.31E-01 | 9.05E-01 | 6.36E-01 | 7.18E-01 | 5.36E-01 | 9.36E-01 |
| *H. effluvii(OTU77)* | 9.68E-01 | 9.78E-01 | 9.65E-01 | 9.65E-01 | 9.57E-01 | 9.83E-01 |
| *S5-A14a(OTU76)* | 6.41E-03 | 3.27E-02 | 6.53E-03 | 2.26E-02 | 8.98E-01 | 9.79E-01 |
| *A. ruminis(OTU75)* | 3.25E-02 | 1.09E-01 | 7.13E-02 | 1.38E-01 | 5.94E-01 | 9.36E-01 |

| | | | | | | |
|---|---|---|---|---|---|---|
| *E. peruensis* | 3.96E-02 | 1.28E-01 | 1.21E-04 | 2.35E-03 | 4.51E-02 | 7.72E-01 |
| *B. caecicola(OTU70)* | 8.73E-01 | 9.10E-01 | 2.43E-01 | 3.32E-01 | 1.15E-01 | 7.72E-01 |
| *B. coprosuis(OTU67)* | 3.57E-01 | 5.40E-01 | 2.37E-01 | 3.28E-01 | 7.01E-01 | 9.36E-01 |
| *H. effluvii(OTU64)* | 2.62E-01 | 4.42E-01 | 5.15E-01 | 5.95E-01 | 7.49E-01 | 9.44E-01 |
| *D. formicigenerans* | 8.47E-02 | 2.00E-01 | 4.13E-02 | 8.91E-02 | 8.48E-01 | 9.68E-01 |
| *P. sp. 2007b* | 1.47E-03 | 1.30E-02 | 4.54E-03 | 1.76E-02 | 5.58E-01 | 9.36E-01 |
| *P. buccalis* | 4.24E-03 | 2.29E-02 | 8.35E-04 | 5.40E-03 | 2.01E-01 | 8.85E-01 |
| *P. ivorii* | 3.25E-04 | 7.88E-03 | 3.45E-06 | 1.68E-04 | 9.32E-01 | 9.83E-01 |
| Lachnospiraceae(OTU55) | 1.74E-02 | 6.48E-02 | 4.54E-03 | 1.76E-02 | 8.98E-01 | 9.79E-01 |
| *P. chartae(OTU54)* | 5.75E-01 | 7.15E-01 | 5.74E-01 | 6.55E-01 | 7.33E-01 | 9.36E-01 |
| *B. hydrogenotrophica* | 4.19E-04 | 8.13E-03 | 1.54E-04 | 2.49E-03 | 7.09E-01 | 9.36E-01 |
| *E. rectale(OTU52)* | 2.64E-03 | 1.96E-02 | 2.61E-04 | 2.68E-03 | 3.32E-01 | 9.09E-01 |
| *B. caecicola(OTU51)* | 1.31E-01 | 2.82E-01 | 2.83E-03 | 1.31E-02 | 1.53E-01 | 8.25E-01 |
| *W. ghanensis* | 1.06E-01 | 2.39E-01 | 1.25E-02 | 3.56E-02 | 2.28E-01 | 8.85E-01 |
| *E. rectale(OTU47)* | 1.81E-01 | 3.45E-01 | 8.64E-02 | 1.61E-01 | 4.69E-01 | 9.36E-01 |
| *S. variabile(OTU44)* | 4.22E-02 | 1.32E-01 | 3.85E-02 | 8.49E-02 | 5.94E-01 | 9.36E-01 |
| *R. lactaris(OTU43)* | 5.57E-01 | 7.01E-01 | 1.53E-02 | 4.02E-02 | 4.07E-02 | 7.72E-01 |
| *B. gallinaceum(OTU42)* | 6.12E-01 | 7.51E-01 | 1.65E-01 | 2.62E-01 | 3.71E-01 | 9.36E-01 |
| *B. propionicifaciens(OTU41)* | 4.15E-01 | 5.99E-01 | 7.79E-01 | 8.21E-01 | 2.63E-01 | 8.85E-01 |
| *F. magna* | 4.33E-05 | 4.20E-03 | 1.21E-04 | 2.35E-03 | 8.81E-01 | 9.79E-01 |
| *R. lactaris(OTU38)* | 1.00E+00 | 1.00E+00 | 2.04E-01 | 2.95E-01 | 1.10E-01 | 7.72E-01 |
| *B. hansenii* | 2.93E-04 | 7.88E-03 | 5.70E-03 | 2.05E-02 | 8.23E-01 | 9.68E-01 |

| | | | | | | |
|---|---|---|---|---|---|---|
| *Ruminiclostridium(OTU36)* | 6.14E-02 | 1.57E-01 | 3.75E-01 | 4.66E-01 | 5.79E-01 | 9.36E-01 |
| *C. comes* | 1.49E-01 | 3.14E-01 | 1.80E-02 | 4.48E-02 | 2.77E-01 | 8.85E-01 |
| *P. lacrimalis* | 2.31E-03 | 1.87E-02 | 7.50E-04 | 5.20E-03 | 1.05E-01 | 7.72E-01 |
| *F. plautii* | 4.38E-01 | 5.99E-01 | 7.23E-01 | 7.88E-01 | 6.39E-01 | 9.36E-01 |
| *F. saccharivorans* | 8.35E-04 | 1.16E-02 | 2.46E-04 | 2.68E-03 | 6.31E-01 | 9.36E-01 |
| *Bilophila(OTU30)* | 4.71E-01 | 6.17E-01 | 1.83E-01 | 2.78E-01 | 4.30E-01 | 9.36E-01 |
| *B. magnum* | 5.43E-02 | 1.55E-01 | 2.00E-03 | 1.07E-02 | 1.20E-01 | 7.72E-01 |
| *B. gallinaceum(OTU27)* | 2.73E-01 | 4.42E-01 | 1.32E-01 | 2.16E-01 | 5.09E-01 | 9.36E-01 |
| *A. indistinctus(OTU26)* | 2.62E-01 | 4.42E-01 | 1.04E-01 | 1.87E-01 | 4.43E-01 | 9.36E-01 |
| *L. pacaense* | 8.94E-01 | 9.22E-01 | 6.25E-02 | 1.24E-01 | 9.21E-02 | 7.72E-01 |
| *B. coprophilus* | 4.71E-01 | 6.17E-01 | 8.36E-01 | 8.72E-01 | 6.24E-01 | 9.36E-01 |
| Ruminococcaceae(OTU23) | 1.53E-01 | 3.15E-01 | 1.36E-02 | 3.76E-02 | 2.63E-01 | 8.85E-01 |
| Ruminococcaceae(OTU22) | 4.71E-01 | 6.17E-01 | 9.19E-02 | 1.68E-01 | 1.93E-01 | 8.85E-01 |
| Lachnospiraceae(OTU18) | 6.79E-01 | 7.68E-01 | 1.14E-01 | 2.01E-01 | 2.63E-01 | 8.85E-01 |
| *E. oxidoreducens* | 1.01E-03 | 1.23E-02 | 2.76E-04 | 2.68E-03 | 6.31E-01 | 9.36E-01 |
| *B. caecicola(OTU16)* | 8.52E-01 | 9.08E-01 | 1.32E-01 | 2.16E-01 | 1.07E-01 | 7.72E-01 |
| *B. sp. K4410.MGS-46* | 2.67E-01 | 4.42E-01 | 1.74E-01 | 2.68E-01 | 7.33E-01 | 9.36E-01 |
| *UBA1819(OTU14)* | 2.00E-01 | 3.65E-01 | 3.75E-01 | 4.66E-01 | 9.83E-01 | 9.83E-01 |
| *C. amygdalinum* | 1.73E-01 | 3.35E-01 | 4.25E-01 | 5.21E-01 | 4.69E-01 | 9.36E-01 |
| *S. thermophilus TH1435* | 8.24E-02 | 2.00E-01 | 3.33E-02 | 7.69E-02 | 7.65E-01 | 9.52E-01 |
| *R. torques* | 1.95E-01 | 3.64E-01 | 2.09E-01 | 2.98E-01 | 9.41E-01 | 9.83E-01 |
| *A. octavius* | 1.93E-04 | 7.88E-03 | 1.80E-06 | 1.68E-04 | 2.72E-01 | 8.85E-01 |

| | | | | | | |
|---|---|---|---|---|---|---|
| *B. caecicola(OTU10)* | 3.78E-01 | 5.55E-01 | 7.01E-01 | 7.72E-01 | 6.93E-01 | 9.36E-01 |
| *D. longicatena* | 2.09E-01 | 3.76E-01 | 6.79E-01 | 7.57E-01 | 4.30E-01 | 9.36E-01 |
| *F. prausnitzii(OTU8)* | 5.78E-02 | 1.56E-01 | 2.37E-01 | 3.28E-01 | 7.17E-01 | 9.36E-01 |
| *O. valericigenes* | 5.21E-01 | 6.65E-01 | 7.67E-01 | 8.18E-01 | 8.65E-01 | 9.75E-01 |
| *R. hominis* | 2.73E-01 | 4.42E-01 | 6.25E-02 | 1.24E-01 | 6.54E-01 | 9.36E-01 |
| *P. secunda* | 4.87E-01 | 6.30E-01 | 3.52E-01 | 4.55E-01 | 7.98E-01 | 9.68E-01 |
| *B. caecicola(OTU4)* | 7.48E-01 | 8.25E-01 | 4.60E-01 | 5.58E-01 | 2.96E-01 | 8.85E-01 |
| Ruminococcaceae(OTU3) | 8.00E-02 | 1.99E-01 | 3.09E-02 | 7.31E-02 | 5.65E-01 | 9.36E-01 |
| *A. ruminis(OTU2)* | 1.62E-02 | 6.27E-02 | 2.87E-02 | 6.95E-02 | 9.49E-01 | 9.83E-01 |
| *R. lactaris(OTU0)* | 1.73E-01 | 3.35E-01 | 3.75E-01 | 4.66E-01 | 5.02E-01 | 9.36E-01 |
| *B. caecicola(OTU72)* | 1.03E-01 | 2.38E-01 | 8.50E-03 | 2.75E-02 | 4.56E-01 | 9.36E-01 |
| *B. gallinaceum(OTU83)* | 3.10E-01 | 4.85E-01 | 5.10E-02 | 1.05E-01 | 2.68E-01 | 8.85E-01 |
| *B. propionicifaciens(OTU29)* | 2.19E-01 | 3.86E-01 | 1.39E-01 | 2.25E-01 | 6.24E-01 | 9.36E-01 |

Cont.

| Abb. | HITS score | | | | | |
|---|---|---|---|---|---|---|
| | normal_hits_score | normal_hits_pval | obese_hits_score | obese_hits_pval | nash_hits_score | nash_hits_pval |
| *P. loveana* | 0.033 | 1.30E-02 | 0.005 | 9.29E-01 | 0.017 | 8.30E-02 |
| *A. indistinctus(OTU57)* | 0.003 | 8.25E-01 | 0.004 | 9.62E-01 | 0.033 | 1.00E-03 |
| *D. pneumosintes* | 0.002 | 8.52E-01 | 0.034 | 1.00E-03 | 0.025 | 5.00E-03 |
| *A. caccae* | 0.036 | 1.60E-02 | 0.037 | 1.00E-03 | 0.008 | 6.64E-01 |
| Lachnospiraceae(OTU85) | 0.038 | 1.40E-02 | 0.005 | 9.33E-01 | 0.007 | 7.56E-01 |
| *B. barnesiae* | 0.001 | 9.08E-01 | 0.012 | 3.02E-01 | 0.020 | 2.40E-02 |
| *Escherichia-Shigella(OTU88)* | 0.002 | 8.56E-01 | 0.000 | 1.00E+00 | 0.002 | 9.77E-01 |
| *B. producta(OTU32)* | 0.022 | 1.12E-01 | 0.035 | 1.00E-03 | 0.009 | 6.24E-01 |
| *F. prausnitzii(OTU68)* | 0.000 | 1.00E+00 | 0.014 | 1.53E-01 | 0.004 | 9.53E-01 |
| *B. caecicola(OTU84)* | 0.012 | 3.07E-01 | 0.003 | 9.92E-01 | 0.004 | 9.14E-01 |
| *B. coprosuis(OTU62)* | 0.007 | 5.53E-01 | 0.004 | 9.67E-01 | 0.006 | 8.23E-01 |
| *B. caecicola(OTU86)* | 0.001 | 9.15E-01 | 0.008 | 7.34E-01 | 0.027 | 4.00E-03 |
| *B. propionicifaciens(OTU73)* | 0.005 | 6.92E-01 | 0.009 | 5.87E-01 | 0.027 | 2.00E-03 |
| *M. faecis* | 0.001 | 9.02E-01 | 0.016 | 9.40E-02 | 0.001 | 9.99E-01 |
| *C. pinnipediorum* | 0.006 | 5.82E-01 | 0.003 | 9.86E-01 | 0.000 | 1.00E+00 |
| *R. lituseburensis* | 0.004 | 7.52E-01 | 0.002 | 9.97E-01 | 0.004 | 9.35E-01 |
| *A. indistinctus(OTU20)* | 0.003 | 8.22E-01 | 0.005 | 9.47E-01 | 0.000 | 1.00E+00 |
| *C. catus* | 0.002 | 8.45E-01 | 0.007 | 7.65E-01 | 0.004 | 9.60E-01 |
| *E. ramulus* | 0.002 | 8.55E-01 | 0.025 | 2.00E-03 | 0.019 | 3.50E-02 |
| *B. gallinaceum(OTU1)* | 0.002 | 8.54E-01 | 0.000 | 1.00E+00 | 0.001 | 9.99E-01 |
| *B. xylanolyticus* | 0.005 | 6.46E-01 | 0.001 | 9.99E-01 | 0.003 | 9.78E-01 |
| *B. producta(OTU81)* | 0.022 | 8.70E-02 | 0.032 | 1.00E-03 | 0.008 | 6.44E-01 |

| | | | | | | |
|---|---|---|---|---|---|---|
| Ruminococcaceae(OTU91) | 0.002 | 8.92E-01 | 0.023 | 7.00E-03 | 0.001 | 9.95E-01 |
| *P. olsenii* | 0.012 | 3.40E-01 | 0.018 | 4.20E-02 | 0.024 | 7.00E-03 |
| *P. chartae(OTU69)* | 0.001 | 9.31E-01 | 0.005 | 9.17E-01 | 0.001 | 9.99E-01 |
| *O. denticanis* | 0.000 | 9.50E-01 | 0.004 | 9.62E-01 | 0.005 | 8.65E-01 |
| *I. bartlettii* | 0.000 | 9.43E-01 | 0.003 | 9.80E-01 | 0.000 | 1.00E+00 |
| Ruminococcaceae(OTU46) | 0.000 | 9.44E-01 | 0.003 | 9.68E-01 | 0.003 | 9.71E-01 |
| *L. fallax* | 0.000 | 9.48E-01 | 0.000 | 1.00E+00 | 0.000 | 1.00E+00 |
| *Ruminiclostridium(OTU96)* | 0.000 | 9.42E-01 | 0.005 | 8.91E-01 | 0.016 | 1.04E-01 |
| *Ezakiella(OTU94)* | 0.010 | 4.07E-01 | 0.019 | 3.20E-02 | 0.018 | 5.50E-02 |
| *S. variabile(OTU93)* | 0.002 | 8.83E-01 | 0.012 | 3.32E-01 | 0.003 | 9.73E-01 |
| *R. gnavus* | 0.000 | 9.50E-01 | 0.034 | 1.00E-03 | 0.003 | 9.87E-01 |
| *B. zoogleoformans* | 0.002 | 8.62E-01 | 0.013 | 2.73E-01 | 0.006 | 8.78E-01 |
| *F. prausnitzii(OTU80)* | 0.062 | 1.00E-03 | 0.032 | 1.00E-03 | 0.011 | 3.86E-01 |
| *F. prausnitzii(OTU79)* | 0.000 | 1.00E+00 | 0.009 | 5.76E-01 | 0.000 | 1.00E+00 |
| *B. caecicola(OTU78)* | 0.000 | 9.59E-01 | 0.001 | 1.00E+00 | 0.019 | 4.50E-02 |
| *H. effluvii(OTU77)* | 0.000 | 1.00E+00 | 0.011 | 3.97E-01 | 0.010 | 4.69E-01 |
| *S5-A14a(OTU76)* | 0.037 | 1.10E-02 | 0.003 | 9.93E-01 | 0.009 | 6.08E-01 |
| *A. ruminis(OTU75)* | 0.072 | 1.00E-03 | 0.010 | 5.16E-01 | 0.015 | 1.65E-01 |
| *E. peruensis* | 0.021 | 1.21E-01 | 0.000 | 1.00E+00 | 0.038 | 1.00E-03 |
| *B. caecicola(OTU70)* | 0.002 | 8.92E-01 | 0.013 | 2.37E-01 | 0.012 | 2.85E-01 |
| *B. coprosuis(OTU67)* | 0.001 | 9.27E-01 | 0.000 | 1.00E+00 | 0.006 | 8.13E-01 |
| *H. effluvii(OTU64)* | 0.000 | 1.00E+00 | 0.007 | 7.91E-01 | 0.025 | 9.00E-03 |
| *D. formicigenerans* | 0.000 | 9.52E-01 | 0.030 | 1.00E-03 | 0.017 | 8.10E-02 |
| *P. sp. 2007b* | 0.002 | 8.67E-01 | 0.021 | 1.60E-02 | 0.033 | 2.00E-03 |
| *P. buccalis* | 0.011 | 3.54E-01 | 0.042 | 1.00E-03 | 0.006 | 8.27E-01 |

| | | | | | | |
|---|---|---|---|---|---|---|
| *P. ivorii* | 0.052 | 5.00E-03 | 0.008 | 7.40E-01 | 0.029 | 1.00E-03 |
| Lachnospiraceae(OTU55) | 0.038 | 9.00E-03 | 0.001 | 1.00E+00 | 0.012 | 3.14E-01 |
| *P. chartae(OTU54)* | 0.002 | 8.25E-01 | 0.004 | 9.56E-01 | 0.009 | 6.00E-01 |
| *B. hydrogenotrophica* | 0.011 | 3.74E-01 | 0.019 | 3.10E-02 | 0.008 | 6.86E-01 |
| *E. rectale(OTU52)* | 0.029 | 5.40E-02 | 0.023 | 1.20E-02 | 0.003 | 9.61E-01 |
| *B. caecicola(OTU51)* | 0.015 | 2.52E-01 | 0.009 | 5.91E-01 | 0.016 | 1.25E-01 |
| *W. ghanensis* | 0.000 | 1.00E+00 | 0.001 | 1.00E+00 | 0.000 | 1.00E+00 |
| *E. rectale(OTU47)* | 0.028 | 5.80E-02 | 0.008 | 6.73E-01 | 0.002 | 9.85E-01 |
| *S. variabile(OTU44)* | 0.007 | 5.65E-01 | 0.010 | 4.51E-01 | 0.010 | 5.30E-01 |
| *R. lactaris(OTU43)* | 0.000 | 1.00E+00 | 0.008 | 6.90E-01 | 0.000 | 9.95E-01 |
| *B. gallinaceum(OTU42)* | 0.024 | 1.12E-01 | 0.003 | 9.77E-01 | 0.008 | 6.62E-01 |
| *B. propionicifaciens(OTU41)* | 0.001 | 9.15E-01 | 0.001 | 9.99E-01 | 0.023 | 1.10E-02 |
| *F. magna* | 0.046 | 6.00E-03 | 0.003 | 9.85E-01 | 0.010 | 4.91E-01 |
| *R. lactaris(OTU38)* | 0.006 | 6.38E-01 | 0.009 | 6.54E-01 | 0.005 | 8.93E-01 |
| *B. hansenii* | 0.038 | 1.20E-02 | 0.029 | 1.00E-03 | 0.005 | 8.88E-01 |
| *Ruminiclostridium(OTU36)* | 0.003 | 8.17E-01 | 0.001 | 9.99E-01 | 0.000 | 9.99E-01 |
| *C. comes* | 0.001 | 8.96E-01 | 0.028 | 1.00E-03 | 0.008 | 6.59E-01 |
| *P. lacrimalis* | 0.003 | 7.60E-01 | 0.038 | 1.00E-03 | 0.038 | 1.00E-03 |
| *F. plautii* | 0.000 | 9.40E-01 | 0.016 | 9.00E-02 | 0.006 | 8.31E-01 |
| *F. saccharivorans* | 0.017 | 2.03E-01 | 0.006 | 8.36E-01 | 0.014 | 1.94E-01 |
| *Bilophila(OTU30)* | 0.002 | 8.84E-01 | 0.000 | 1.00E+00 | 0.001 | 9.99E-01 |
| *B. magnum* | 0.000 | 1.00E+00 | 0.008 | 6.82E-01 | 0.003 | 9.60E-01 |
| *B. gallinaceum(OTU27)* | 0.046 | 2.00E-03 | 0.002 | 9.88E-01 | 0.011 | 3.96E-01 |
| *A. indistinctus(OTU26)* | 0.000 | 1.00E+00 | 0.001 | 1.00E+00 | 0.001 | 9.99E-01 |
| *L. pacaense* | 0.011 | 4.07E-01 | 0.000 | 9.99E-01 | 0.002 | 9.90E-01 |

| | | | | | | |
|---|---|---|---|---|---|---|
| *B. coprophilus* | 0.000 | 9.56E-01 | 0.000 | 1.00E+00 | 0.000 | 1.00E+00 |
| Ruminococcaceae(OTU23) | 0.002 | 8.81E-01 | 0.000 | 1.00E+00 | 0.003 | 9.82E-01 |
| Ruminococcaceae(OTU22) | 0.000 | 1.00E+00 | 0.001 | 9.96E-01 | 0.000 | 9.97E-01 |
| Lachnospiraceae(OTU18) | 0.000 | 1.00E+00 | 0.020 | 2.20E-02 | 0.007 | 7.56E-01 |
| *E. oxidoreducens* | 0.003 | 8.00E-01 | 0.005 | 9.30E-01 | 0.006 | 8.48E-01 |
| *B. caecicola(OTU16)* | 0.001 | 9.09E-01 | 0.002 | 9.98E-01 | 0.012 | 3.13E-01 |
| *B. sp. K4410.MGS-46* | 0.002 | 8.75E-01 | 0.033 | 1.00E-03 | 0.001 | 9.94E-01 |
| *UBA1819(OTU14)* | 0.000 | 1.00E+00 | 0.001 | 1.00E+00 | 0.009 | 5.32E-01 |
| *C. amygdalinum* | 0.012 | 3.30E-01 | 0.009 | 5.70E-01 | 0.000 | 1.00E+00 |
| *S. thermophilus TH1435* | 0.004 | 7.47E-01 | 0.012 | 3.34E-01 | 0.003 | 9.75E-01 |
| *R. torques* | 0.001 | 9.14E-01 | 0.018 | 4.20E-02 | 0.006 | 8.42E-01 |
| *A. octavius* | 0.021 | 1.64E-01 | 0.001 | 9.97E-01 | 0.009 | 5.97E-01 |
| *B. caecicola(OTU10)* | 0.002 | 8.86E-01 | 0.003 | 9.85E-01 | 0.030 | 2.00E-03 |
| *D. longicatena* | 0.002 | 8.48E-01 | 0.016 | 9.70E-02 | 0.006 | 8.55E-01 |
| *F. prausnitzii(OTU8)* | 0.000 | 9.36E-01 | 0.012 | 3.41E-01 | 0.010 | 5.06E-01 |
| *O. valericigenes* | 0.000 | 1.00E+00 | 0.001 | 9.99E-01 | 0.005 | 9.08E-01 |
| *R. hominis* | 0.002 | 8.61E-01 | 0.006 | 8.59E-01 | 0.001 | 9.97E-01 |
| *P. secunda* | 0.000 | 9.36E-01 | 0.002 | 9.97E-01 | 0.033 | 1.00E-03 |
| *B. caecicola(OTU4)* | 0.002 | 8.69E-01 | 0.005 | 8.96E-01 | 0.012 | 3.52E-01 |
| Ruminococcaceae(OTU3) | 0.019 | 1.32E-01 | 0.007 | 7.40E-01 | 0.005 | 8.89E-01 |
| *A. ruminis(OTU2)* | 0.065 | 1.00E-03 | 0.008 | 6.69E-01 | 0.012 | 3.18E-01 |
| *R. lactaris(OTU0)* | 0.000 | 1.00E+00 | 0.000 | 1.00E+00 | 0.000 | 1.00E+00 |
| *B. caecicola(OTU72)* | 0.004 | 7.65E-01 | 0.012 | 3.08E-01 | 0.021 | 2.20E-02 |
| *B. gallinaceum(OTU83)* | 0.022 | 1.04E-01 | 0.006 | 8.57E-01 | 0.036 | 1.00E-03 |
| *B. propionicifaciens(OTU29)* | 0.011 | 3.65E-01 | 0.005 | 9.48E-01 | 0.042 | 1.00E-03 |

Cont.

| Abb. | Intervention score | | | | | |
|---|---|---|---|---|---|---|
| | obese_intervention_score | obese_cumulative_intervention_score | obese_cumulative_intervention_rank | nash_intervention_score | nash_cumulative_intervention_score | nash_cumulative_intervention_rank |
| *P. loveana* | 0.059 | 0.963 | 47 | 0.346 | 0.346 | 1 |
| *A. indistinctus(OTU57)* | 0.022 | 0.963 | 53 | 0.206 | 0.552 | 2 |
| *D. pneumosintes* | 0.279 | 0.963 | 59 | 0.294 | 0.664 | 3 |
| *A. caccae* | 0.281 | 0.963 | 57 | 0.042 | 0.664 | 4 |
| Lachnospiraceae(OTU85) | 0.025 | 0.963 | 37 | -0.039 | 0.740 | 5 |
| *B. barnesiae* | -0.156 | 0.776 | 7 | 0.111 | 0.809 | 6 |
| *Escherichia-Shigella(OTU88)* | 0.002 | 0.952 | 19 | 0.061 | 0.870 | 7 |
| *B. producta(OTU32)* | 0.256 | 0.963 | 70 | 0.196 | 0.923 | 8 |
| *F. prausnitzii(OTU68)* | 0.104 | 0.963 | 46 | 0.012 | 0.935 | 9 |
| *B. caecicola(OTU84)* | -0.022 | 0.963 | 38 | 0.011 | 0.947 | 10 |
| *B. coprosuis(OTU62)* | 0.015 | 0.963 | 50 | 0.079 | 0.947 | 11 |
| *B. caecicola(OTU86)* | -0.013 | 0.963 | 36 | -0.131 | 0.954 | 12 |
| *B. propionicifaciens(OTU73)* | -0.011 | 0.962 | 26 | 0.171 | 0.966 | 13 |
| *M. faecis* | -0.080 | 0.942 | 16 | 0.009 | 0.974 | 14 |

| | | | | | | |
|---|---|---|---|---|---|---|
| *C. pinnipediorum* | 0.031 | 0.915 | 12 | 0.006 | 0.981 | 15 |
| *R. lituseburensis* | 0.008 | 0.947 | 17 | 0.095 | 0.984 | 16 |
| *A. indistinctus(OTU20)* | 0.016 | 0.916 | 79 | 0.003 | 0.987 | 17 |
| *C. catus* | 0.079 | 0.963 | 51 | -0.003 | 0.989 | 18 |
| *E. ramulus* | -0.112 | 0.963 | 40 | 0.182 | 0.991 | 19 |
| *B. gallinaceum(OTU1)* | 0.002 | 0.956 | 21 | -0.015 | 0.993 | 20 |
| *B. xylanolyticus* | 0.011 | 0.963 | 52 | 0.003 | 0.995 | 21 |
| *B. producta(OTU81)* | 0.201 | 0.963 | 41 | 0.157 | 0.995 | 22 |
| Ruminococcaceae(OTU91) | 0.178 | 0.935 | 15 | -0.021 | 0.996 | 23 |
| *P. olsenii* | 0.149 | 0.817 | 8 | 0.084 | 0.998 | 24 |
| *P. chartae(OTU69)* | -0.030 | 0.963 | 45 | 0.001 | 0.999 | 25 |
| *O. denticanis* | 0.109 | 0.958 | 22 | -0.014 | 0.999 | 26 |
| *I. bartlettii* | -0.106 | 0.963 | 28 | -0.007 | 1.000 | 27 |
| Ruminococcaceae(OTU46) | -0.095 | 0.963 | 58 | 0.003 | 1.000 | 28 |
| *L. fallax* | 0.000 | 0.963 | 64 | 0.000 | 1.000 | 29 |
| *Ruminiclostridium(OTU96)* | 0.026 | 0.963 | 32 | 0.050 | 1.000 | 30 |
| *Ezakiella(OTU94)* | 0.146 | 0.963 | 33 | 0.181 | 1.000 | 31 |
| *S. variabile(OTU93)* | 0.132 | 0.934 | 14 | 0.004 | 1.000 | 32 |
| *R. gnavus* | 0.249 | 0.963 | 34 | 0.034 | 1.000 | 33 |
| *B. zoogleoformans* | 0.089 | 0.647 | 4 | 0.076 | 1.000 | 34 |

| | | | | | | |
|---|---|---|---|---|---|---|
| *F. prausnitzii(OTU80)* | 0.242 | 0.963 | 42 | 0.200 | 1.000 | 35 |
| *F. prausnitzii(OTU79)* | 0.092 | 0.880 | 10 | 0.000 | 1.000 | 36 |
| *B. caecicola(OTU78)* | 0.004 | 0.963 | 29 | 0.122 | 1.000 | 37 |
| *H. effluvii(OTU77)* | -0.029 | 0.963 | 43 | -0.035 | 1.000 | 38 |
| *S5-A14a(OTU76)* | 0.074 | 0.745 | 6 | 0.162 | 1.000 | 39 |
| *A. ruminis(OTU75)* | 0.238 | 0.475 | 2 | 0.254 | 1.000 | 40 |
| *E. peruensis* | 0.074 | 0.963 | 44 | 0.247 | 1.000 | 41 |
| *B. caecicola(OTU70)* | -0.012 | 0.854 | 9 | -0.020 | 1.000 | 42 |
| *B. coprosuis(OTU67)* | 0.003 | 0.963 | 27 | 0.010 | 1.000 | 43 |
| *H. effluvii(OTU64)* | -0.074 | 0.963 | 48 | -0.125 | 1.000 | 44 |
| *D. formicigenerans* | 0.237 | 0.963 | 49 | 0.231 | 1.000 | 45 |
| *P. sp. 2007b* | 0.240 | 0.240 | 1 | 0.376 | 1.000 | 46 |
| *P. buccalis* | 0.397 | 0.963 | 35 | 0.174 | 1.000 | 47 |
| *P. ivorii* | 0.186 | 0.587 | 3 | 0.136 | 1.000 | 48 |
| Lachnospiraceae(OTU55) | 0.121 | 0.926 | 13 | 0.073 | 1.000 | 49 |
| *P. chartae(OTU54)* | -0.003 | 0.963 | 30 | -0.033 | 1.000 | 50 |
| *B. hydrogenotrophica* | 0.138 | 0.963 | 54 | 0.100 | 1.000 | 51 |
| *E. rectale(OTU52)* | 0.153 | 0.963 | 55 | 0.139 | 1.000 | 52 |
| *B. caecicola(OTU51)* | -0.008 | 0.963 | 56 | 0.075 | 1.000 | 53 |
| *W. ghanensis* | 0.021 | 0.963 | 31 | -0.002 | 1.000 | 54 |

| | | | | | | |
|---|---|---|---|---|---|---|
| *E. rectale(OTU47)* | 0.189 | 0.916 | 95 | 0.061 | 1.000 | 55 |
| *S. variabile(OTU44)* | 0.121 | 0.963 | 60 | 0.018 | 1.000 | 56 |
| *R. lactaris(OTU43)* | -0.237 | 0.963 | 61 | -0.001 | 1.000 | 57 |
| *B. gallinaceum(OTU42)* | -0.006 | 0.903 | 11 | 0.076 | 1.000 | 58 |
| *B. propionicifaciens(OTU41)* | 0.010 | 0.963 | 62 | 0.107 | 1.000 | 59 |
| *F. magna* | 0.134 | 0.963 | 63 | 0.129 | 1.000 | 60 |
| *R. lactaris(OTU38)* | 0.013 | 0.963 | 65 | 0.090 | 1.000 | 61 |
| *B. hansenii* | 0.284 | 0.963 | 66 | 0.116 | 1.000 | 62 |
| *Ruminiclostridium(OTU36)* | 0.013 | 0.950 | 18 | 0.005 | 1.000 | 63 |
| *C. comes* | 0.202 | 0.963 | 67 | 0.101 | 1.000 | 64 |
| *P. lacrimalis* | 0.191 | 0.963 | 68 | 0.289 | 1.000 | 65 |
| *F. plautii* | 0.156 | 0.963 | 69 | 0.050 | 1.000 | 66 |
| *F. saccharivorans* | 0.127 | 0.963 | 71 | 0.238 | 1.000 | 67 |
| *Bilophila(OTU30)* | 0.002 | 0.954 | 20 | -0.023 | 1.000 | 68 |
| *B. magnum* | 0.104 | 0.963 | 72 | 0.135 | 1.000 | 69 |
| *B. gallinaceum(OTU27)* | 0.027 | 0.961 | 25 | 0.104 | 1.000 | 70 |
| *A. indistinctus(OTU26)* | -0.014 | 0.916 | 74 | 0.021 | 1.000 | 71 |
| *L. pacaense* | 0.048 | 0.916 | 75 | 0.011 | 1.000 | 72 |
| *B. coprophilus* | -0.005 | 0.916 | 76 | 0.000 | 1.000 | 73 |

| | | | | | | |
|---|---|---|---|---|---|---|
| Ruminococcaceae(OTU23) | 0.002 | 0.959 | 23 | 0.005 | 1.000 | 74 |
| Ruminococcaceae(OTU22) | 0.004 | 0.916 | 77 | 0.006 | 1.000 | 75 |
| Lachnospiraceae(OTU18) | 0.174 | 0.916 | 80 | 0.115 | 1.000 | 76 |
| *E. oxidoreducens* | 0.084 | 0.916 | 81 | 0.154 | 1.000 | 77 |
| *B. caecicola(OTU16)* | -0.001 | 0.961 | 24 | -0.015 | 1.000 | 78 |
| *B. sp. K4410.MGS-46* | 0.165 | 0.916 | 82 | 0.040 | 1.000 | 79 |
| *UBA1819(OTU14)* | -0.007 | 0.903 | 96 | 0.192 | 1.000 | 80 |
| *C. amygdalinum* | 0.028 | 0.708 | 5 | 0.012 | 1.000 | 81 |
| *S. thermophilus TH1435* | 0.269 | 0.916 | 78 | 0.035 | 0.923 | 82 |
| *R. torques* | 0.288 | 0.916 | 83 | 0.039 | 0.923 | 83 |
| *A. octavius* | 0.033 | 0.916 | 84 | 0.126 | 0.923 | 84 |
| *B. caecicola(OTU10)* | -0.016 | 0.916 | 85 | 0.188 | 0.923 | 85 |
| *D. longicatena* | 0.069 | 0.916 | 86 | 0.026 | 0.923 | 86 |
| *F. prausnitzii(OTU8)* | 0.099 | 0.916 | 87 | 0.175 | 0.923 | 87 |
| *O. valericigenes* | 0.009 | 0.916 | 88 | -0.001 | 0.923 | 88 |
| *R. hominis* | 0.021 | 0.916 | 89 | 0.002 | 0.923 | 89 |
| *P. secunda* | 0.003 | 0.916 | 90 | -0.240 | 0.923 | 90 |
| *B. caecicola(OTU4)* | -0.012 | 0.916 | 91 | -0.010 | 0.923 | 91 |
| Ruminococcaceae(OTU3) | 0.045 | 0.916 | 92 | 0.204 | 0.923 | 92 |

| | | | | | | |
|---|---|---|---|---|---|---|
| *A. ruminis(OTU2)* | 0.184 | 0.916 | 93 | 0.138 | 0.923 | 93 |
| *R. lactaris(OTU0)* | 0.000 | 0.916 | 94 | 0.000 | 0.923 | 94 |
| *B. caecicola(OTU72)* | 0.025 | 0.832 | 97 | 0.056 | 0.921 | 95 |
| *B. gallinaceum(OTU83)* | -0.059 | 0.963 | 39 | 0.140 | 0.917 | 96 |
| *B. propionicifaciens(OTU29)* | 0.005 | 0.916 | 73 | 0.181 | 0.912 | 97 |

Taxonomic annotation of OTUs, relative abundance, differential abundance, HITS score and intervention scores were provided and sorted by optimal cumulative intervention order. Differential abundance analysis was performed with Wilcoxon rank-sum test and adjusted with Benjamini-Hochberg method. HITS score was calculated using package Networkx of Python (3.6.0) with permutation test for its significance. Intervention score indicated the performance of the intervention by each candidate species in dynamic intervention modeling. And the optimal combinations of the keystone species with the maximum cumulative intervention score was determined by Iterative Feature Elimination.

**Table S3. The intervention effects of the keystone species, *P. loveana, A. indistinctus* and *D. pneumosinte*.**

| Abb. | Family | SI | DA | *P. loveana* | *A. indistinctus* | *D. pneumosintes* | Three keystone |
|---|---|---|---|---|---|---|---|
| *A. ruminis(OTU75)* | Lachnospiraceae | 2.541E-01 | -1 | 1 | 0 | 1 | 1 |
| *A. ruminis(OTU2)* | Lachnospiraceae | 1.377E-01 | -1 | 1 | 0 | 1 | 1 |
| *B. hansenii* | Lachnospiraceae | 1.155E-01 | -1 | 1 | 0 | 0 | 1 |
| Lachnospiraceae(OTU85) | Lachnospiraceae | -3.863E-02 | 1 | 1 | 0 | 1 | 1 |
| Lachnospiraceae(OTU55) | Lachnospiraceae | 7.266E-02 | -1 | 0 | 1 | 1 | 1 |
| *A. caccae* | Lachnospiraceae | 4.225E-02 | -1 | 1 | 0 | 0 | 1 |
| *E. rectale(OTU52)* | Lachnospiraceae | 1.388E-01 | -1 | 0 | 0 | 0 | 0 |
| *E. rectale(OTU47)* | Lachnospiraceae | 6.120E-02 | -1 | 1 | 0 | 0 | 1 |
| *B. producta(OTU81)* | Lachnospiraceae | 1.571E-01 | -1 | 1 | 0 | 1 | 1 |
| *B. producta(OTU32)* | Lachnospiraceae | 1.962E-01 | -1 | 1 | 0 | 0 | 1 |
| *F. saccharivorans* | Lachnospiraceae | 2.382E-01 | -1 | 1 | 0 | 0 | 1 |
| *C. amygdalinum* | Lachnospiraceae | 1.216E-02 | 1 | 0 | 0 | 0 | 0 |
| *B. hydrogenotrophica* | Lachnospiraceae | 9.967E-02 | -1 | 1 | 0 | 1 | 1 |
| *L. pacaense* | Lachnospiraceae | 1.110E-02 | 1 | 0 | 0 | 0 | 0 |
| *R. lactaris(OTU38)* | Lachnospiraceae | 8.973E-02 | 1 | 0 | 0 | 0 | 0 |
| *B. xylanolyticus* | Lachnospiraceae | 2.730E-03 | -1 | 0 | 0 | 0 | 0 |
| *E. oxidoreducens* | Lachnospiraceae | 1.542E-01 | -1 | 1 | 0 | 0 | 1 |
| *C. catus* | Lachnospiraceae | -2.579E-03 | 1 | 0 | 0 | 0 | 0 |
| *E. ramulus* | Lachnospiraceae | 1.817E-01 | -1 | 1 | 0 | 1 | 1 |
| *D. longicatena* | Lachnospiraceae | 2.579E-02 | -1 | 0 | 0 | 0 | 0 |
| *R. hominis* | Lachnospiraceae | 1.522E-03 | -1 | 0 | 0 | 0 | 0 |
| *C. comes* | Lachnospiraceae | 1.006E-01 | -1 | 0 | 0 | 1 | 1 |
| *M. faecis* | Lachnospiraceae | 8.501E-03 | 1 | 0 | 0 | 0 | 0 |

| | | | | | | | |
|---|---|---|---|---|---|---|---|
| *R. torques* | Lachnospiraceae | 3.877E-02 | -1 | 1 | 0 | 0 | 1 |
| *R. gnavus* | Lachnospiraceae | 3.367E-02 | -1 | 1 | 0 | 0 | 1 |
| *D. formicigenerans* | Lachnospiraceae | 2.310E-01 | -1 | 1 | 0 | 1 | 1 |
| *R. lactaris(OTU0)* | Lachnospiraceae | 0.000E+00 | -1 | 0 | 0 | 0 | 0 |
| *H. effluvii(OTU64)* | Lachnospiraceae | -1.254E-01 | 1 | 0 | 0 | 1 | 1 |
| Lachnospiraceae(OTU18) | Lachnospiraceae | 1.149E-01 | -1 | 0 | 0 | 0 | 0 |
| *H. effluvii(OTU77)* | Lachnospiraceae | -3.483E-02 | 1 | 0 | 0 | 1 | 1 |
| *F. prausnitzii(OTU80)* | Ruminococcaceae | 2.004E-01 | -1 | 0 | 0 | 1 | 1 |
| Ruminococcaceae(OTU3) | Ruminococcaceae | 2.041E-01 | -1 | 1 | 0 | 0 | 1 |
| *S. variabile(OTU44)* | Ruminococcaceae | 1.765E-02 | -1 | 0 | 0 | 1 | 1 |
| *Ruminiclostridium(OTU36)* | Ruminococcaceae | 4.583E-03 | 1 | 0 | 0 | 0 | 0 |
| *B. sp. K4410.MGS-46* | Ruminococcaceae | 3.999E-02 | -1 | 0 | 0 | 1 | 1 |
| *S. variabile(OTU93)* | Ruminococcaceae | 3.760E-03 | -1 | 0 | 0 | 1 | 1 |
| Ruminococcaceae(OTU23) | Ruminococcaceae | 5.041E-03 | -1 | 0 | 0 | 0 | 0 |
| Ruminococcaceae(OTU91) | Ruminococcaceae | -2.079E-02 | -1 | 0 | 0 | 0 | 0 |
| *F. prausnitzii(OTU8)* | Ruminococcaceae | 1.747E-01 | -1 | 0 | 0 | 1 | 1 |
| *Ruminiclostridium(OTU96)* | Ruminococcaceae | 4.982E-02 | -1 | 0 | 1 | 0 | 1 |
| *F. plautii* | Ruminococcaceae | 4.977E-02 | 1 | -1 | -1 | 0 | -1 |
| Ruminococcaceae(OTU46) | Ruminococcaceae | 3.426E-03 | -1 | 0 | 0 | 0 | 0 |

| | | | | | | | |
|---|---|---|---|---|---|---|---|
| *F. prausnitzii(OTU79)* | Ruminococcaceae | 0.000E+00 | -1 | 0 | 0 | 0 | 0 |
| *O. valericigenes* | Ruminococcaceae | -7.404E-04 | 1 | 0 | 0 | 0 | 0 |
| *F. prausnitzii(OTU68)* | Ruminococcaceae | 1.232E-02 | -1 | 0 | 0 | 0 | 0 |
| *R. lactaris(OTU43)* | Ruminococcaceae | -9.993E-04 | -1 | 0 | 0 | 0 | 0 |
| Ruminococcaceae(OTU22) | Ruminococcaceae | 5.693E-03 | -1 | 0 | 0 | 0 | 0 |
| *UBA1819(OTU14)* | Ruminococcaceae | 1.915E-01 | 1 | -1 | 0 | 0 | -1 |
| *P. ivorii* | Family XI | 1.363E-01 | 1 | 0 | -1 | 0 | -1 |
| *F. magna* | Family XI | 1.291E-01 | 1 | 0 | 0 | 0 | 0 |
| *E. peruensis* | Family XI | 2.469E-01 | 1 | 0 | -1 | 0 | -1 |
| *A. octavius* | Family XI | 1.265E-01 | 1 | 0 | -1 | 0 | -1 |
| *P. olsenii* | Family XI | 8.354E-02 | 1 | 0 | -1 | 0 | -1 |
| *Ezakiella(OTU94)* | Family XI | 1.806E-01 | 1 | -1 | 0 | 0 | -1 |
| *P. lacrimalis* | Family XI | 2.894E-01 | 1 | 0 | -1 | -1 | -1 |
| *B. gallinaceum(OTU27)* | Bacteroidaceae | 1.043E-01 | -1 | 0 | 0 | 0 | 0 |
| *B. gallinaceum(OTU42)* | Bacteroidaceae | 7.565E-02 | -1 | 0 | 0 | 0 | 0 |
| *B. gallinaceum(OTU83)* | Bacteroidaceae | 1.396E-01 | -1 | 0 | 1 | 0 | 1 |
| *B. caecicola(OTU51)* | Bacteroidaceae | 7.521E-02 | -1 | 0 | 1 | 0 | 1 |
| *B. caecicola(OTU84)* | Bacteroidaceae | 1.146E-02 | -1 | 0 | 0 | 0 | 0 |
| *B. propionicifaciens(OTU29)* | Bacteroidaceae | 1.809E-01 | -1 | 0 | 1 | 0 | 1 |
| *B. coprosuis(OTU62)* | Bacteroidaceae | 7.924E-02 | -1 | 0 | 0 | 0 | 0 |

| | | | | | | | |
|---|---|---|---|---|---|---|---|
| *B. propionicifaciens(OTU73)* | Bacteroidaceae | 1.712E-01 | -1 | 0 | 0 | 0 | 0 |
| *B. caecicola(OTU72)* | Bacteroidaceae | 5.611E-02 | -1 | 0 | 1 | 0 | 1 |
| *B. caecicola(OTU4)* | Bacteroidaceae | -9.908E-03 | 1 | 0 | 0 | 0 | 0 |
| *B. zoogleoformans* | Bacteroidaceae | 7.631E-02 | -1 | 0 | 0 | 0 | 0 |
| *B. caecicola(OTU10)* | Bacteroidaceae | 1.878E-01 | -1 | 0 | 0 | 0 | 0 |
| *B. gallinaceum(OTU1)* | Bacteroidaceae | -1.513E-02 | 1 | 0 | 0 | 0 | 0 |
| *B. caecicola(OTU70)* | Bacteroidaceae | -1.955E-02 | 1 | 0 | 0 | 0 | 0 |
| *B. caecicola(OTU16)* | Bacteroidaceae | -1.474E-02 | 1 | 0 | 0 | 0 | 0 |
| *B. propionicifaciens(OTU41)* | Bacteroidaceae | 1.067E-01 | -1 | 0 | 1 | 0 | 1 |
| *B. barnesiae* | Bacteroidaceae | 1.105E-01 | -1 | 0 | 0 | 1 | 1 |
| *B. caecicola(OTU86)* | Bacteroidaceae | -1.305E-01 | 1 | 0 | 0 | 0 | 0 |
| *B. coprosuis(OTU67)* | Bacteroidaceae | 9.792E-03 | -1 | 0 | 0 | 0 | 0 |
| *B. caecicola(OTU78)* | Bacteroidaceae | 1.218E-01 | -1 | 0 | 0 | 0 | 0 |
| *B. coprophilus* | Bacteroidaceae | 6.620E-47 | 1 | 0 | 0 | 0 | 0 |
| *B. magnum* | Bifidobacteriaceae | 1.349E-01 | -1 | 0 | 0 | 0 | 0 |
| *P. secunda* | Burkholderiaceae | -2.403E-01 | 1 | 1 | 1 | 0 | 1 |
| *C. pinnipediorum* | Campylobacteraceae | 6.495E-03 | 1 | 0 | 0 | 0 | 0 |
| *Bilophila(OTU30)* | Desulfovibrionaceae | -2.332E-02 | 1 | 0 | 0 | 0 | 0 |
| *Escherichia-Shigella(OTU88)* | Enterobacteriaceae | 6.113E-02 | 1 | 0 | 0 | 0 | 0 |
| *S5-A14a(OTU76)* | Family XIII | 1.619E-01 | 1 | 0 | 0 | -1 | -1 |

| | | | | | | | |
|---|---|---|---|---|---|---|---|
| *L. fallax* | Leuconostocaceae | 1.970E-05 | 1 | 0 | 0 | 0 | 0 |
| *W. ghanensis* | Leuconostocaceae | -1.727E-03 | 1 | 0 | 0 | 0 | 0 |
| *O. denticanis* | Marinifilaceae | -1.380E-02 | 1 | 0 | 0 | 0 | 0 |
| *R. lituseburensis* | Peptostreptococcaceae | 9.468E-02 | -1 | 0 | 0 | 0 | 0 |
| *I. bartlettii* | Peptostreptococcaceae | -6.565E-03 | 1 | 0 | 0 | 0 | 0 |
| *P. loveana* | Porphyromonadaceae | 3.457E-01 | 1 | -1 | 0 | 0 | -1 |
| *P. sp. 2007b* | Porphyromonadaceae | 3.758E-01 | 1 | 0 | -1 | -1 | -1 |
| *P. buccalis* | Prevotellaceae | 1.740E-01 | 1 | 0 | 0 | 0 | 0 |
| *A. indistinctus(OTU20)* | Rikenellaceae | 2.805E-03 | -1 | 0 | 0 | 0 | 0 |
| *A. indistinctus(OTU57)* | Rikenellaceae | 2.065E-01 | -1 | 0 | 1 | 1 | 1 |
| *A. indistinctus(OTU26)* | Rikenellaceae | 2.061E-02 | -1 | 0 | 0 | 0 | 0 |
| *S. thermophilus TH1435* | Streptococcaceae | 3.521E-02 | -1 | 1 | 0 | 0 | 1 |
| *P. chartae(OTU54)* | Tannerellaceae | -3.310E-02 | 1 | 0 | 0 | 0 | 0 |
| *P. chartae(OTU69)* | Tannerellaceae | 1.030E-03 | -1 | 0 | 0 | 0 | 0 |
| *D. pneumosintes* | Veillonellaceae | 2.939E-01 | 1 | 0 | -1 | -1 | -1 |

The abundance changes of every OTU when intervention were listed, in which -1, 0 and 1 means the abundance was down-regulated, no change and up-regulated. OTUs were grouped by family.

SI: Single species intervention score

DA: Differentail abundance (1:up-regulated, -1:down-regulated)

*P. loveana*: Intervention of *P. loveana* (1:up-regulated, -1:down-regulated, 0:no change)

*A. indistinctus*: Intervention of *A. indistinctus* (1:up-regulated, -1:down-regulated, 0:no change)

*D. pneumosintes*: Intervention of *D. pneumosintes* (1:up-regulated, -1:down-regulated, 0:no change)

**Table S4. KEGG module enrichment analysis on keystone species.**

| Module ID | Pathway annotation | | |
|---|---|---|---|
| | Pathway Level1 | Pathway Level2 | Module name |
| M00004 | Carbohydrate metabolism | Central carbohydrate metabolism | Pentose phosphate pathway (Pentose phosphate cycle) |
| M00009 | Carbohydrate metabolism | Central carbohydrate metabolism | Citrate cycle (TCA cycle, Krebs cycle) |
| M00011 | Carbohydrate metabolism | Central carbohydrate metabolism | Citrate cycle, second carbon oxidation, 2-oxoglutarate => oxaloacetate |
| M00532 | Carbohydrate metabolism | Other carbohydrate metabolism | Photorespiration |
| M00632 | Carbohydrate metabolism | Other carbohydrate metabolism | Galactose degradation, Leloir pathway, galactose => alpha-D-glucose-1P |
| M00740 | Carbohydrate metabolism | Other carbohydrate metabolism | Methylaspartate cycle |
| M00167 | Energy metabolism | Carbon fixation | Reductive pentose phosphate cycle, glyceraldehyde-3P => ribulose-5P |
| M00169 | Energy metabolism | Carbon fixation | CAM (Crassulacean acid metabolism), light |
| M00173 | Energy metabolism | Carbon fixation | Reductive citrate cycle (Arnon-Buchanan cycle) |
| M00374 | Energy metabolism | Carbon fixation | Dicarboxylate-hydroxybutyrate cycle |
| M00376 | Energy metabolism | Carbon fixation | 3-Hydroxypropionate bi-cycle |
| M00620 | Energy metabolism | Carbon fixation | Incomplete reductive citrate cycle, acetyl-CoA => oxoglutarate |
| M00346 | Energy metabolism | Methane metabolism | Formaldehyde assimilation, serine pathway |
| M00144 | Energy metabolism | ATP synthesis | NADH:quinone oxidoreductase, prokaryotes |
| M00149 | Energy metabolism | ATP synthesis | Succinate dehydrogenase, prokaryotes |
| M00157 | Energy metabolism | ATP synthesis | F-type ATPase, prokaryotes and chloroplasts |
| M00159 | Energy metabolism | ATP synthesis | V/A-type ATPase, prokaryotes |
| M00018 | Amino acid metabolism | Serine and threonine metabolism | Threonine biosynthesis, aspartate => homoserine => threonine |
| M00020 | Amino acid metabolism | Serine and threonine metabolism | Serine biosynthesis, glycerate-3P => serine |
| M00017 | Amino acid metabolism | Cysteine and methionine metabolism | Methionine biosynthesis, apartate => homoserine => methionine |
| M00034 | Amino acid metabolism | Cysteine and methionine metabolism | Methionine salvage pathway |
| M00019 | Amino acid metabolism | Branched-chain amino acid metabolism | Valine/isoleucine biosynthesis, pyruvate => valine / 2-oxobutanoate => isoleucine |

| | | | |
|---|---|---|---|
| M00432 | Amino acid metabolism | Branched-chain amino acid metabolism | Leucine biosynthesis, 2-oxoisovalerate => 2-oxoisocaproate |
| M00535 | Amino acid metabolism | Branched-chain amino acid metabolism | Isoleucine biosynthesis, pyruvate => 2-oxobutanoate |
| M00570 | Amino acid metabolism | Branched-chain amino acid metabolism | Isoleucine biosynthesis, threonine => 2-oxobutanoate => isoleucine |
| M00015 | Amino acid metabolism | Arginine and proline metabolism | Proline biosynthesis, glutamate => proline |
| M00028 | Amino acid metabolism | Arginine and proline metabolism | Ornithine biosynthesis, glutamate => ornithine |
| M00845 | Amino acid metabolism | Arginine and proline metabolism | Arginine biosynthesis, glutamate => acetylcitrulline => arginine |
| M00026 | Amino acid metabolism | Histidine metabolism | Histidine biosynthesis, PRPP => histidine |
| M00045 | Amino acid metabolism | Histidine metabolism | Histidine degradation, histidine => N-formiminoglutamate => glutamate |
| M00115 | Metabolism of cofactors and vitamins | Cofactor and vitamin metabolism | NAD biosynthesis, aspartate => NAD |
| M00117 | Metabolism of cofactors and vitamins | Cofactor and vitamin metabolism | Ubiquinone biosynthesis, prokaryotes, chorismate => ubiquinone |
| M00119 | Metabolism of cofactors and vitamins | Cofactor and vitamin metabolism | Pantothenate biosynthesis, valine/L-aspartate => pantothenate |
| M00121 | Metabolism of cofactors and vitamins | Cofactor and vitamin metabolism | Heme biosynthesis, plants and bacteria, glutamate => heme |
| M00122 | Metabolism of cofactors and vitamins | Cofactor and vitamin metabolism | Cobalamin biosynthesis, cobinamide => cobalamin |
| M00123 | Metabolism of cofactors and vitamins | Cofactor and vitamin metabolism | Biotin biosynthesis, pimeloyl-ACP/CoA => biotin |
| M00124 | Metabolism of cofactors and vitamins | Cofactor and vitamin metabolism | Pyridoxal biosynthesis, erythrose-4P => pyridoxal-5P |
| M00125 | Metabolism of cofactors and vitamins | Cofactor and vitamin metabolism | Riboflavin biosynthesis, GTP => riboflavin/FMN/FAD |
| M00126 | Metabolism of cofactors and vitamins | Cofactor and vitamin metabolism | Tetrahydrofolate biosynthesis, GTP => THF |
| M00127 | Metabolism of cofactors and vitamins | Cofactor and vitamin metabolism | Thiamine biosynthesis, AIR => thiamine-P/thiamine-2P |
| M00140 | Metabolism of cofactors and vitamins | Cofactor and vitamin metabolism | C1-unit interconversion, prokaryotes |
| M00572 | Metabolism of cofactors and vitamins | Cofactor and vitamin metabolism | Pimeloyl-ACP biosynthesis, BioC-BioH pathway, malonyl-ACP => pimeloyl-ACP |
| M00573 | Metabolism of cofactors and vitamins | Cofactor and vitamin metabolism | Biotin biosynthesis, BioI pathway, long-chain-acyl-ACP => pimeloyl-ACP => biotin |
| M00577 | Metabolism of cofactors and vitamins | Cofactor and vitamin metabolism | Biotin biosynthesis, BioW pathway, pimelate => pimeloyl-CoA => biotin |
| M00846 | Metabolism of cofactors and vitamins | Cofactor and vitamin metabolism | Siroheme biosynthesis, glutamate => siroheme |

Cont.

| Module ID | KO in Module | *P. loveana* | | | | *A. indistinctus* | | | | *D. pneumosintes* | | | |
|---|---|---|---|---|---|---|---|---|---|---|---|---|---|
| | | KO | OddsRatio | p-valule | FDR | KO | OddsRatio | p-valule | FDR | KO | OddsRatio | p-valule | FDR |
| M00004 | 15 | 4 | 10.20 | 1.43E-03 | 1.08E-02 | 5 | 13.43 | 1.34E-04 | 1.36E-03 | 6 | 18.90 | 6.14E-06 | 1.10E-04 |
| M00009 | 40 | 12 | 12.14 | 6.15E-09 | 3.10E-07 | 10 | 9.00 | 1.13E-06 | 2.33E-05 | 4 | 3.14 | 4.74E-02 | 2.56E-01 |
| M00011 | 34 | 12 | 15.45 | 7.31E-10 | 7.20E-08 | 10 | 11.25 | 2.12E-07 | 6.42E-06 | 4 | 3.77 | 2.82E-02 | 1.62E-01 |
| M00532 | 15 | 6 | 18.75 | 6.42E-06 | 1.01E-04 | 6 | 17.93 | 8.21E-06 | 1.16E-04 | 2 | 4.34 | 9.19E-02 | 4.53E-01 |
| M00632 | 4 | 3 | 84.09 | 1.60E-04 | 1.54E-03 | 3 | 80.41 | 1.82E-04 | 1.68E-03 | 1 | 9.39 | 1.30E-01 | 5.84E-01 |
| M00740 | 12 | 4 | 14.03 | 5.62E-04 | 4.81E-03 | 1 | 2.43 | 3.57E-01 | 1.00E+00 | 0 | 0.00 | 1.00E+00 | 1.00E+00 |
| M00167 | 11 | 3 | 10.51 | 5.52E-03 | 3.51E-02 | 3 | 10.05 | 6.22E-03 | 3.77E-02 | 4 | 16.16 | 3.74E-04 | 4.30E-03 |
| M00169 | 2 | 2 | inf | 1.19E-03 | 9.41E-03 | 2 | inf | 1.30E-03 | 9.67E-03 | 0 | 0.00 | 1.00E+00 | 1.00E+00 |
| M00173 | 43 | 14 | 13.70 | 9.61E-11 | 1.89E-08 | 12 | 10.48 | 2.47E-08 | 1.22E-06 | 6 | 4.59 | 3.30E-03 | 2.72E-02 |
| M00374 | 24 | 8 | 14.09 | 8.86E-07 | 2.18E-05 | 5 | 7.07 | 1.45E-03 | 1.03E-02 | 2 | 2.56 | 1.98E-01 | 8.41E-01 |
| M00376 | 21 | 5 | 8.78 | 6.26E-04 | 5.24E-03 | 5 | 8.39 | 7.60E-04 | 6.11E-03 | 4 | 6.65 | 5.17E-03 | 3.84E-02 |
| M00620 | 17 | 8 | 25.06 | 3.63E-08 | 1.43E-06 | 6 | 14.67 | 1.91E-05 | 2.59E-04 | 2 | 3.76 | 1.14E-01 | 5.54E-01 |
| M00346 | 10 | 3 | 12.01 | 4.12E-03 | 2.70E-02 | 4 | 17.89 | 2.97E-04 | 2.60E-03 | 2 | 7.05 | 4.41E-02 | 2.41E-01 |
| M00144 | 17 | 0 | 0.00 | 1.00E+00 | 1.00E+00 | 11 | 49.61 | 1.29E-12 | 5.07E-10 | 0 | 0.00 | 1.00E+00 | 1.00E+00 |
| M00149 | 4 | 3 | 84.09 | 1.60E-04 | 1.54E-03 | 3 | 80.41 | 1.82E-04 | 1.68E-03 | 0 | 0.00 | 1.00E+00 | 1.00E+00 |
| M00157 | 8 | 0 | 0.00 | 1.00E+00 | 1.00E+00 | 0 | 0.00 | 1.00E+00 | 1.00E+00 | 8 | inf | 1.86E-12 | 7.31E-10 |
| M00159 | 9 | 6 | 56.27 | 1.29E-07 | 3.90E-06 | 6 | 53.80 | 1.66E-07 | 5.45E-06 | 0 | 0.00 | 1.00E+00 | 1.00E+00 |
| M00018 | 10 | 3 | 12.01 | 4.12E-03 | 2.70E-02 | 3 | 11.48 | 4.65E-03 | 2.91E-02 | 5 | 28.32 | 1.03E-05 | 1.76E-04 |
| M00020 | 3 | 3 | inf | 4.12E-05 | 4.92E-04 | 3 | inf | 4.68E-05 | 5.27E-04 | 2 | 56.43 | 3.45E-03 | 2.72E-02 |
| M00017 | 11 | 3 | 10.51 | 5.52E-03 | 3.51E-02 | 4 | 15.33 | 4.54E-04 | 3.80E-03 | 3 | 10.59 | 5.40E-03 | 3.94E-02 |
| M00034 | 19 | 1 | 1.55 | 4.88E-01 | 1.00E+00 | 1 | 1.48 | 5.03E-01 | 1.00E+00 | 6 | 13.08 | 2.96E-05 | 4.17E-04 |
| M00019 | 6 | 1 | 5.59 | 1.90E-01 | 7.35E-01 | 5 | 134.35 | 3.52E-07 | 9.25E-06 | 0 | 0.00 | 1.00E+00 | 1.00E+00 |

| | | | | | | | | | | | | | |
|---|---|---|---|---|---|---|---|---|---|---|---|---|---|
| M00432 | 5 | 0 | 0.00 | 1.00E+00 | 1.00E+00 | 4 | 107.35 | 8.18E-06 | 1.16E-04 | 0 | 0.00 | 1.00E+00 | 1.00E+00 |
| M00535 | 4 | 0 | 0.00 | 1.00E+00 | 1.00E+00 | 4 | inf | 1.68E-06 | 2.89E-05 | 0 | 0.00 | 1.00E+00 | 1.00E+00 |
| M00570 | 8 | 1 | 3.99 | 2.45E-01 | 8.95E-01 | 5 | 44.78 | 3.09E-06 | 4.88E-05 | 0 | 0.00 | 1.00E+00 | 1.00E+00 |
| M00015 | 4 | 0 | 0.00 | 1.00E+00 | 1.00E+00 | 1 | 8.91 | 1.37E-01 | 4.57E-01 | 3 | 84.75 | 1.57E-04 | 1.93E-03 |
| M00028 | 14 | 1 | 2.15 | 3.89E-01 | 1.00E+00 | 4 | 10.73 | 1.26E-03 | 9.56E-03 | 0 | 0.00 | 1.00E+00 | 1.00E+00 |
| M00845 | 7 | 1 | 4.66 | 2.18E-01 | 8.19E-01 | 4 | 35.78 | 5.40E-05 | 5.91E-04 | 2 | 11.28 | 2.20E-02 | 1.29E-01 |
| M00026 | 17 | 0 | 0.00 | 1.00E+00 | 1.00E+00 | 7 | 18.85 | 1.10E-06 | 2.33E-05 | 1 | 1.76 | 4.48E-01 | 1.00E+00 |
| M00045 | 8 | 4 | 28.06 | 8.88E-05 | 9.46E-04 | 4 | 26.83 | 1.05E-04 | 1.09E-03 | 0 | 0.00 | 1.00E+00 | 1.00E+00 |
| M00115 | 7 | 5 | 70.25 | 9.67E-07 | 2.24E-05 | 5 | 67.17 | 1.20E-06 | 2.36E-05 | 2 | 11.28 | 2.20E-02 | 1.29E-01 |
| M00117 | 10 | 1 | 3.11 | 2.97E-01 | 1.00E+00 | 4 | 17.89 | 2.97E-04 | 2.60E-03 | 0 | 0.00 | 1.00E+00 | 1.00E+00 |
| M00119 | 6 | 4 | 56.13 | 2.01E-05 | 2.48E-04 | 4 | 53.67 | 2.38E-05 | 2.76E-04 | 0 | 0.00 | 1.00E+00 | 1.00E+00 |
| M00121 | 15 | 6 | 18.75 | 6.42E-06 | 1.01E-04 | 3 | 6.70 | 1.54E-02 | 8.31E-02 | 7 | 24.84 | 2.76E-07 | 8.36E-06 |
| M00122 | 10 | 8 | 112.83 | 8.34E-11 | 1.89E-08 | 2 | 6.69 | 4.83E-02 | 2.15E-01 | 8 | 113.72 | 7.85E-11 | 1.03E-08 |
| M00123 | 6 | 4 | 56.13 | 2.01E-05 | 2.48E-04 | 1 | 5.35 | 1.98E-01 | 6.19E-01 | 3 | 28.25 | 7.45E-04 | 7.16E-03 |
| M00124 | 6 | 5 | 140.51 | 2.84E-07 | 8.00E-06 | 4 | 53.67 | 2.38E-05 | 2.76E-04 | 1 | 5.63 | 1.89E-01 | 8.09E-01 |
| M00125 | 19 | 5 | 10.03 | 3.79E-04 | 3.47E-03 | 4 | 7.15 | 4.23E-03 | 2.73E-02 | 5 | 10.11 | 3.66E-04 | 4.30E-03 |
| M00126 | 17 | 7 | 19.71 | 8.26E-07 | 2.17E-05 | 7 | 18.85 | 1.10E-06 | 2.33E-05 | 6 | 15.46 | 1.43E-05 | 2.35E-04 |
| M00127 | 8 | 3 | 16.82 | 2.02E-03 | 1.48E-02 | 3 | 16.08 | 2.29E-03 | 1.58E-02 | 5 | 47.20 | 2.41E-06 | 4.75E-05 |
| M00140 | 5 | 3 | 42.04 | 3.91E-04 | 3.50E-03 | 2 | 17.85 | 1.21E-02 | 6.62E-02 | 3 | 42.38 | 3.82E-04 | 4.30E-03 |
| M00572 | 10 | 5 | 28.10 | 1.06E-05 | 1.61E-04 | 3 | 11.48 | 4.65E-03 | 2.91E-02 | 3 | 12.10 | 4.03E-03 | 3.05E-02 |
| M00573 | 6 | 4 | 56.13 | 2.01E-05 | 2.48E-04 | 1 | 5.35 | 1.98E-01 | 6.19E-01 | 3 | 28.25 | 7.45E-04 | 7.16E-03 |
| M00577 | 6 | 4 | 56.13 | 2.01E-05 | 2.48E-04 | 1 | 5.35 | 1.98E-01 | 6.19E-01 | 3 | 28.25 | 7.45E-04 | 7.16E-03 |
| M00846 | 15 | 3 | 7.00 | 1.37E-02 | 7.95E-02 | 2 | 4.12 | 1.00E-01 | 3.87E-01 | 7 | 24.84 | 2.76E-07 | 8.36E-06 |

KO annotation of keystone species was predicted by PICRUST2. The enrichment analysis of KEGG module was conducted with one-sided Fisher's exact test and adjusted with Benjamini-Hochberg method. Modules with FDR≤0.01 were considered significantly enriched modules.

#### Table S5. Differential abundance, HITS and intervention scores of the species in validation cohort.

| | Taxonomy | | | | Relative Abundance (%) | | |
|---|---|---|---|---|---|---|---|
| **Abb.** | **OTU** | **Class** | **Family** | **Species** | **normal_abun** | **nonaf_abun** | **cirrhosis_abun** |
| *D. formicigenerans* | 2f3327540661a5b628a1 03352e2d9731fb7bd13c | Clostridiales | Lachnospiraceae | *Dorea formicigenerans* | 0.243 | 0.251 | 0.348 |
| *H. porcina* | 3e5949b5957586cb373a 4fc1ff516e0dc5811c44 | Clostridiales | Lachnospiraceae | *Hespellia porcina* | 0.035 | 0.035 | 0.061 |
| *AD3011(OTU78)* | 16beeaa3ec13a7c9bebed dc43dddbbb389868ea2 | Clostridiales | Family XIII | *AD3011(OTU78)* | 0.068 | 0.032 | 0.032 |
| *B. sp. Marseille-P3087* | 46eaed43232c0befcbeaa 88c2a78e346521b86c0 | Clostridiales | Lachnospiraceae | *Blautia sp. Marseille-P3087* | 0.075 | 0.107 | 0.153 |
| *R. hominis(OTU106)* | 5c3307470a80cc114615 671dde3c6958e3de89de | Clostridiales | Lachnospiraceae | *Roseburia hominis(OTU106)* | 0.476 | 0.043 | 0.028 |
| *R. lituseburensis* | f6479b19f128dca30479 850ac88ee680e77f5a7d | Clostridiales | Peptostreptococcaceae | *Romboutsia lituseburensis* | 0.494 | 1.023 | 0.054 |
| *E. ramulus(OTU103)* | dd04fe7034b9f803cd16f 1d8755679574b681e38 | Clostridiales | Lachnospiraceae | *Eubacterium ramulus(OTU103)* | 0.119 | 0.027 | 0.067 |
| *A. caccae* | 3ab368af55d977e3b60b 558ae78e83f77bb5ae30 | Clostridiales | Lachnospiraceae | *Anaerostipes caccae* | 1.529 | 0.817 | 2.212 |
| *F. saccharivorans* | 12821fc1962342d552c7 5c911fee8353f055070e | Clostridiales | Lachnospiraceae | *Fusicatenibacter saccharivorans* | 1.262 | 0.747 | 0.821 |
| *B. hansenii(OTU14)* | fe1664de87020723ce6e 1a3aff25cc3b23423694 | Clostridiales | Lachnospiraceae | *Blautia hansenii(OTU14)* | 0.847 | 1.035 | 1.389 |
| *C. eutactus* | 4b9beaa541cbded795edf 2a6094c961b9818cf45 | Clostridiales | Lachnospiraceae | *Coprococcus eutactus* | 0.230 | 0.471 | 0.195 |

| | | | | | | | |
|---|---|---|---|---|---|---|---|
| *B. sp. K4410.MGS-46* | 8065d0c2d70ace77e99b 3731455f32317b8d2185 | Clostridiales | Ruminococcaceae | *Butyricicoccus sp. K4410.MGS-46* | 0.056 | 0.036 | 0.061 |
| *C. hylemonae* | 4ff5c5d092eba42696f4d ca89b3061f35ce61530 | Clostridiales | Lachnospiraceae | *Clostridium hylemonae* | 0.199 | 0.039 | 0.056 |
| Erysipelotrichaceae(OTU 96) | f1ea8d661cc5a8d536ac9 1ebb62fb64b8b4e3dae | Erysipelotrichales | Erysipelotrichaceae | Erysipelotrichaceae(O TU96) | 1.256 | 1.039 | 0.250 |
| *F. prausnitzii(OTU92)* | 10b6c57e4f5116e43a58 0f595da27e809b5e5743 | Clostridiales | Ruminococcaceae | *Faecalibacterium prausnitzii(OTU92)* | 0.423 | 0.425 | 0.152 |
| *E. ramosum(OTU89)* | cdccef0b1896c438c380a 9533859660b900097be | Erysipelotrichales | Erysipelotrichaceae | *Erysipelatoclostridium ramosum(OTU89)* | 0.080 | 0.040 | 0.090 |
| *B. hydrogenotrophica(OTU 118)* | 242649cefb9a3d66d218 b2eb9d5f66594456b8d0 | Clostridiales | Lachnospiraceae | *Blautia hydrogenotrophica(OT U118)* | 0.343 | 0.027 | 0.189 |
| *L. phytofermentans* | 45a35f09ab07a4d097c3 940123745f0f1c7d3bbb | Clostridiales | Lachnospiraceae | *Lachnoclostridium phytofermentans* | 0.072 | 0.074 | 0.601 |
| *C. xylanolyticum* | 71822b27140f23c7990c 8b9d58b12b0354b97c9c | Clostridiales | Lachnospiraceae | *Clostridium xylanolyticum* | 0.010 | 0.010 | 0.005 |
| *E. ramosum(OTU77)* | 61da89b142e6949e8752 55103d0d9a60db6e1b10 | Erysipelotrichales | Erysipelotrichaceae | *Erysipelatoclostridium ramosum(OTU77)* | 0.440 | 0.181 | 0.249 |
| *A. glycaniphila(OTU81)* | e2bc23d00411acdc5f6ec 76dadfc25ec45c6f567 | Verrucomicrobiales | Akkermansiaceae | *Akkermansia glycaniphila(OTU81)* | 1.375 | 1.733 | 1.449 |
| *E. oxidoreducens* | b135b820362617faaa43 9d6711f945e7e1e08cd6 | Clostridiales | Lachnospiraceae | *Eubacterium oxidoreducens* | 0.597 | 0.596 | 0.312 |
| *B. producta* | 1d4a7ecd548fc7f32ecc9 2a1562431443c920bdc | Clostridiales | Lachnospiraceae | *Blautia producta* | 0.722 | 0.539 | 0.712 |

| | | | | | | | |
|---|---|---|---|---|---|---|---|
| *R. hungatei(OTU107)* | d18241496baec625c74b173f59a6a0d192ab81fc | Clostridiales | Ruminococcaceae | *Ruminiclostridium hungatei(OTU107)* | 0.030 | 0.039 | 0.013 |
| *R. gnavus(OTU97)* | 171f468b34a93a5410d19bd51172a6671375edbc | Clostridiales | Lachnospiraceae | *Ruminococcus gnavus(OTU97)* | 0.551 | 0.464 | 0.607 |
| *B. barnesiae(OTU102)* | e28d4ae021cf98ce300cf2677b764b26ab858a79 | Bacteroidales | Bacteroidaceae | *Bacteroides barnesiae(OTU102)* | 0.108 | 0.142 | 0.210 |
| *B. hydrogenotrophica(OTU124)* | 26b6322c365bfcdcc5c06bdda44ee87eebebd3de | Clostridiales | Lachnospiraceae | *Blautia hydrogenotrophica(OTU124)* | 0.049 | 0.002 | 0.012 |
| *A. naeslundii* | 21eee70d5297e76473a8b9e0f47008c788a73276 | Actinomycetales | Actinomycetaceae | *Actinomyces naeslundii* | 0.023 | 0.040 | 0.034 |
| *E. caecimuris* | 0b876fed0626b041a5ab6564b911a0cd940d455d | Coriobacteriales | Eggerthellaceae | *Enterorhabdus caecimuris* | 0.007 | 0.004 | 0.003 |
| *E. rectale(OTU12)* | 07df84235f76f8f66d918d96fcd3b9661d3a3a51 | Clostridiales | Lachnospiraceae | *Eubacterium rectale(OTU12)* | 0.129 | 0.005 | 0.054 |
| *L. multipara(OTU37)* | 7166331027f0d6dceeb54c68a5878844349188d9 | Clostridiales | Lachnospiraceae | *Lachnospira multipara(OTU37)* | 0.094 | 0.063 | 0.099 |
| *A. glycaniphila(OTU110)* | 88dff74ce990a14706a5e6e96d731e219236cc9b | Verrucomicrobiales | Akkermansiaceae | *Akkermansia glycaniphila(OTU110)* | 0.096 | 0.080 | 0.172 |
| *CAG-352(OTU123)* | 7db290f9682e85efafd163b40f4d50d92deb3194 | Clostridiales | Ruminococcaceae | *CAG-352(OTU123)* | 0.852 | 0.003 | 0.003 |
| Lachnospiraceae(OTU4) | a0076e37a7d89de1eb15d0a9ebeadabf693b76e1 | Clostridiales | Lachnospiraceae | Lachnospiraceae(OTU4) | 0.034 | 0.037 | 0.031 |
| *P. psychrophila* | dc442c6a133138407ec45653f7df631bc25d5248 | Pseudomonadales | Pseudomonadaceae | *Pseudomonas psychrophila* | 6.617 | 0.005 | 0.013 |

| | | | | | | | |
|---|---|---|---|---|---|---|---|
| Ruminococcaceae(OTU52) | 90e449e0901749c492d2c2ee93571381090b0d94 | Clostridiales | Ruminococcaceae | Ruminococcaceae(OTU52) | 0.192 | 0.022 | 0.082 |
| *F. prausnitzii(OTU61)* | eaa89e412e9ee480b7019f5b31997726bb8661b3 | Clostridiales | Ruminococcaceae | *Faecalibacterium prausnitzii(OTU61)* | 3.957 | 2.356 | 1.050 |
| *B. ihuae(OTU101)* | c1d03cfa67e45fe3ad18ae4005c3a24d81fa8778 | Bacteroidales | Bacteroidaceae | *Bacteroides ihuae(OTU101)* | 1.806 | 0.923 | 0.663 |
| *L. pacaense(OTU93)* | cbd1dd0959b6c88fa5a73e7da59abec48f77fe09 | Clostridiales | Lachnospiraceae | *Lachnoclostridium pacaense(OTU93)* | 0.056 | 0.017 | 0.140 |
| *R. sp. N15.MGS-57* | cce8d260ef05ec8f52d1fe0d314543f5094283d6 | Selenomonadales | Acidaminococcaceae | *Ruminococcus sp. N15.MGS-57* | 0.880 | 0.110 | 0.274 |
| *A. indistinctus(OTU108)* | b0b3aa7030d3cd21aa8ced7b41d668b9df07635e | Bacteroidales | Rikenellaceae | *Alistipes indistinctus(OTU108)* | 0.215 | 0.025 | 0.120 |
| *S. lutetiensis* | 34b29086276f6215476afcaa8bd8c4daa9f79a8c | Lactobacillales | Streptococcaceae | *Streptococcus lutetiensis* | 0.060 | 0.201 | 0.640 |
| *T. sanguinis* | 339464b70eaaf1c574e5eb9ca829d3f8c0df833b | Erysipelotrichales | Erysipelotrichaceae | *Turicibacter sanguinis* | 0.051 | 0.083 | 0.048 |
| Ruminococcaceae(OTU25) | 9e5551b92019ab14f7339e0924c8d88f5ead5cc0 | Clostridiales | Ruminococcaceae | Ruminococcaceae(OTU25) | 0.171 | 0.025 | 0.043 |
| *Ruminiclostridium(OTU90)* | b6c85c9e12449d66ac5b62da812df9dde60d1d2c | Clostridiales | Ruminococcaceae | *Ruminiclostridium(OTU90)* | 0.089 | 0.046 | 0.119 |
| *UBA1819(OTU105)* | b8f9520f69922ac0d9e6deacbca4d610257d8563 | Clostridiales | Ruminococcaceae | *UBA1819(OTU105)* | 0.105 | 0.056 | 0.050 |
| *K. alysoides* | 945439d1f216d1088f6f2da5c74c3f43ffcf7a5d | Clostridiales | Lachnospiraceae | *Kineothrix alysoides* | 0.073 | 0.142 | 0.133 |

| | | | | | | | |
|---|---|---|---|---|---|---|---|
| *B. hydrogenotrophica(OTU7)* | 1a7ef1856ae5858ab962562c01c00eeead6c6648 | Clostridiales | Lachnospiraceae | *Blautia hydrogenotrophica(OTU7)* | 1.098 | 0.509 | 0.715 |
| *B. hansenii(OTU11)* | a7b60ca68019d05595466f955775ac5726c61240 | Clostridiales | Lachnospiraceae | *Blautia hansenii(OTU11)* | 1.321 | 1.637 | 4.359 |
| Ruminococcaceae(OTU115) | c11bb277ef45c226027ee3c7b25bb396fab7b2ef | Clostridiales | Ruminococcaceae | Ruminococcaceae(OTU115) | 0.068 | 0.010 | 0.017 |
| *R. lactatiformans* | b426157d1512c2c137df2465628640dcfbb90cb1 | Clostridiales | Ruminococcaceae | *Ruthenibacterium lactatiformans* | 0.021 | 0.019 | 0.038 |
| *P. secunda* | 1971c99c7ccabcf8eecd59cd442bc8234e1d1680 | Betaproteobacteriales | Burkholderiaceae | *Parasutterella secunda* | 0.159 | 0.045 | 0.056 |
| *C. cystitidis* | 56b9a169b58b500031b0fa973de40279e6aa304b | Corynebacteriales | Corynebacteriaceae | *Corynebacterium cystitidis* | 0.003 | 0.002 | 0.002 |
| *S. pseudolugdunensis* | ea4a29b8b59fb031b08cf1a7575658013dd45baa | Bacillales | Staphylococcaceae | *Staphylococcus pseudolugdunensis* | 0.038 | 0.033 | 0.472 |
| *Bilophila(OTU79)* | a8a500cff13b924b2879b627f6a153fe60d4041d | Desulfovibrionales | Desulfovibrionaceae | *Bilophila(OTU79)* | 0.068 | 0.110 | 0.046 |
| *R. inulinivorans* | 4743356252242920c9f26cf1217f05a86ef3248f | Clostridiales | Lachnospiraceae | *Roseburia inulinivorans* | 0.097 | 0.016 | 0.014 |
| Ruminococcaceae(OTU44) | 8af975f74bdf06d47aaa762096cb2d181f5af001 | Clostridiales | Ruminococcaceae | Ruminococcaceae(OTU44) | 0.158 | 0.011 | 0.047 |
| *R. bromii* | 0fab7ba69e9a53216d231dd0458b41129e4515bd | Clostridiales | Ruminococcaceae | *Ruminococcus bromii* | 0.835 | 0.584 | 1.298 |
| *R. hungatei(OTU63)* | 49ab52c9da5c10c5c05969f6a72a91a1c6c8bc65 | Clostridiales | Ruminococcaceae | *Ruminiclostridium hungatei(OTU63)* | 0.352 | 0.002 | 0.017 |

| | | | | | | | |
|---|---|---|---|---|---|---|---|
| *B. cucumis* | 0aadd7693dfc5e2e04e57b530101f36127951b00 | Bacillales | None | *Bacillus cucumis* | 0.044 | 0.972 | 0.005 |
| *D. pneumosintes* | c3626e26a31e3108bf91f06bb550697ddfd540a4 | Selenomonadales | Veillonellaceae | *Dialister pneumosintes* | 0.130 | 0.240 | 0.535 |
| *Ruminiclostridium(OTU62)* | 5fefe85ee87c6543858517caa7bb445c3cba1c00 | Clostridiales | Ruminococcaceae | *Ruminiclostridium(OTU62)* | 0.044 | 0.018 | 0.029 |
| *R. hominis(OTU29)* | 1831466480c94b307f8527f88e4b6ee339db0c10 | Clostridiales | Lachnospiraceae | *Roseburia hominis(OTU29)* | 0.119 | 0.071 | 0.385 |
| *B. magnum* | 40534050235d816268f98a1aca56cde478e66a00 | Bifidobacteriales | Bifidobacteriaceae | *Bifidobacterium magnum* | 0.839 | 1.980 | 7.167 |
| *I. bartlettii* | c2a7b89a736f924b4c68f4043efa8e4d05c45708 | Clostridiales | Peptostreptococcaceae | *Intestinibacter bartlettii* | 0.106 | 0.171 | 0.107 |
| *P. chartae(OTU125)* | fe5cc67bc49e188dd6592eef4f718cfdb21d93c3 | Bacteroidales | Tannerellaceae | *Parabacteroides chartae(OTU125)* | 0.430 | 0.094 | 0.248 |
| *H. effluvii* | a1898f9fbef02022cacb08de810f11491f49b957 | Clostridiales | Lachnospiraceae | *Hungatella effluvii* | 0.393 | 0.019 | 0.209 |
| *C. faecale* | e1e772d8a28ecc6e94463e9789e9dd4776d15556 | Corynebacteriales | Corynebacteriaceae | *Corynebacterium faecale* | 0.004 | 0.003 | 0.006 |
| *A. senegalensis* | 6abf6ac7c19c19bff2dcef35deb27d681d375c2b | Clostridiales | Family XI | *Anaerococcus senegalensis* | 0.003 | 0.002 | 0.003 |
| *P. chartae(OTU70)* | d3379224de8a6ad25d0f135fe666cfd0a8ce554e | Bacteroidales | Tannerellaceae | *Parabacteroides chartae(OTU70)* | 0.480 | 0.058 | 0.154 |
| *Ezakiella(OTU53)* | dc0bc1b9271f066ac72e2a607f221a12619e253d | Clostridiales | Family XI | *Ezakiella(OTU53)* | 0.007 | 0.002 | 0.013 |

| | | | | | | | |
|---|---|---|---|---|---|---|---|
| Ruminococcaceae(OTU74) | 60661167d1f8deb1c315 42a83c3a59f364508501 | Clostridiales | Ruminococcaceae | Ruminococcaceae(OTU74) | 0.144 | 0.014 | 0.099 |
| *T. nexilis* | d323e75e73433f6c7243 a543208f95d8aaa43288 | Clostridiales | Lachnospiraceae | *Tyzzerella nexilis* | 0.467 | 0.338 | 0.065 |
| *L. pacaense(OTU122)* | 81ff288bda3c6aeb68ae2 4b265180372959ca74d | Clostridiales | Lachnospiraceae | *Lachnoclostridium pacaense(OTU122)* | 0.032 | 0.015 | 0.027 |
| *B. sp. YHC-4* | 524d5509ecc18d29af0b be333938568425e56dd5 | Clostridiales | Lachnospiraceae | *Blautia sp. YHC-4* | 0.387 | 0.223 | 0.280 |
| *B. xylanolyticus* | fba450290326ad4d04b6 d7ba67fd557113c19c57 | Clostridiales | Lachnospiraceae | *Bacteroides xylanolyticus* | 0.523 | 0.092 | 0.064 |
| *S. variabile(OTU104)* | 3c612a1ee0b3bd27cdf6 2a3d5d2b20d4daed3b01 | Clostridiales | Ruminococcaceae | *Subdoligranulum variabile(OTU104)* | 1.041 | 0.406 | 1.050 |
| *A. ruminis* | e07591b0049b8831e12c bb7caca207d48628bf53 | Clostridiales | Lachnospiraceae | *Agathobacter ruminis* | 2.800 | 2.596 | 0.869 |
| *C. innocuum* | 43dfb5c51be438f5eedd6 b2e655a818a92490ddc | Erysipelotrichales | Erysipelotrichaceae | *Clostridium innocuum* | 0.050 | 0.011 | 0.070 |
| *V. criceti* | 9698ee80039ca2196853 dab6ce9214a29a3562c6 | Selenomonadales | Veillonellaceae | *Veillonella criceti* | 0.010 | 0.010 | 3.218 |
| Lachnospiraceae(OTU86) | c123a770771185e7e43f d102c1e6347e9eb3a2f7 | Clostridiales | Lachnospiraceae | Lachnospiraceae(OTU86) | 0.421 | 0.020 | 0.031 |
| *S. thermophilus TH1435* | b4fd15d98cc6502ba4bb 5390d3fd8e8ce0385286 | Lactobacillales | Streptococcaceae | *Streptococcus thermophilus TH1435* | 0.283 | 1.497 | 0.971 |
| *R. lactaris(OTU83)* | 97b18743ced53a3753fb 858108fefb8a9ba036bd | Clostridiales | Lachnospiraceae | *Ruminococcus lactaris(OTU83)* | 0.318 | 0.167 | 0.382 |

| | | | | | | | |
|---|---|---|---|---|---|---|---|
| *E. ramulus(OTU80)* | 5eabf30cb6e4f55f6a6f7d2db306474db18bb669 | Clostridiales | Ruminococcaceae | *Eubacterium ramulus(OTU80)* | 0.203 | 0.011 | 0.128 |
| Ruminococcaceae(OTU75) | 176e5c52f5cecaaaec1ff8c19c4c6a7bcd390aba | Clostridiales | Ruminococcaceae | Ruminococcaceae(OTU75) | 0.518 | 0.044 | 0.167 |
| *O. hirsuta* | 425d8d40aa61788d154c087e91d5057aa20c4442 | Chloroplast | None | *Okeania hirsuta* | 0.007 | 0.006 | 0.006 |
| Lachnospiraceae(OTU68) | f0da08923004a661f9c5980dbfe67306f05e301a | Clostridiales | Lachnospiraceae | Lachnospiraceae(OTU68) | 0.073 | 0.345 | 0.053 |
| *A. equolifaciens* | c494a72102793289e9a70a5b2f41cc10d347b8cd | Coriobacteriales | Eggerthellaceae | *Adlercreutzia equolifaciens* | 0.035 | 0.014 | 0.031 |
| *B. coprophilus* | 5cf26424a75b88a07cebd8237e26d4140386495c | Bacteroidales | Bacteroidaceae | *Bacteroides coprophilus* | 2.713 | 0.975 | 4.265 |
| Christensenellaceae(OTU59) | d7fffca6ce3c49792d705378ce321222b2181ef0 | Clostridiales | Christensenellaceae | Christensenellaceae(OTU59) | 0.182 | 0.125 | 0.144 |
| *L. pacaense(OTU54)* | 3e03d0dc7b060a2ce0dd0ffdca3fef1fc022b80d | Clostridiales | Lachnospiraceae | *Lachnoclostridium pacaense(OTU54)* | 0.061 | 0.021 | 0.076 |
| *F. plautii* | d49eacfeb41d662ad2cfa153d67bd71301266776 | Clostridiales | Ruminococcaceae | *Flavonifractor plautii* | 0.065 | 0.089 | 0.285 |
| *E. rectale(OTU51)* | af50580985a38ec3bcbc6b2076c9c9c21a33a19a | Clostridiales | Lachnospiraceae | *Eubacterium rectale(OTU51)* | 0.131 | 0.099 | 0.161 |
| *A. sp. AL-1* | a274b1d5902c14f41aa9bb23efdc4c7515af231f | Bacteroidales | Rikenellaceae | *Alistipes sp. AL-1* | 0.234 | 0.015 | 0.019 |
| *R. gnavus(OTU47)* | 48756705930044845fe1761fa6da3630522a4ef0 | Clostridiales | Lachnospiraceae | *Ruminococcus gnavus(OTU47)* | 0.329 | 0.778 | 0.531 |

| | | | | | | | |
|---|---|---|---|---|---|---|---|
| *C. comes* | ddae9a7fd743a629690a 92566615f797d36764f1 | Clostridiales | Lachnospiraceae | *Coprococcus comes* | 0.051 | 0.034 | 0.044 |
| *R. albus* | 93acb379a33cb2c4c074 284c70bdf163e7ff76ad | Clostridiales | Ruminococcaceae | *Ruminococcus albus* | 0.195 | 0.212 | 0.089 |
| *B. barnesiae(OTU40)* | e9da02b57b4e5fcd6ec3c 6a7308a44b1e1cd3a80 | Bacteroidales | Bacteroidaceae | *Bacteroides barnesiae(OTU40)* | 0.397 | 0.155 | 0.222 |
| *B. paurosaccharolyticus* | d849afc1e3de091c2646e 176f0f009c49e528ac0 | Bacteroidales | Bacteroidaceae | *Bacteroides paurosaccharolyticus* | 0.399 | 0.071 | 0.143 |
| *B. coprosuis* | cd4d5c1364663f21332a 8dd6ce46b226eea40551 | Bacteroidales | Bacteroidaceae | *Bacteroides coprosuis* | 0.133 | 0.100 | 0.141 |
| *S. variabile(OTU33)* | 4e4339ab92c072ae49a5 95333c4961607b1eb366 | Clostridiales | Ruminococcaceae | *Subdoligranulum variabile(OTU33)* | 1.182 | 0.757 | 2.351 |
| *E. hermanniensis* | cdabc39e03c431c9371d ce614d05d189379ad617 | Lactobacillales | Enterococcaceae | *Enterococcus hermanniensis* | 4.037 | 1.603 | 2.430 |
| *C. tanakaei* | 6da9624f67e5f1560204 7ba96a034d3dd9765e7b | Coriobacteriales | Coriobacteriaceae | *Collinsella tanakaei* | 0.268 | 1.012 | 0.786 |
| *E. rectale(OTU30)* | 436ed55c566ef292159e 45206fef12ca75fdca22 | Clostridiales | Lachnospiraceae | *Eubacterium rectale(OTU30)* | 0.717 | 1.005 | 1.304 |
| *A. indistinctus(OTU26)* | 025c3c0b72c2a0391e82 ee8845a0b060ea61bbb0 | Bacteroidales | Rikenellaceae | *Alistipes indistinctus(OTU26)* | 0.198 | 0.012 | 0.024 |
| Lachnospiraceae(OUT23) | 9d25e88c6107b366b445 e7de89448129222f964d | Clostridiales | Lachnospiraceae | Lachnospiraceae(OUT 23) | 0.012 | 0.007 | 0.016 |
| *O. jeddahense* | c3162b02a2d3d97d9772 108c246fd56c81ef1275 | Bacillales | Bacillaceae | *Oceanobacillus jeddahense* | 0.775 | 5.871 | 0.359 |

| | | | | | | | |
|---|---|---|---|---|---|---|---|
| *R. gauvreauii* | 64cc797b32cf29750a54 539aa3251851ac9f7894 | Clostridiales | Lachnospiraceae | *Ruminococcus gauvreauii* | 0.023 | 0.013 | 0.028 |
| *C. europaeus* | 0a40c389b2c0ae23f865e 444feebf820a9d353e6 | Enterobacteriales | Enterobacteriaceae | *Citrobacter europaeus* | 3.162 | 0.029 | 2.522 |
| *B. ihuae(OTU16)* | 7b38ac67bdd77bf68971 c5d1c9a637eed6b42f65 | Bacteroidales | Bacteroidaceae | *Bacteroides ihuae(OTU16)* | 5.837 | 5.300 | 3.445 |
| *C. autoethanogenum* | f152e68345172c4c0bd4 3c66dedf60acdd99fb72 | Clostridiales | Clostridiaceae 1 | *Clostridium autoethanogenum* | 0.115 | 0.484 | 0.160 |
| *E. tayi* | e90b611ddf657ec21be5 cb12621d1122a11eb192 | Clostridiales | Lachnospiraceae | *Eisenbergiella tayi* | 0.025 | 0.031 | 0.069 |
| *L. multipara(OTU6)* | 6c967ed9b68e96299f65 615180cf94efaf5f93ec | Clostridiales | Lachnospiraceae | *Lachnospira multipara(OTU6)* | 0.135 | 0.159 | 0.041 |
| *D. longicatena* | f319b2de319bd91c97ac 339c71a881594b34a156 | Clostridiales | Lachnospiraceae | *Dorea longicatena* | 0.560 | 1.596 | 0.631 |
| *E. sinensis* | 27ce7f321d9d8556bbc8f 16430c9795b2690a3ea | Coriobacteriales | Eggerthellaceae | *Eggerthella sinensis* | 0.119 | 0.038 | 0.061 |
| *Ruminiclostridium(OTU2 )* | 77f20b8915b46cef9458c 74f13e4281822a835d7 | Clostridiales | Ruminococcaceae | *Ruminiclostridium(OT U2)* | 0.094 | 0.049 | 0.069 |
| *B. graminisolvens* | bbe2a34c279534c0d745 d5189b6e064db7ffe9b8 | Bacteroidales | Bacteroidaceae | *Bacteroides graminisolvens* | 0.613 | 0.085 | 0.582 |
| *P. nigrescens* | d2023745a4597a48a9eb d12eda7c373b10a0d677 | Bacteroidales | Prevotellaceae | *Prevotella nigrescens* | 0.620 | 3.732 | 0.168 |
| *T. glycolicus* | bdd613213365a4931bf4 b0d14833329c48b0670e | Clostridiales | Peptostreptococcaceae | *Terrisporobacter glycolicus* | 0.027 | 0.028 | 0.005 |

| | | | | | | | |
|---|---|---|---|---|---|---|---|
| *O. valericigenes* | 71779fab49fba8ab4f2d6 8b74c5a9b70dd48f475 | Clostridiales | Ruminococcaceae | *Oscillibacter valericigenes* | 0.086 | 0.062 | 0.012 |
| *B. barnesiae(OTU73)* | a2368c772c7357241588 05a67408e4067b4f507b | Bacteroidales | Bacteroidaceae | *Bacteroides barnesiae(OTU73)* | 0.868 | 0.440 | 0.229 |
| Lachnospiraceae(OTU42 ) | 97507b6f378610511f55 7e17f42b076714044221 | Clostridiales | Lachnospiraceae | Lachnospiraceae(OTU 42) | 0.706 | 0.215 | 0.043 |
| *P. soli* | b8894367786589691928 ee9d7e771168fc2c6399 | Bacillales | Planococcaceae | *Psychrobacillus soli* | 2.827 | 7.338 | 0.005 |
| *R. lactaris(OTU117)* | 6f6159f166ea40e5f1001 37ca85a9d142fdb91cc | Clostridiales | Lachnospiraceae | *Ruminococcus lactaris(OTU117)* | 0.052 | 0.485 | 0.930 |
| *Escherichia-Shigella(OTU72)* | 7e3f625115c48cf9721e9 ffe63b3c48b5149ba18 | Enterobacteriales | Enterobacteriaceae | *Escherichia-Shigella(OTU72)* | 8.034 | 12.447 | 20.860 |
| *R. torques* | ba9d37124891e5eac8d8 02723297e54ec38ab03d | Clostridiales | Lachnospiraceae | *Ruminococcus torques* | 0.138 | 0.159 | 0.007 |
| *L. hircilactis* | 84cd82846ea355b3052d ff451eb7c5cd7b7abb16 | Lactobacillales | Streptococcaceae | *Lactococcus hircilactis* | 0.449 | 2.681 | 0.012 |

Cont.

| Abb. | Differential abundance | | | | | |
|---|---|---|---|---|---|---|
| | nonaf_vs_normal_abun_pval | nonaf_vs_normal_abun_fdr | cirrhosis_vs_normal_abun_pval | cirrhosis_vs_normal_abun_fdr | cirrhosis_vs_nonaf_pval | cirrhosis_vs_nonaf_abun_fdr |
| *D. formicigenerans* | 3.91E-01 | 6.94E-01 | 1.05E-01 | 2.60E-01 | 4.14E-01 | 8.27E-01 |
| *H. porcina* | 1.00E+00 | 1.00E+00 | 7.86E-01 | 8.53E-01 | 6.18E-01 | 8.40E-01 |
| *AD3011(OTU78)* | 5.48E-01 | 7.48E-01 | 2.30E-02 | 1.16E-01 | 4.14E-01 | 8.27E-01 |
| *B. sp. Marseille-P3087* | 4.05E-01 | 6.94E-01 | 4.50E-01 | 5.66E-01 | 9.28E-01 | 9.58E-01 |
| *R. hominis(OTU106)* | 2.11E-01 | 5.71E-01 | 5.11E-03 | 6.83E-02 | 1.38E-01 | 6.26E-01 |
| *R. lituseburensis* | 2.11E-01 | 5.71E-01 | 9.35E-04 | 4.25E-02 | 8.72E-04 | 5.54E-02 |
| *E. ramulus(OTU103)* | 3.52E-01 | 6.57E-01 | 2.77E-01 | 4.05E-01 | 5.86E-01 | 8.40E-01 |
| *A. caccae* | 6.77E-01 | 8.19E-01 | 6.11E-01 | 7.12E-01 | 7.85E-01 | 8.99E-01 |
| *F. saccharivorans* | 2.11E-01 | 5.71E-01 | 6.31E-02 | 2.03E-01 | 3.18E-01 | 7.77E-01 |
| *B. hansenii(OTU14)* | 6.41E-01 | 8.19E-01 | 7.07E-02 | 2.09E-01 | 9.60E-02 | 5.81E-01 |
| *C. eutactus* | 6.77E-01 | 8.19E-01 | 1.56E-01 | 2.84E-01 | 1.16E-01 | 5.82E-01 |
| *B. sp. K4410.MGS-46* | 2.30E-01 | 5.95E-01 | 1.54E-01 | 2.83E-01 | 6.94E-01 | 8.64E-01 |
| *C. hylemonae* | 1.22E-01 | 5.18E-01 | 4.25E-01 | 5.45E-01 | 4.31E-01 | 8.27E-01 |
| Erysipelotrichaceae(OTU96) | 1.62E-01 | 5.37E-01 | 6.61E-07 | 8.39E-05 | 5.93E-04 | 5.54E-02 |
| *F. prausnitzii(OTU92)* | 4.05E-01 | 6.94E-01 | 4.90E-02 | 1.78E-01 | 4.49E-01 | 8.27E-01 |
| *E. ramosum(OTU89)* | 4.62E-01 | 7.05E-01 | 9.29E-02 | 2.41E-01 | 4.31E-01 | 8.27E-01 |
| *B. hydrogenotrophica(OTU118)* | 1.48E-01 | 5.37E-01 | 2.77E-01 | 4.05E-01 | 8.80E-01 | 9.39E-01 |
| *L. phytofermentans* | 7.50E-01 | 8.50E-01 | 2.63E-01 | 4.02E-01 | 2.38E-01 | 7.75E-01 |

| | | | | | | |
|---|---|---|---|---|---|---|
| *C. xylanolyticum* | 1.94E-01 | 5.59E-01 | 9.13E-02 | 2.41E-01 | 9.76E-01 | 9.91E-01 |
| *E. ramosum(OTU77)* | 7.87E-01 | 8.85E-01 | 1.42E-02 | 9.90E-02 | 4.92E-02 | 4.79E-01 |
| *A. glycaniphila(OTU81)* | 9.22E-01 | 9.52E-01 | 5.96E-02 | 2.03E-01 | 1.30E-01 | 6.13E-01 |
| *E. oxidoreducens* | 1.78E-01 | 5.37E-01 | 2.90E-03 | 5.14E-02 | 7.42E-02 | 5.81E-01 |
| *B. producta* | 4.18E-01 | 6.99E-01 | 1.29E-01 | 2.68E-01 | 5.86E-01 | 8.40E-01 |
| *R. hungatei(OTU107)* | 3.27E-01 | 6.48E-01 | 3.42E-01 | 4.72E-01 | 9.04E-01 | 9.48E-01 |
| *R. gnavus(OTU97)* | 9.80E-01 | 9.88E-01 | 2.48E-01 | 3.90E-01 | 2.96E-01 | 7.77E-01 |
| *B. barnesiae(OTU102)* | 1.74E-01 | 5.37E-01 | 5.30E-01 | 6.41E-01 | 6.94E-01 | 8.64E-01 |
| *B. hydrogenotrophica(OTU124)* | 1.08E-02 | 2.78E-01 | 5.30E-02 | 1.87E-01 | 3.11E-01 | 7.77E-01 |
| *A. naeslundii* | 4.77E-01 | 7.05E-01 | 1.64E-01 | 2.89E-01 | 2.38E-01 | 7.75E-01 |
| *E. caecimuris* | 2.91E-02 | 2.84E-01 | 2.38E-03 | 5.14E-02 | 7.51E-01 | 8.99E-01 |
| *E. rectale(OTU12)* | 1.80E-02 | 2.78E-01 | 3.23E-03 | 5.14E-02 | 5.35E-01 | 8.40E-01 |
| *L. multipara(OTU37)* | 3.98E-01 | 6.94E-01 | 1.01E-02 | 8.58E-02 | 1.19E-01 | 5.82E-01 |
| *A. glycaniphila(OTU110)* | 6.59E-01 | 8.19E-01 | 6.96E-01 | 7.76E-01 | 4.77E-01 | 8.40E-01 |
| *CAG-352(OTU123)* | 6.77E-01 | 8.19E-01 | 1.31E-01 | 2.68E-01 | 3.80E-01 | 8.27E-01 |
| Lachnospiraceae(OTU4) | 4.77E-01 | 7.05E-01 | 2.56E-01 | 3.96E-01 | 6.28E-01 | 8.40E-01 |
| *P. psychrophila* | 4.26E-01 | 7.01E-01 | 3.09E-01 | 4.35E-01 | 9.01E-02 | 5.81E-01 |
| Ruminococcaceae(OTU52) | 6.07E-01 | 7.86E-01 | 1.80E-01 | 3.05E-01 | 3.64E-01 | 8.27E-01 |
| *F. prausnitzii(OTU61)* | 3.03E-01 | 6.21E-01 | 8.53E-03 | 8.33E-02 | 2.04E-01 | 7.37E-01 |
| *B. ihuae(OTU101)* | 2.39E-01 | 6.08E-01 | 7.48E-02 | 2.11E-01 | 5.86E-01 | 8.40E-01 |
| *L. pacaense(OTU93)* | 8.39E-02 | 4.44E-01 | 1.31E-01 | 2.68E-01 | 5.45E-01 | 8.40E-01 |

| | | | | | | |
|---|---|---|---|---|---|---|
| *R. sp. N15.MGS-57* | 4.70E-02 | 3.32E-01 | 1.38E-01 | 2.69E-01 | 4.86E-01 | 8.40E-01 |
| *A. indistinctus(OTU108)* | 1.52E-01 | 5.37E-01 | 1.47E-01 | 2.74E-01 | 7.97E-01 | 9.01E-01 |
| *S. lutetiensis* | 7.50E-01 | 8.50E-01 | 2.62E-02 | 1.16E-01 | 9.60E-02 | 5.81E-01 |
| *T. sanguinis* | 5.48E-01 | 7.48E-01 | 7.54E-01 | 8.25E-01 | 4.40E-01 | 8.27E-01 |
| Ruminococcaceae(OTU25) | 1.99E-02 | 2.78E-01 | 1.44E-01 | 2.74E-01 | 2.63E-01 | 7.77E-01 |
| *Ruminiclostridium(OTU90)* | 5.24E-01 | 7.39E-01 | 2.42E-01 | 3.90E-01 | 7.85E-01 | 8.99E-01 |
| *UBA1819(OTU105)* | 1.35E-01 | 5.37E-01 | 3.68E-01 | 4.92E-01 | 4.31E-01 | 8.27E-01 |
| *K. alysoides* | 4.47E-01 | 7.01E-01 | 6.90E-01 | 7.76E-01 | 6.28E-01 | 8.40E-01 |
| *B. hydrogenotrophica(OTU7)* | 3.03E-01 | 6.21E-01 | 2.22E-01 | 3.66E-01 | 7.62E-01 | 8.99E-01 |
| *B. hansenii(OTU11)* | 7.31E-01 | 8.50E-01 | 3.68E-01 | 4.92E-01 | 5.65E-01 | 8.40E-01 |
| Ruminococcaceae(OTU115) | 1.22E-01 | 5.18E-01 | 2.01E-02 | 1.16E-01 | 5.65E-01 | 8.40E-01 |
| *R. lactatiformans* | 3.52E-01 | 6.57E-01 | 6.71E-01 | 7.61E-01 | 8.09E-01 | 9.01E-01 |
| *P. secunda* | 2.30E-01 | 5.95E-01 | 1.14E-01 | 2.60E-01 | 5.15E-01 | 8.40E-01 |
| *C. cystitidis* | 8.54E-01 | 9.19E-01 | 1.75E-01 | 3.04E-01 | 2.76E-01 | 7.77E-01 |
| *S. pseudolugdunensis* | 4.77E-01 | 7.05E-01 | 4.76E-01 | 5.87E-01 | 2.76E-01 | 7.77E-01 |
| *Bilophila(OTU79)* | 2.81E-01 | 6.21E-01 | 4.34E-02 | 1.67E-01 | 6.50E-01 | 8.42E-01 |
| *R. inulinivorans* | 3.50E-02 | 3.17E-01 | 3.46E-02 | 1.37E-01 | 5.86E-01 | 8.40E-01 |
| Ruminococcaceae(OTU44) | 9.07E-02 | 4.61E-01 | 3.15E-03 | 5.14E-02 | 2.15E-01 | 7.37E-01 |
| *R. bromii* | 8.45E-01 | 9.17E-01 | 6.29E-01 | 7.19E-01 | 5.45E-01 | 8.40E-01 |

| | | | | | | |
|---|---|---|---|---|---|---|
| *R. hungatei(OTU63)* | 3.82E-03 | 2.78E-01 | 5.97E-03 | 6.89E-02 | 5.25E-01 | 8.40E-01 |
| *B. cucumis* | 3.39E-01 | 6.57E-01 | 2.51E-02 | 1.16E-01 | 4.49E-01 | 8.27E-01 |
| *D. pneumosintes* | 5.32E-01 | 7.42E-01 | 8.92E-01 | 9.06E-01 | 6.94E-01 | 8.64E-01 |
| *Ruminiclostridium(OTU 62)* | 1.55E-01 | 5.37E-01 | 6.94E-02 | 2.09E-01 | 9.04E-01 | 9.48E-01 |
| *R. hominis(OTU29)* | 6.68E-01 | 8.19E-01 | 3.87E-01 | 5.06E-01 | 6.50E-01 | 8.42E-01 |
| *B. magnum* | 1.74E-02 | 2.78E-01 | 3.25E-02 | 1.33E-01 | 8.32E-01 | 9.19E-01 |
| *I. bartlettii* | 1.63E-02 | 2.78E-01 | 3.13E-01 | 4.36E-01 | 8.47E-03 | 1.79E-01 |
| *P. chartae(OTU125)* | 1.03E-01 | 4.85E-01 | 3.46E-01 | 4.73E-01 | 3.40E-01 | 8.16E-01 |
| *H. effluvii* | 4.33E-01 | 7.01E-01 | 3.78E-01 | 4.99E-01 | 1.93E-01 | 7.22E-01 |
| *C. faecale* | 3.03E-01 | 6.21E-01 | 8.65E-01 | 9.00E-01 | 4.14E-01 | 8.27E-01 |
| *A. senegalensis* | 8.45E-01 | 9.17E-01 | 8.79E-01 | 9.00E-01 | 1.00E+00 | 1.00E+00 |
| *P. chartae(OTU70)* | 2.12E-02 | 2.78E-01 | 1.09E-01 | 2.60E-01 | 4.14E-01 | 8.27E-01 |
| *Ezakiella(OTU53)* | 5.24E-01 | 7.39E-01 | 8.39E-01 | 8.80E-01 | 3.97E-01 | 8.27E-01 |
| Ruminococcaceae(OTU7 4) | 2.26E-02 | 2.78E-01 | 4.71E-02 | 1.76E-01 | 9.64E-01 | 9.87E-01 |
| *T. nexilis* | 2.81E-01 | 6.21E-01 | 6.05E-01 | 7.11E-01 | 6.07E-01 | 8.40E-01 |
| *L. pacaense(OTU122)* | 9.22E-01 | 9.52E-01 | 1.86E-01 | 3.10E-01 | 1.93E-01 | 7.22E-01 |
| *B. sp. YHC-4* | 8.83E-01 | 9.27E-01 | 9.46E-01 | 9.53E-01 | 8.68E-01 | 9.39E-01 |
| *B. xylanolyticus* | 1.03E-01 | 4.85E-01 | 1.14E-02 | 9.09E-02 | 3.18E-01 | 7.77E-01 |
| *S. variabile(OTU104)* | 1.59E-01 | 5.37E-01 | 1.25E-01 | 2.68E-01 | 3.72E-01 | 8.27E-01 |
| *A. ruminis* | 7.31E-01 | 8.50E-01 | 1.07E-01 | 2.60E-01 | 5.45E-01 | 8.40E-01 |
| *C. innocuum* | 4.47E-01 | 7.01E-01 | 1.33E-01 | 2.68E-01 | 3.18E-01 | 7.77E-01 |
| *V. criceti* | 4.47E-01 | 7.01E-01 | 2.93E-01 | 4.18E-01 | 2.69E-01 | 7.77E-01 |

| | | | | | | |
|---|---|---|---|---|---|---|
| Lachnospiraceae(OTU86) | 6.59E-02 | 3.99E-01 | 1.12E-03 | 4.25E-02 | 3.96E-02 | 4.68E-01 |
| *S. thermophilus TH1435* | 4.18E-01 | 6.99E-01 | 8.81E-02 | 2.41E-01 | 5.86E-01 | 8.40E-01 |
| *R. lactaris(OTU83)* | 1.47E-02 | 2.78E-01 | 9.59E-01 | 9.59E-01 | 4.26E-02 | 4.68E-01 |
| *E. ramulus(OTU80)* | 7.04E-01 | 8.36E-01 | 3.91E-01 | 5.07E-01 | 7.28E-01 | 8.89E-01 |
| Ruminococcaceae(OTU75) | 6.24E-02 | 3.96E-01 | 7.48E-02 | 2.11E-01 | 9.88E-01 | 9.96E-01 |
| *O. hirsuta* | 8.25E-01 | 9.17E-01 | 2.70E-01 | 4.03E-01 | 6.72E-01 | 8.62E-01 |
| Lachnospiraceae(OTU68) | 9.32E-01 | 9.54E-01 | 1.87E-02 | 1.13E-01 | 4.42E-02 | 4.68E-01 |
| *A. equolifaciens* | 1.59E-01 | 5.37E-01 | 9.65E-03 | 8.58E-02 | 3.56E-01 | 8.27E-01 |
| *B. coprophilus* | 4.70E-02 | 3.32E-01 | 4.45E-01 | 5.65E-01 | 2.50E-01 | 7.77E-01 |
| Christensenellaceae(OTU59) | 2.81E-01 | 6.21E-01 | 6.43E-02 | 2.03E-01 | 3.04E-01 | 7.77E-01 |
| *L. pacaense(OTU54)* | 5.81E-01 | 7.72E-01 | 6.56E-02 | 2.03E-01 | 2.09E-01 | 7.37E-01 |
| *F. plautii* | 3.52E-01 | 6.57E-01 | 2.70E-01 | 4.03E-01 | 3.55E-02 | 4.68E-01 |
| *E. rectale(OTU51)* | 2.92E-01 | 6.21E-01 | 1.22E-01 | 2.68E-01 | 8.45E-02 | 5.81E-01 |
| *A. sp. AL-1* | 2.91E-02 | 2.84E-01 | 1.56E-02 | 9.90E-02 | 9.28E-01 | 9.58E-01 |
| *R. gnavus(OTU47)* | 3.91E-01 | 6.94E-01 | 5.64E-01 | 6.76E-01 | 1.83E-01 | 7.22E-01 |
| *C. comes* | 5.90E-01 | 7.72E-01 | 4.81E-01 | 5.88E-01 | 8.56E-01 | 9.37E-01 |
| *R. albus* | 8.64E-01 | 9.22E-01 | 1.18E-01 | 2.64E-01 | 1.69E-01 | 6.90E-01 |
| *B. barnesiae(OTU40)* | 1.52E-01 | 5.37E-01 | 8.25E-01 | 8.74E-01 | 1.16E-01 | 5.82E-01 |
| *B. paurosaccharolyticus* | 4.44E-02 | 3.32E-01 | 1.35E-01 | 2.69E-01 | 5.35E-01 | 8.40E-01 |
| *B. coprosuis* | 2.81E-01 | 6.21E-01 | 4.76E-01 | 5.87E-01 | 7.85E-01 | 8.99E-01 |
| *S. variabile(OTU33)* | 4.92E-01 | 7.11E-01 | 8.79E-01 | 9.00E-01 | 7.85E-01 | 8.99E-01 |

| | | | | | | |
|---|---|---|---|---|---|---|
| *E. hermanniensis* | 3.09E-01 | 6.23E-01 | 5.99E-01 | 7.11E-01 | 6.28E-01 | 8.40E-01 |
| *C. tanakaei* | 2.54E-01 | 6.21E-01 | 1.77E-01 | 3.04E-01 | 1.69E-01 | 6.90E-01 |
| *E. rectale(OTU30)* | 2.49E-01 | 6.21E-01 | 2.48E-01 | 3.90E-01 | 8.45E-02 | 5.81E-01 |
| *A. indistinctus(OTU26)* | 1.74E-02 | 2.78E-01 | 1.34E-03 | 4.25E-02 | 5.45E-01 | 8.40E-01 |
| Lachnospiraceae(OUT23) | 3.03E-01 | 6.21E-01 | 1.14E-01 | 2.60E-01 | 7.17E-01 | 8.83E-01 |
| *O. jeddahense* | 2.41E-02 | 2.78E-01 | 7.99E-01 | 8.53E-01 | 7.08E-03 | 1.79E-01 |
| *R. gauvreauii* | 7.41E-01 | 8.50E-01 | 6.29E-01 | 7.19E-01 | 4.40E-01 | 8.27E-01 |
| *C. europaeus* | 6.86E-01 | 8.22E-01 | 2.10E-02 | 1.16E-01 | 7.08E-03 | 1.79E-01 |
| *B. ihuae(OTU16)* | 1.70E-01 | 5.37E-01 | 2.62E-02 | 1.16E-01 | 8.80E-01 | 9.39E-01 |
| *C. autoethanogenum* | 4.92E-01 | 7.11E-01 | 1.56E-02 | 9.90E-02 | 2.72E-02 | 4.31E-01 |
| *E. tayi* | 7.75E-02 | 4.28E-01 | 7.22E-01 | 7.97E-01 | 2.56E-01 | 7.77E-01 |
| *L. multipara(OTU6)* | 8.45E-01 | 9.17E-01 | 1.14E-01 | 2.60E-01 | 1.46E-01 | 6.41E-01 |
| *D. longicatena* | 1.94E-01 | 5.59E-01 | 2.74E-02 | 1.16E-01 | 8.47E-03 | 1.79E-01 |
| *E. sinensis* | 6.50E-01 | 8.19E-01 | 2.85E-01 | 4.11E-01 | 8.09E-01 | 9.01E-01 |
| *Ruminiclostridium(OTU2)* | 7.75E-02 | 4.28E-01 | 1.47E-01 | 2.74E-01 | 7.39E-01 | 8.94E-01 |
| *B. graminisolvens* | 3.83E-02 | 3.24E-01 | 1.61E-01 | 2.89E-01 | 4.58E-01 | 8.32E-01 |
| *P. nigrescens* | 8.83E-01 | 9.27E-01 | 7.99E-01 | 8.53E-01 | 6.50E-01 | 8.42E-01 |
| *T. glycolicus* | 5.90E-01 | 7.72E-01 | 6.43E-02 | 2.03E-01 | 5.45E-01 | 8.40E-01 |
| *O. valericigenes* | 2.59E-01 | 6.21E-01 | 5.38E-03 | 6.83E-02 | 2.96E-01 | 7.77E-01 |
| *B. barnesiae(OTU73)* | 4.98E-02 | 3.33E-01 | 1.14E-01 | 2.60E-01 | 6.07E-01 | 8.40E-01 |
| Lachnospiraceae(OTU42) | 3.77E-01 | 6.94E-01 | 8.11E-03 | 8.33E-02 | 1.02E-01 | 5.82E-01 |
| *P. soli* | 4.77E-01 | 7.05E-01 | 2.48E-01 | 3.90E-01 | 1.16E-01 | 5.82E-01 |

| | | | | | | |
|---|---|---|---|---|---|---|
| *R. lactaris(OTU117)* | 1.17E-01 | 5.18E-01 | 1.52E-02 | 9.90E-02 | 6.28E-01 | 8.40E-01 |
| *Escherichia-Shigella(OTU72)* | 1.70E-01 | 5.37E-01 | 9.29E-02 | 2.41E-01 | 2.32E-02 | 4.21E-01 |
| *R. torques* | 9.61E-01 | 9.76E-01 | 2.30E-02 | 1.16E-01 | 6.94E-02 | 5.81E-01 |
| *L. hircilactis* | 5.90E-01 | 7.72E-01 | 2.74E-02 | 1.16E-01 | 5.28E-02 | 4.79E-01 |

Cont.

| Abb. | HITS score | | | | | |
|---|---|---|---|---|---|---|
| | normal_hits_score | normal_hits_pval | nonaf_hits_score | nonaf_hits_pval | cirrhosis_hits_score | cirrhosis_hits_pval |
| *D. formicigenerans* | 0.041 | 1.00E-03 | 0.033 | 2.20E-02 | 0.049 | 1.00E-03 |
| *H. porcina* | 0.049 | 1.00E-03 | 0.007 | 4.10E-01 | 0.042 | 1.00E-03 |
| *AD3011(OTU78)* | 0.001 | 9.98E-01 | 0.005 | 5.17E-01 | 0.037 | 1.00E-03 |
| *B. sp. Marseille-P3087* | 0.025 | 1.00E-03 | 0.013 | 1.70E-01 | 0.047 | 1.00E-03 |
| *R. hominis(OTU106)* | 0.003 | 9.41E-01 | 0.005 | 5.13E-01 | 0.002 | 9.44E-01 |
| *R. lituseburensis* | 0.019 | 1.70E-02 | 0.003 | 6.47E-01 | 0.005 | 6.39E-01 |
| *E. ramulus(OTU103)* | 0.031 | 1.00E-03 | 0.000 | 1.00E+00 | 0.000 | 1.00E+00 |
| *A. caccae* | 0.020 | 6.00E-03 | 0.000 | 1.00E+00 | 0.001 | 9.83E-01 |
| *F. saccharivorans* | 0.031 | 1.00E-03 | 0.000 | 1.00E+00 | 0.004 | 7.70E-01 |
| *B. hansenii(OTU14)* | 0.014 | 5.30E-02 | 0.035 | 1.50E-02 | 0.031 | 3.00E-03 |
| *C. eutactus* | 0.016 | 3.80E-02 | 0.020 | 9.10E-02 | 0.001 | 9.77E-01 |
| *B. sp. K4410.MGS-46* | 0.007 | 5.21E-01 | 0.004 | 6.23E-01 | 0.007 | 5.53E-01 |
| *C. hylemonae* | 0.008 | 4.66E-01 | 0.001 | 8.43E-01 | 0.000 | 9.88E-01 |
| Erysipelotrichaceae(OTU96) | 0.026 | 1.00E-03 | 0.028 | 2.80E-02 | 0.002 | 8.85E-01 |
| *F. prausnitzii(OTU92)* | 0.002 | 9.65E-01 | 0.000 | 1.00E+00 | 0.001 | 9.84E-01 |
| *E. ramosum(OTU89)* | 0.021 | 2.00E-03 | 0.001 | 8.51E-01 | 0.000 | 9.84E-01 |
| *B. hydrogenotrophica(OTU118)* | 0.005 | 8.02E-01 | 0.004 | 5.84E-01 | 0.011 | 2.25E-01 |
| *L. phytofermentans* | 0.001 | 9.87E-01 | 0.003 | 6.36E-01 | 0.034 | 1.00E-03 |
| *C. xylanolyticum* | 0.003 | 9.10E-01 | 0.036 | 1.90E-02 | 0.002 | 8.82E-01 |
| *E. ramosum(OTU77)* | 0.017 | 1.80E-02 | 0.000 | 1.00E+00 | 0.003 | 8.70E-01 |
| *A. glycaniphila(OTU81)* | 0.015 | 3.70E-02 | 0.001 | 8.12E-01 | 0.000 | 1.00E+00 |
| *E. oxidoreducens* | 0.011 | 2.07E-01 | 0.007 | 4.22E-01 | 0.000 | 9.84E-01 |

| | | | | | | |
|---|---|---|---|---|---|---|
| *B. producta* | 0.032 | 1.00E-03 | 0.026 | 3.90E-02 | 0.041 | 1.00E-03 |
| *R. hungatei(OTU107)* | 0.006 | 6.09E-01 | 0.002 | 7.84E-01 | 0.001 | 9.80E-01 |
| *R. gnavus(OTU97)* | 0.006 | 6.21E-01 | 0.000 | 1.00E+00 | 0.002 | 9.02E-01 |
| *B. barnesiae(OTU102)* | 0.006 | 7.07E-01 | 0.021 | 8.20E-02 | 0.022 | 1.00E-02 |
| *B. hydrogenotrophica(OTU124)* | 0.011 | 2.05E-01 | 0.007 | 4.13E-01 | 0.000 | 1.00E+00 |
| *A. naeslundii* | 0.007 | 5.06E-01 | 0.023 | 5.80E-02 | 0.017 | 4.30E-02 |
| *E. caecimuris* | 0.000 | 1.00E+00 | 0.002 | 8.16E-01 | 0.024 | 6.00E-03 |
| *E. rectale(OTU12)* | 0.006 | 6.34E-01 | 0.001 | 8.26E-01 | 0.001 | 9.74E-01 |
| *L. multipara(OTU37)* | 0.001 | 9.92E-01 | 0.009 | 3.53E-01 | 0.000 | 9.92E-01 |
| *A. glycaniphila(OTU110)* | 0.000 | 1.00E+00 | 0.010 | 2.79E-01 | 0.002 | 9.26E-01 |
| *CAG-352(OTU123)* | 0.002 | 9.74E-01 | 0.000 | 1.00E+00 | 0.000 | 9.91E-01 |
| Lachnospiraceae(OTU4) | 0.004 | 8.19E-01 | 0.000 | 1.00E+00 | 0.030 | 3.00E-03 |
| *P. psychrophila* | 0.012 | 1.67E-01 | 0.007 | 4.31E-01 | 0.005 | 6.17E-01 |
| Ruminococcaceae(OTU52) | 0.008 | 4.29E-01 | 0.000 | 8.98E-01 | 0.001 | 9.73E-01 |
| *F. prausnitzii(OTU61)* | 0.008 | 4.32E-01 | 0.014 | 1.79E-01 | 0.001 | 9.61E-01 |
| *B. ihuae(OTU101)* | 0.007 | 5.06E-01 | 0.056 | 6.00E-03 | 0.001 | 9.78E-01 |
| *L. pacaense(OTU93)* | 0.003 | 9.23E-01 | 0.001 | 8.29E-01 | 0.001 | 9.81E-01 |
| *R. sp. N15.MGS-57* | 0.002 | 9.57E-01 | 0.001 | 8.22E-01 | 0.000 | 9.80E-01 |
| *A. indistinctus(OTU108)* | 0.006 | 6.53E-01 | 0.013 | 2.11E-01 | 0.000 | 1.00E+00 |
| *S. lutetiensis* | 0.003 | 9.16E-01 | 0.002 | 7.16E-01 | 0.001 | 9.83E-01 |
| *T. sanguinis* | 0.003 | 9.24E-01 | 0.004 | 5.95E-01 | 0.000 | 9.82E-01 |
| Ruminococcaceae(OTU25) | 0.003 | 9.15E-01 | 0.012 | 2.26E-01 | 0.001 | 9.59E-01 |
| *Ruminiclostridium(OTU90)* | 0.003 | 9.00E-01 | 0.005 | 5.14E-01 | 0.002 | 9.06E-01 |
| *UBA1819(OTU105)* | 0.003 | 8.89E-01 | 0.001 | 8.41E-01 | 0.002 | 8.89E-01 |
| *K. alysoides* | 0.006 | 6.64E-01 | 0.002 | 7.56E-01 | 0.010 | 2.91E-01 |

| | | | | | | |
|---|---|---|---|---|---|---|
| *B. hydrogenotrophica(OTU7)* | 0.017 | 1.80E-02 | 0.004 | 5.70E-01 | 0.003 | 8.85E-01 |
| *B. hansenii(OTU11)* | 0.038 | 1.00E-03 | 0.004 | 5.93E-01 | 0.017 | 5.70E-02 |
| Ruminococcaceae(OTU115) | 0.003 | 9.39E-01 | 0.001 | 8.34E-01 | 0.000 | 9.92E-01 |
| *R. lactatiformans* | 0.002 | 9.77E-01 | 0.000 | 8.98E-01 | 0.011 | 2.23E-01 |
| *P. secunda* | 0.004 | 8.87E-01 | 0.000 | 1.00E+00 | 0.000 | 1.00E+00 |
| *C. cystitidis* | 0.001 | 9.93E-01 | 0.000 | 1.00E+00 | 0.000 | 9.86E-01 |
| *S. pseudolugdunensis* | 0.000 | 9.97E-01 | 0.000 | 1.00E+00 | 0.001 | 9.85E-01 |
| *Bilophila(OTU79)* | 0.003 | 9.25E-01 | 0.002 | 7.88E-01 | 0.000 | 1.00E+00 |
| *R. inulinivorans* | 0.003 | 9.36E-01 | 0.012 | 2.32E-01 | 0.001 | 9.88E-01 |
| Ruminococcaceae(OTU44) | 0.002 | 9.81E-01 | 0.011 | 2.80E-01 | 0.001 | 9.79E-01 |
| *R. bromii* | 0.005 | 7.81E-01 | 0.027 | 3.40E-02 | 0.005 | 6.51E-01 |
| *R. hungatei(OTU63)* | 0.002 | 9.75E-01 | 0.005 | 5.06E-01 | 0.001 | 9.83E-01 |
| *B. cucumis* | 0.001 | 9.96E-01 | 0.000 | 1.00E+00 | 0.000 | 1.00E+00 |
| *D. pneumosintes* | 0.001 | 9.98E-01 | 0.011 | 2.53E-01 | 0.001 | 9.53E-01 |
| *Ruminiclostridium(OTU62)* | 0.001 | 9.98E-01 | 0.004 | 5.77E-01 | 0.000 | 9.83E-01 |
| *R. hominis(OTU29)* | 0.000 | 9.97E-01 | 0.000 | 1.00E+00 | 0.001 | 9.77E-01 |
| *B. magnum* | 0.001 | 1.00E+00 | 0.002 | 7.91E-01 | 0.001 | 9.79E-01 |
| *I. bartlettii* | 0.001 | 1.00E+00 | 0.000 | 1.00E+00 | 0.000 | 9.79E-01 |
| *P. chartae(OTU125)* | 0.001 | 9.97E-01 | 0.009 | 3.03E-01 | 0.000 | 9.90E-01 |
| *H. effluvii* | 0.000 | 9.98E-01 | 0.010 | 2.74E-01 | 0.023 | 1.20E-02 |
| *C. faecale* | 0.000 | 9.96E-01 | 0.006 | 4.71E-01 | 0.000 | 9.92E-01 |
| *A. senegalensis* | 0.000 | 9.98E-01 | 0.003 | 6.38E-01 | 0.000 | 1.00E+00 |
| *P. chartae(OTU70)* | 0.000 | 9.99E-01 | 0.007 | 4.32E-01 | 0.000 | 1.00E+00 |
| *Ezakiella(OTU53)* | 0.000 | 1.00E+00 | 0.000 | 1.00E+00 | 0.002 | 9.34E-01 |
| Ruminococcaceae(OTU74) | 0.002 | 9.69E-01 | 0.012 | 2.14E-01 | 0.026 | 7.00E-03 |

| | | | | | | |
|---|---|---|---|---|---|---|
| *T. nexilis* | 0.000 | 9.99E-01 | 0.002 | 7.47E-01 | 0.001 | 9.70E-01 |
| *L. pacaense(OTU122)* | 0.018 | 1.50E-02 | 0.012 | 2.14E-01 | 0.002 | 9.33E-01 |
| *B. sp. YHC-4* | 0.009 | 2.93E-01 | 0.000 | 1.00E+00 | 0.004 | 7.81E-01 |
| *B. xylanolyticus* | 0.004 | 8.44E-01 | 0.004 | 6.13E-01 | 0.001 | 9.78E-01 |
| *S. variabile(OTU104)* | 0.002 | 9.69E-01 | 0.022 | 6.50E-02 | 0.055 | 1.00E-03 |
| *A. ruminis* | 0.006 | 6.62E-01 | 0.004 | 6.02E-01 | 0.000 | 1.00E+00 |
| *C. innocuum* | 0.006 | 6.97E-01 | 0.002 | 7.81E-01 | 0.018 | 3.70E-02 |
| *V. criceti* | 0.003 | 9.28E-01 | 0.014 | 1.77E-01 | 0.010 | 2.81E-01 |
| Lachnospiraceae(OTU86) | 0.004 | 8.17E-01 | 0.001 | 8.61E-01 | 0.003 | 8.11E-01 |
| *S. thermophilus TH1435* | 0.003 | 8.91E-01 | 0.001 | 8.33E-01 | 0.002 | 9.47E-01 |
| *R. lactaris(OTU83)* | 0.006 | 6.42E-01 | 0.001 | 8.58E-01 | 0.007 | 4.77E-01 |
| *E. ramulus(OTU80)* | 0.002 | 9.38E-01 | 0.001 | 8.58E-01 | 0.000 | 9.91E-01 |
| Ruminococcaceae(OTU75) | 0.008 | 4.74E-01 | 0.009 | 3.07E-01 | 0.000 | 9.88E-01 |
| *O. hirsuta* | 0.000 | 1.00E+00 | 0.025 | 5.50E-02 | 0.003 | 8.67E-01 |
| Lachnospiraceae(OTU68) | 0.004 | 8.57E-01 | 0.007 | 3.87E-01 | 0.001 | 9.80E-01 |
| *A. equolifaciens* | 0.013 | 8.40E-02 | 0.015 | 1.69E-01 | 0.013 | 1.75E-01 |
| *B. coprophilus* | 0.001 | 9.84E-01 | 0.024 | 5.00E-02 | 0.010 | 2.59E-01 |
| Christensenellaceae(OTU59) | 0.011 | 2.39E-01 | 0.006 | 4.50E-01 | 0.015 | 9.80E-02 |
| *L. pacaense(OTU54)* | 0.011 | 1.77E-01 | 0.005 | 5.56E-01 | 0.000 | 9.86E-01 |
| *F. plautii* | 0.001 | 9.94E-01 | 0.037 | 1.60E-02 | 0.039 | 1.00E-03 |
| *E. rectale(OTU51)* | 0.023 | 2.00E-03 | 0.008 | 3.58E-01 | 0.034 | 2.00E-03 |
| *A. sp. AL-1* | 0.001 | 9.83E-01 | 0.001 | 8.91E-01 | 0.000 | 9.86E-01 |
| *R. gnavus(OTU47)* | 0.028 | 1.00E-03 | 0.029 | 1.80E-02 | 0.019 | 3.40E-02 |
| *C. comes* | 0.023 | 3.00E-03 | 0.012 | 2.45E-01 | 0.004 | 8.10E-01 |
| *R. albus* | 0.003 | 9.20E-01 | 0.007 | 4.00E-01 | 0.000 | 9.94E-01 |

| | | | | | | |
|---|---|---|---|---|---|---|
| *B. barnesiae(OTU40)* | 0.006 | 6.81E-01 | 0.007 | 4.00E-01 | 0.000 | 9.84E-01 |
| *B. paurosaccharolyticus* | 0.002 | 9.72E-01 | 0.011 | 2.39E-01 | 0.000 | 9.87E-01 |
| *B. coprosuis* | 0.003 | 9.37E-01 | 0.019 | 9.90E-02 | 0.002 | 8.92E-01 |
| *S. variabile(OTU33)* | 0.003 | 9.29E-01 | 0.005 | 4.84E-01 | 0.021 | 1.60E-02 |
| *E. hermanniensis* | 0.017 | 1.50E-02 | 0.007 | 4.21E-01 | 0.001 | 9.58E-01 |
| *C. tanakaei* | 0.007 | 5.94E-01 | 0.000 | 8.70E-01 | 0.025 | 1.20E-02 |
| *E. rectale(OTU30)* | 0.028 | 2.00E-03 | 0.016 | 1.22E-01 | 0.037 | 2.00E-03 |
| *A. indistinctus(OTU26)* | 0.003 | 8.83E-01 | 0.000 | 8.82E-01 | 0.001 | 9.73E-01 |
| Lachnospiraceae(OUT23) | 0.000 | 1.00E+00 | 0.000 | 8.78E-01 | 0.001 | 9.81E-01 |
| *O. jeddahense* | 0.000 | 1.00E+00 | 0.000 | 9.39E-01 | 0.000 | 9.91E-01 |
| *R. gauvreauii* | 0.000 | 9.99E-01 | 0.000 | 1.00E+00 | 0.012 | 1.78E-01 |
| *C. europaeus* | 0.004 | 8.20E-01 | 0.000 | 9.00E-01 | 0.005 | 7.09E-01 |
| *B. ihuae(OTU16)* | 0.008 | 3.99E-01 | 0.009 | 3.28E-01 | 0.001 | 9.72E-01 |
| *C. autoethanogenum* | 0.000 | 1.00E+00 | 0.003 | 6.83E-01 | 0.001 | 9.69E-01 |
| *E. tayi* | 0.004 | 8.21E-01 | 0.000 | 1.00E+00 | 0.000 | 9.83E-01 |
| *L. multipara(OTU6)* | 0.000 | 1.00E+00 | 0.033 | 2.50E-02 | 0.000 | 9.88E-01 |
| *D. longicatena* | 0.042 | 1.00E-03 | 0.021 | 9.90E-02 | 0.042 | 1.00E-03 |
| *E. sinensis* | 0.018 | 1.60E-02 | 0.003 | 6.63E-01 | 0.000 | 9.85E-01 |
| *Ruminiclostridium(OTU2)* | 0.012 | 1.67E-01 | 0.003 | 6.42E-01 | 0.005 | 6.77E-01 |
| *B. graminisolvens* | 0.002 | 9.91E-01 | 0.005 | 5.31E-01 | 0.000 | 9.87E-01 |
| *P. nigrescens* | 0.001 | 9.94E-01 | 0.011 | 2.57E-01 | 0.003 | 8.50E-01 |
| *T. glycolicus* | 0.025 | 2.00E-03 | 0.002 | 7.55E-01 | 0.000 | 9.84E-01 |
| *O. valericigenes* | 0.001 | 9.87E-01 | 0.011 | 2.43E-01 | 0.001 | 9.63E-01 |
| *B. barnesiae(OTU73)* | 0.001 | 9.92E-01 | 0.002 | 7.72E-01 | 0.004 | 7.32E-01 |
| Lachnospiraceae(OTU42) | 0.003 | 9.13E-01 | 0.000 | 1.00E+00 | 0.001 | 9.65E-01 |

| | | | | | | |
|---|---|---|---|---|---|---|
| *P. soli* | 0.001 | 9.82E-01 | 0.015 | 1.54E-01 | 0.000 | 9.85E-01 |
| *R. lactaris(OTU117)* | 0.003 | 8.98E-01 | 0.010 | 2.95E-01 | 0.008 | 4.09E-01 |
| *Escherichia-Shigella(OTU72)* | 0.003 | 9.41E-01 | 0.000 | 1.00E+00 | 0.010 | 2.62E-01 |
| *R. torques* | 0.007 | 5.42E-01 | 0.000 | 1.00E+00 | 0.033 | 2.00E-03 |
| *L. hircilactis* | 0.002 | 9.78E-01 | 0.004 | 5.72E-01 | 0.006 | 5.54E-01 |

Cont.

| Abb. | Intervention score | | | | | |
|---|---|---|---|---|---|---|
| | nonaf_intervention_score | nonaf_cumulative_intervention_score | nonaf_cumulative_intervention_rank | cirrhosis_intervention_score | cirrhosis_cumulative_intervention_score | cirrhosis_cumulative_intervention_rank |
| *D. formicigenerans* | 0.128 | 0.633 | 17 | 0.168 | 0.168 | 1 |
| *H. porcina* | 0.063 | 0.305 | 5 | 0.060 | 0.173 | 2 |
| *AD3011(OTU78)* | -0.032 | 0.944 | 85 | -0.014 | 0.238 | 3 |
| *B. sp. Marseille-P3087* | 0.002 | 0.944 | 84 | 0.050 | 0.326 | 4 |
| *R. hominis(OTU106)* | 0.003 | 0.906 | 50 | 0.034 | 0.383 | 5 |
| *R. lituseburensis* | 0.041 | 0.396 | 7 | 0.049 | 0.434 | 6 |
| *E. ramulus(OTU103)* | 0.031 | 0.552 | 12 | 0.031 | 0.464 | 7 |
| *A. caccae* | 0.020 | 0.653 | 18 | 0.029 | 0.494 | 8 |
| *F. saccharivorans* | 0.031 | 0.522 | 11 | 0.017 | 0.496 | 9 |
| *B. hansenii(OTU14)* | 0.195 | 0.944 | 92 | 0.180 | 0.525 | 10 |
| *C. eutactus* | 0.069 | 0.616 | 15 | 0.009 | 0.533 | 11 |
| *B. sp. K4410.MGS-46* | -0.012 | 0.768 | 27 | 0.005 | 0.562 | 12 |
| *C. hylemonae* | -0.004 | 0.716 | 23 | 0.026 | 0.588 | 13 |
| Erysipelotrichaceae(OTU96) | -0.017 | 0.944 | 80 | 0.004 | 0.613 | 14 |
| *F. prausnitzii(OTU92)* | 0.002 | 0.930 | 59 | 0.023 | 0.636 | 15 |
| *E. ramosum(OTU89)* | 0.018 | 0.670 | 19 | 0.020 | 0.656 | 16 |

| | | | | | | |
|---|---|---|---|---|---|---|
| *B. hydrogenotrophica(OTU118)* | 0.011 | 0.822 | 35 | 0.015 | 0.656 | 17 |
| *L. phytofermentans* | 0.001 | 0.944 | 112 | -0.161 | 0.671 | 18 |
| *C. xylanolyticum* | -0.045 | 0.491 | 10 | 0.008 | 0.691 | 19 |
| *E. ramosum(OTU77)* | 0.017 | 0.944 | 86 | 0.020 | 0.708 | 20 |
| *A. glycaniphila(OTU81)* | 0.003 | 0.754 | 26 | 0.015 | 0.723 | 21 |
| *E. oxidoreducens* | 0.017 | 0.688 | 20 | 0.023 | 0.737 | 22 |
| *B. producta* | 0.002 | 0.943 | 70 | -0.137 | 0.751 | 23 |
| *R. hungatei(OTU107)* | 0.007 | 0.852 | 39 | 0.017 | 0.762 | 24 |
| *R. gnavus(OTU97)* | 0.006 | 0.865 | 41 | -0.003 | 0.774 | 25 |
| *B. barnesiae(OTU102)* | 0.047 | 0.353 | 6 | -0.120 | 0.786 | 26 |
| *B. hydrogenotrophica(OTU124)* | 0.011 | 0.800 | 33 | 0.011 | 0.798 | 27 |
| *A. naeslundii* | -0.012 | 0.944 | 89 | 0.050 | 0.805 | 28 |
| *E. caecimuris* | 0.000 | 0.944 | 98 | -0.032 | 0.816 | 29 |
| *E. rectale(OTU12)* | 0.009 | 0.944 | 117 | 0.015 | 0.823 | 30 |
| *L. multipara(OTU37)* | -0.024 | 0.940 | 65 | 0.004 | 0.833 | 31 |
| *A. glycaniphila(OTU110)* | 0.066 | 0.944 | 79 | -0.018 | 0.842 | 32 |

| | | | | | | |
|---|---|---|---|---|---|---|
| *CAG-352(OTU123)* | 0.002 | 0.932 | 60 | 0.009 | 0.851 | 33 |
| Lachnospiraceae(OTU4) | 0.004 | 0.895 | 47 | -0.084 | 0.860 | 34 |
| *P. psychrophila* | -0.013 | 0.778 | 31 | 0.038 | 0.860 | 35 |
| Ruminococcaceae(OTU52) | -0.002 | 0.779 | 29 | -0.003 | 0.868 | 36 |
| *F. prausnitzii(OTU61)* | 0.033 | 0.839 | 37 | 0.032 | 0.876 | 37 |
| *B. ihuae(OTU101)* | 0.015 | 0.424 | 8 | 0.021 | 0.883 | 38 |
| *L. pacaense(OTU93)* | -0.003 | 0.882 | 44 | -0.032 | 0.886 | 39 |
| *R. sp. N15.MGS-57* | 0.007 | 0.812 | 34 | 0.006 | 0.893 | 40 |
| *A. indistinctus(OTU108)* | -0.018 | 0.739 | 25 | 0.006 | 0.899 | 41 |
| *S. lutetiensis* | 0.031 | 0.944 | 119 | -0.034 | 0.899 | 42 |
| *T. sanguinis* | 0.012 | 0.781 | 28 | 0.000 | 0.905 | 43 |
| Ruminococcaceae(OTU25) | -0.014 | 0.944 | 110 | 0.014 | 0.910 | 44 |
| *Ruminiclostridium(OTU90)* | 0.011 | 0.944 | 81 | -0.011 | 0.910 | 45 |
| *UBA1819(OTU105)* | 0.022 | 0.597 | 14 | -0.004 | 0.900 | 46 |
| *K. alysoides* | 0.007 | 0.944 | 90 | 0.011 | 0.900 | 47 |
| *B. hydrogenotrophica(OTU7)* | 0.004 | 0.944 | 120 | -0.026 | 0.900 | 48 |

| | | | | | | |
|---|---|---|---|---|---|---|
| *B. hansenii(OTU11)* | -0.001 | 0.944 | 118 | 0.073 | 0.904 | 49 |
| Ruminococcaceae(OTU115) | 0.003 | 0.925 | 57 | 0.007 | 0.908 | 50 |
| *R. lactatiformans* | 0.009 | 0.944 | 101 | -0.001 | 0.912 | 51 |
| *P. secunda* | 0.004 | 0.899 | 48 | 0.004 | 0.916 | 52 |
| *C. cystitidis* | 0.001 | 0.934 | 61 | 0.004 | 0.919 | 53 |
| *S. pseudolugdunensis* | 0.000 | 0.944 | 82 | -0.013 | 0.923 | 54 |
| *Bilophila(OTU79)* | 0.004 | 0.859 | 40 | 0.003 | 0.926 | 55 |
| *R. inulinivorans* | -0.022 | 0.918 | 54 | -0.009 | 0.929 | 56 |
| Ruminococcaceae(OTU44) | 0.032 | 0.724 | 24 | -0.010 | 0.929 | 57 |
| *R. bromii* | -0.074 | 0.612 | 16 | -0.010 | 0.931 | 58 |
| *R. hungatei(OTU63)* | 0.009 | 0.937 | 63 | -0.010 | 0.932 | 59 |
| *B. cucumis* | 0.001 | 0.942 | 67 | 0.001 | 0.934 | 60 |
| *D. pneumosintes* | 0.023 | 0.944 | 104 | -0.012 | 0.935 | 61 |
| *Ruminiclostridium(OTU62)* | 0.026 | 0.257 | 4 | -0.003 | 0.936 | 62 |
| *R. hominis(OTU29)* | 0.003 | 0.944 | 109 | -0.011 | 0.937 | 63 |
| *B. magnum* | 0.001 | 0.942 | 68 | -0.011 | 0.937 | 64 |
| *I. bartlettii* | 0.001 | 0.943 | 69 | 0.001 | 0.938 | 65 |
| *P. chartae(OTU125)* | -0.030 | 0.939 | 64 | -0.002 | 0.935 | 66 |
| *H. effluvii* | 0.007 | 0.944 | 97 | 0.100 | 0.936 | 67 |
| *C. faecale* | -0.020 | 0.944 | 71 | 0.000 | 0.936 | 68 |
| *A. senegalensis* | 0.040 | 0.944 | 72 | 0.000 | 0.936 | 69 |
| *P. chartae(OTU70)* | -0.024 | 0.944 | 73 | 0.000 | 0.937 | 70 |

| | | | | | | |
|---|---|---|---|---|---|---|
| *Ezakiella(OTU53)* | 0.000 | 0.944 | 74 | -0.027 | 0.937 | 71 |
| Ruminococcaceae(OTU74) | -0.015 | 0.928 | 58 | -0.072 | 0.937 | 72 |
| *T. nexilis* | -0.023 | 0.944 | 75 | -0.017 | 0.937 | 73 |
| *L. pacaense(OTU122)* | 0.025 | 0.575 | 13 | 0.015 | 0.937 | 74 |
| *B. sp. YHC-4* | 0.009 | 0.944 | 76 | 0.014 | 0.937 | 75 |
| *B. xylanolyticus* | -0.015 | 0.831 | 36 | 0.012 | 0.937 | 76 |
| *S. variabile(OTU104)* | 0.009 | 0.920 | 125 | 0.133 | 0.937 | 77 |
| *A. ruminis* | -0.017 | 0.871 | 42 | 0.006 | 0.937 | 78 |
| *C. innocuum* | 0.005 | 0.877 | 43 | -0.044 | 0.937 | 79 |
| *V. criceti* | 0.027 | 0.923 | 56 | -0.097 | 0.937 | 80 |
| Lachnospiraceae(OTU86) | 0.004 | 0.891 | 46 | 0.007 | 0.937 | 81 |
| *S. thermophilus TH1435* | 0.045 | 0.902 | 49 | -0.035 | 0.937 | 82 |
| *R. lactaris(OTU83)* | 0.025 | 0.944 | 83 | -0.009 | 0.937 | 83 |
| *E. ramulus(OTU80)* | 0.041 | 0.141 | 2 | 0.002 | 0.937 | 84 |
| Ruminococcaceae(OTU75) | 0.039 | 0.944 | 87 | 0.012 | 0.937 | 85 |
| *O. hirsuta* | -0.016 | 0.899 | 126 | 0.003 | 0.937 | 86 |
| Lachnospiraceae(OTU68) | 0.003 | 0.912 | 52 | 0.008 | 0.937 | 87 |
| *A. equolifaciens* | -0.007 | 0.944 | 94 | 0.016 | 0.937 | 88 |

| | | | | | | |
|---|---|---|---|---|---|---|
| *B. coprophilus* | -0.016 | 0.944 | 95 | -0.023 | 0.937 | 89 |
| Christensenellaceae(OTU59) | 0.018 | 0.944 | 96 | 0.024 | 0.937 | 90 |
| *L. pacaense(OTU54)* | 0.011 | 0.789 | 32 | 0.010 | 0.937 | 91 |
| *F. plautii* | -0.030 | 0.944 | 106 | -0.085 | 0.937 | 92 |
| *E. rectale(OTU51)* | 0.052 | 0.944 | 100 | 0.112 | 0.937 | 93 |
| *A. sp. AL-1* | 0.018 | 0.944 | 102 | 0.003 | 0.937 | 94 |
| *R. gnavus(OTU47)* | 0.101 | 0.101 | 1 | 0.099 | 0.937 | 95 |
| *C. comes* | -0.040 | 0.205 | 3 | 0.039 | 0.937 | 96 |
| *R. albus* | -0.022 | 0.915 | 53 | 0.007 | 0.937 | 97 |
| *B. barnesiae(OTU40)* | -0.027 | 0.766 | 30 | 0.006 | 0.937 | 98 |
| *B. paurosaccharolyticus* | -0.041 | 0.835 | 127 | 0.005 | 0.937 | 99 |
| *B. coprosuis* | -0.033 | 0.944 | 107 | 0.001 | 0.937 | 100 |
| *S. variabile(OTU33)* | 0.012 | 0.944 | 108 | 0.170 | 0.937 | 101 |
| *E. hermanniensis* | 0.010 | 0.459 | 9 | 0.019 | 0.937 | 102 |
| *C. tanakaei* | 0.009 | 0.944 | 103 | 0.045 | 0.873 | 103 |
| *E. rectale(OTU30)* | 0.044 | 0.944 | 91 | 0.152 | 0.873 | 104 |
| *A. indistinctus(OTU26)* | 0.014 | 0.702 | 21 | 0.005 | 0.873 | 105 |
| Lachnospiraceae(OUT23) | 0.005 | 0.944 | 111 | 0.010 | 0.873 | 106 |
| *O. jeddahense* | 0.002 | 0.939 | 123 | 0.021 | 0.873 | 107 |
| *R. gauvreauii* | 0.000 | 0.941 | 66 | 0.044 | 0.873 | 108 |
| *C. europaeus* | -0.004 | 0.944 | 113 | 0.016 | 0.873 | 109 |

| | | | | | | |
|---|---|---|---|---|---|---|
| *B. ihuae(OTU16)* | -0.003 | 0.944 | 114 | 0.024 | 0.873 | 110 |
| *C. autoethanogenum* | 0.000 | 0.944 | 115 | -0.018 | 0.873 | 111 |
| *E. tayi* | 0.004 | 0.886 | 45 | 0.002 | 0.873 | 112 |
| *L. multipara(OTU6)* | -0.017 | 0.944 | 121 | 0.002 | 0.873 | 113 |
| *D. longicatena* | 0.124 | 0.944 | 93 | 0.180 | 0.873 | 114 |
| *E. sinensis* | 0.026 | 0.932 | 124 | 0.029 | 0.873 | 115 |
| *Ruminiclostridium(OTU2)* | -0.004 | 0.701 | 22 | 0.012 | 0.873 | 116 |
| *B. graminisolvens* | 0.010 | 0.944 | 122 | 0.008 | 0.873 | 117 |
| *P. nigrescens* | -0.018 | 0.944 | 116 | 0.035 | 0.873 | 118 |
| *T. glycolicus* | 0.024 | 0.944 | 99 | 0.040 | 0.866 | 119 |
| *O. valericigenes* | 0.008 | 0.944 | 78 | 0.006 | 0.859 | 120 |
| *B. barnesiae(OTU73)* | 0.000 | 0.944 | 88 | 0.012 | 0.852 | 121 |
| Lachnospiraceae(OTU42) | 0.003 | 0.909 | 51 | -0.004 | 0.844 | 122 |
| *P. soli* | -0.020 | 0.944 | 105 | -0.001 | 0.835 | 123 |
| *R. lactaris(OTU117)* | -0.008 | 0.944 | 77 | -0.054 | 0.826 | 124 |
| *Escherichia-Shigella(OTU72)* | 0.003 | 0.920 | 55 | -0.033 | 0.816 | 125 |
| *R. torques* | 0.007 | 0.846 | 38 | -0.016 | 0.721 | 126 |
| *L. hircilactis* | 0.024 | 0.935 | 62 | -0.070 | 0.683 | 127 |

Taxonomic annotation of OTUs， relative abundance, differential abundance, HITS score and intervention scores were provided and sorted by optimal cumulative intervention order. Differential abundance analysis was performed with Wilcoxon rank-sum test and adjusted with Benjamini-Hochberg method. HITS score was calculated using package NetworkX of Python (3.6.0) with permutation test for its significance. Intervention score indicated the performance of the intervention by each candidate species in dynamic intervention modeling. And the optimal combinations of the keystone species with the maximum cumulative intervention score was determined by Iterative Feature Elimination. In table, nonaf means NAFLD without advanced fibrosis and cirrhosis means NAFLD with cirrhosis.
